# Modular Fluorescent Cholesterol Naphthalimide Probes And Their Application For Cholesterol Trafficking Studies In Cells

**DOI:** 10.1101/2024.06.24.600118

**Authors:** Vicente Rubio, Nicholas McInchak, Genesis Fernandez, Dana Benavides, Diana Herrera, Catherine Jimenez, Haylee Mesa, Jonathan Meade, Qi Zhang, Maciej J. Stawikowski

## Abstract

Development of fluorescent cholesterol analogs to better understand subcellular cholesterol trafficking is of great interest for cell biology and medicine. Our approach utilizes a bifunctional 1,8-naphthalimide scaffold with a push-pull character, modified on one side with a head group and a linker on the other side connecting it to cholesterol *via* an ester bond. Through structure-function studies, we’ve explored how different substituents—linkers and head groups—affect the ability of these fluorescent cholesterol naphthalimide analogs (CNDs) to mimic natural cholesterol behavior at both molecular and cellular levels.

We categorized the resulting analogs into three groups: neutral, charged, and those featuring a hydroxyl group. Each compound was assessed for its solvatochromic behavior in organic solvents and model membranes. Extensive all-atom molecular dynamics simulations helped us examine how these analogs perform in model membranes compared to cholesterol. Additionally, we investigated the partitioning of these fluorescent probes in phase-separated giant unilamellar vesicles. We evaluated the uptake and distribution of these probes within mouse fibroblast cells and astrocytes, for their subcellular distributions in lysosomes and compared that to BODIPY-cholesterol, a well-regarded fluorescent cholesterol analog. The internalization efficiency of the fluorescent probes varies in different cell types and is affected mainly by the head groups. Our results demonstrate that the modular design significantly simplifies the creation of fluorescent cholesterol probes bearing distinct spectral, biophysical, and cellular targeting features, which makes it a valuable toolkit for the investigation of subcellular distribution and trafficking of cholesterol and its derivatives.

## Introduction

Cholesterol is a major lipid that serves as an indispensable membrane component, behaves as a cofactor for signaling molecules, and acts as a precursor for steroid hormones (1,2). Structurally, cholesterol influences membrane permeability, curvature, and lipid microdomains, in which cholesterol plays a significant role, are vital components of cellular signaling systems (3). The distribution of cholesterol between organelles within cells is facilitated by both vesicular and non-vesicular lipid transport mechanisms (2,4,5). Due to its vital role in many biological functions, the subcellular localization and trafficking of cholesterol is tightly regulated by complex pathways.

Fluorescence imaging of cellular cholesterol have been achieved by using cholesterol sensors, intrinsically fluorescent cholesterol analogs or fluorescently labeled cholesterol derivatives (6). The commonly used fluorescent cholesterol sensors are filipin (intrinsically fluorescent polyene antibiotic that binds to non-esterified cholesterol)(7) and fluorescently tagged perfringolysin O - particularly its domain 4 (D4) (8). The application of filipin has a few drawbacks including nonspecific binding to other lipids and permeabilization of cell membranes, which makes it unsuitable for live-cell applications (7). Additionally, D4-like sensors suffer from its bulky size and potentially interfere with cholesterol behavior (9). Recently, a novel cholesterol sensor named GRAM-W was developed by Koh *et al.*(10). This biosensor contains GRAM domain from GRAMD1b, however it recognizes not only cholesterol but also anionic lipids, such as phosphatidylserine (10,11).

Existing fluorescent cholesterol analogs can be divided into two categories: intrinsically fluorescent sterols or fluorophore-tagged sterols. Due to its complex interaction with various biomolecules, tagged fluorophores often impact cholesterol’s functionality (6). Consequently, different fluorescent analogs are tailored to different applications. Natural fluorescent sterols have several conjugated double bonds in the steroid ring system (e.g., dehydroergosterol, DHE and cholestatrienol, CTL). Their disadvantage is weak fluorescence near UV wavelengths, very similar to filipin and thus unfavorable for imaging (12). DHE also has very limited environmental sensitivity. Although CTL is the closest fluorescent derivative of cholesterol, its fluorescence properties and photostability are similar to, if not worse than those of DHE (13). Fluorescently labeled sterol probes have fluorophores attached to either the steroid backbone or are in the form of fluorescent cholesteryl conjugates (esters). The two most frequently used ones are nitrobenzoxadiazole (NBD) and boron dipyrromethene (BODIPY) core containing fluorophores (14–16). NBD is smaller than the bulky BODIPY group, and unlike BODIPY, NBD is an environmentally sensitive (15). On the other hand, BODIPY is a much brighter fluorophore. Modifying cholesterol’s hydrophobic tail with NBD resulted in the creation of 25-NBD-Chol and 22-NBD-Chol conjugates, which were used in many cellular studies (15,17–21). The BODIPY-Cholesterol (Bchol) was developed to better mimic cholesterol since BODIPY is electrically neutral (has no permanent dipole moment) and fluoresces more brightly than NBD. It has been widely recognized and used in many cellular assays (22,23). However, recent studies have shown that its intracellular trafficking significantly differs from that of cholesterol (23). Thus, the shortcomings of existing fluorescent cholesterol probes and biological importance of cholesterol motivated us to develop new cholesterol probes using convenient modular design. Here, we report a new class of cholesterol probes with spectral properties favorable for imaging and membrane-enhanced fluorescence. Some of them exhibit pH-sensitivity, useful for tracking cholesterol trafficking among different organelles bearing different pH. By live-cell imaging, we showed that some of the new probes are better than Bchol in terms of photoproperties and similarity to endogenous cholesterol.

## Materials and Methods

### Reagents, solvents and glassware

All reagents and solvents (including anhydrous ones) were obtained from commercial sources (Fisher Scientific) and used without purification unless otherwise noted. Synthetic, purification, and analytical steps were performed at room temperature (20 °C) unless otherwise noted. All utilized glassware was oven-dried prior to. Nile red was purchased from ThermoFisher Scientific, LysoView 650 was purchased from Biotium and BODIPY-cholesterol was purchased from Avanti Polar Lipids.

### Thin layer and column chromatography

Thin-layer chromatography (TLC) was performed on aluminum backed plates coated with silica gel (60 Å pore diameter, 200 μm layer thickness, SiliCycle or EMD Millipore). All TLC plates contained a fluorescent indicator and were visualized by UV light (λ_Ex_ = 254 nm or 366 nm) or by staining with 10% H_2_SO_4_ in EtOH solution and subsequent burning using heat gun. All column chromatography purification procedures were performed on columns packed with silica (60 Å pore diameter, 40–60 μm particle size, SiliCycle).

### NMR and mass spectrometry

NMR spectra were recorded on 400 MHz spectrometers at 22 °C in *d*-chloroform or *d*_4_-methanol. The signals in ^1^H and ^13^C NMR spectra were referenced to tetramethylsilane (TMS). The multiplicities of signals are reported as singlet (s), doublet (d), triplet (t), quartet (q), pentet (p), multiplet (m), broad (br) or a combination of these. High-resolution mass spectra (HRMS) were recorded on an Agilent 6230 Time-of-Flight (TOF) Spectrometer using Agilent 1200 series LC system (mobile phase - methanol with 0.1% formic acid) and ionized by ESI. Analysis was performed by University of Florida - Mass Spectrometry Research and Education Center through funding obtained from NIH S10 OD021758-01A1 and S10 OD030250-01A1 grants.

### Synthesis of CND probes

The synthesis scheme and methods for synthesis of CND probes are provided in the supplementary material.

### Determination of absorbance spectra in organic solvents

Stock solutions of the CND probes were diluted to 10 μM in chloroform and evaporated. The residual solids were re-dissolved in their respective solvents to obtain 10 μM solutions. The solvents used were acetone, chloroform, dichloromethane, acetonitrile, ethyl acetate, dimethyl sulfoxide, ethanol, hexanes, and methanol. The absorbance was scanned from 350 nm – 700 nm using the ThermoScientific Evolution 201 UV−VIS spectrophotometer.

### Determination of emission spectra in organic solvents

Stock solutions of the CND probes were diluted to 1 μM in chloroform and evaporated. The residual solids were re-dissolved in their respective solvents to obtain 1μM solutions. The solvents used were acetone, chloroform, dichloromethane, acetonitrile, ethyl acetate, dimethyl sulfoxide, ethanol, hexanes, and methanol. The emission spectra were scanned with a fixed excitation wavelength of 405 nm using the PerkinElmer LS 55 fluorescence spectrometer. The emission spectra were collected from 420 nm – 700 nm.

### Determination of molar extinction coefficient in DMSO and CHCl_3_

Stock solutions of the CND probes in respective solvents were diluted to 100 μM. For each data point, the solutions were diluted by a factor of 0.75 and the dilutions were performed in triplicates. The fixed absorbance values were obtained at the maxima wavelength of absorbance for each solvent. The absorbance values were obtained using the Thermo Scientific Evolution 201 UV−VIS spectrophotometer.

### Fluorescence dependence on pH in 1% OG solution

1% of octyl-β-glucoside solutions (OG) were made at pH’s ranging from 5-10 using Sorenson’s phosphate buffer. The solution was gently shaken and allowed to settle for 30 minutes. The stock solution of the CND probes was diluted to 20 μM in DMSO. For each individual pH, 95 μL of OG solution was added to a 100μL well plate. All fluorescence measurements were recorded on a 96-well plate using a Spectramax Gemini EM plate reader (Molecular Devices). To each pH OG solution, 5 μL of dye was added to each well and aspirated 3 times. The plate was then incubated in darkness for 15 minutes. With a fixed excitation wavelength of 405 nm, the fluorescence was scanned in the visible region from the top of the plate with an excitation cut off at 455 nm. The fluorescence values for each dye were obtained at the maximum wavelength of emission and plotted against pH.

### Preparation and analysis of lipid phase partitioning of CND series in giant unilamellar vesicles

Lipid stock solutions of DOPC (10 mM), brain sphingomyelin (10 mM), cholesterol (10 mM), DiD (100 μM), and CND dye (40 μM) were prepared in 9:1 CHCl_3_:MeOH solutions. To prepare the lipid mix (39.5% DOPC, 39.5% SM, 20.9% cholesterol, 0.1% DID, and 0.1% CND Dye), 34 μL of DOPC, 34 μL of SM, 18 μL of cholesterol, 10 μL of DiD, 22 μL of CND dye, were added to a single vial with a Hamilton syringe. An O-ring was placed at the center of the conductive side of the ITO glass and sealed with vacuum grease. 10 μL of the lipid mix was then added to the middle of the O-ring and air dried. The ITO glass with the dried lipid mix was placed in a desiccator under vacuum for 1 hour. For all aqueous solutions, Milli-Q grade water was filtered with a 0.22 μm pore size filter prior to use. After 1 hour, a 230 μL solution of 50 mM sucrose was placed though a 0.22 μm pore size filter onto the vacuum-dried lipid mix. The ITO glass was then placed onto the Vesicle Prep Pro (VPP) instrument with the conductive sides facing the sucrose/lipid mix solution. The automated VPP protocol was initiated which began the initiation period of heating to 60° C for 10 minutes with the frequency set to 10 Hz and the voltage set to 0 V. After the initial 10 minutes, the voltage was increased to 3 V over a period of 5 minutes. This was followed by a 3-hour period of maintaining 10 Hz and 3 V for electroformation of the vesicles to occur. After the 3-hour electroformation period a 20 minute period of decreasing the voltage to 0 V and the frequency to 5 Hz was performed. Finally, a 40-minute cooling period of the GUV solution to 23° C was performed to obtain the GUV solution. For imaging, the GUV solution was diluted in a 1:2 ratio (v:v) of GUV solution:sucrose. The imaging chamber (well) was prepared using a plastic (polypropylene) cap from HPLC injection vial that was greased and applied to a #1 glass coverslip. To that imaging chamber a 20 μL of the diluted GUV solution was added. To make the vesicles suitable for imaging, 40 μL of Milli-Q filtered water was added to the well and incubated for 15 minutes. GUV images were obtained using fluorescent laser confocal microscope Nikon A1R system using 20x objective. For CND detection, the excitation wavelength was set to 405 nm with and emission detection range 500 nm – 550 nm. For DiD, the excitation was performed at 640 nm with emission range from 663 nm – 738 nm. Individual vesicle images were cropped out one by one from the raw images. Vesicles that displayed clear phase separation as judged by the DiD channel were then processed using GUV-AP plugin to quantify L_o_ partitioning (24). The DiD channel was used for vesicle identification and stitching with a threshold coefficient of 0.12, circle extension coefficient of 1.2 and angle step of 3 degrees. A minimum of 10 vesicles were used for quantification of average L_o_ partitioning and statistical analysis.

### Molecular Dynamics Simulations

All CND probe molecules were built in the CHARM-GUI server utilizing the input generator, ligand reader and modeler modules (25–27). For the piperazine group, a positive charge was added to resemble the protonation state of piperazine at physiological pH. The CHARMM-compatible topology and parameter files of the ligands were created using CGenFF parametrization (28). The membrane structures were built utilizing the membrane builder and the probe was positioned into the membrane to have the naphthalimide moiety at the membrane surface and cholesterol embedded in the hydrophobic portion of the membrane (29). For 22NBD and 25NBD, the 3β-OH was placed at the membrane interface where the sterol and NBD fluorophore were embedded in the hydrophobic portion of the bilayer. The membrane lipids in each simulation consisted of a ratio of 1-palmitoyl-2-oleoyl-glycero-3-phosphocholine (POPC), cholesterol, or stearoyl sphingomyelin (SSM). Systems were constructed to have 0%, 5%, 25%, and 40%, moles of cholesterol with respect to the POPC lipids in the simulation. The sphingomyelin containing system, consisted of 25% sphingomyelin (SSM), 25% cholesterol, and 50% POPC. All lipids were placed symmetrically on both membrane leaflets. For all systems, physiological salt conditions were implemented with 0.15 M NaCl. For the systems that contained a positively charged piperazine an additional chlorine ion was used in the simulation and for systems containing CND10 an additional sodium ion was added to neutralize the system. All systems were simulated using CHARMM36m forcefield and GROMACS 2022.1 simulation package using GPU acceleration (30). After the systems were built, the system underwent a steepest decent energy minimization with hydrogen bond constraints using the LINCS algorithm.(31) Following the energy minimization, the system underwent six periods of equilibration as recommended by CHARMM-GUI protocol. For the equilibration periods, the temperature was coupled using the Berendsen thermostat at a temperature of 298.15 K for a time of 1 ps (32). The pressure was coupled to 1 bar using the semi isotropic Berendsen barostat for 5 ps and a compressibility of 4.5 x 10^-5^ bar^-1^. Following equilibration, the production run was carried out for a total of 100 ns. The production run utilized the NPT ensemble using the Nose-Hoover thermostat. The electrostatics were defined by using the particle mesh Ewald method, with a cut-off of 1.2 nm.

For MD trajectory analysis, the last 80 ns were used. The trajectory files from the simulations were imported onto the Visual Molecular Dynamics (VMD) software for analysis (33). For analysis of membrane properties such as tilt and thickness Membplugin was used (34). For membrane thickness, all phosphorus atoms were selected as means for determining the bilayer thickness. Cholesterol tilt angle analysis was performed by selecting the C10 and C13 carbons on cholesterol for all sterols with the mass distributions of the phosphorus atoms representing the membrane normal.

To calculate the immersion in the membrane, the oxygen (C-3 bound oxygen) atom on the sterol and where applicable the hydroxyl oxygen was used as atom selection. For determination of immersion values the position in the Z-plane for all selected atoms and phosphorus atoms were obtained. To determine the boundaries of the membrane, the Z-position of phosphorus atoms were obtained for each leaflet for every frame. The membrane center was then calculated by adding the averaged values and dividing it by 2 to obtain a Z-value for the membrane center. The distance from the center of membrane (immersion) was calculated by subtracting the position of the selected atoms from the membrane center previously calculated. The hydrogen bonds were analyzed with VMD using H-bond plugin. The number of H-bonds between the probe and membrane component (POPC, SM, water) was calculated using whole molecules as selection (e.g. POPC vs CND) with donor-acceptor cutoff distance of 3Å and the angle cutoff of 30 degrees. The bonds were analyzed across the last 80 ns of trajectory on every frame.

#### Animal Protocols

All animal protocols followed NIH guidelines and were approved by Florida Atlantic University’s Institutional Animal Care and Use Committee (IACUC).

### NIH-3T3 cell culture

NIH 3T3 fibroblasts were obtained from ATCC. One vial of cryopreserved NIH 3T3 cells were thawed in 37 °C water bath and immediately plated in 75-mL culture flasks pre-coated with Matrigel (Corning). They were grown to confluence in 3T3 culture media (DMEM + GlutaMax with 5% fetal bovine serum). Confluent 3T3 cultures were washed with PBS and detached by 1% Trypsin/EDTA. After centrifugation, cell pellets were resuspended in the same media and plated onto quad-divided glass-bottom 35 mm culture dishes at the concentration of 1,000,000 cells/cm^2^. The resulting cultures were fed with the same media as before and were used within 2-5 days.

### Mouse cortical astrocyte cell culture

Postnatal day 0-2 mouse pups were euthanized, and their cortices were dissected. 0.1% Trypsin/EDTA treatment followed by mechanical dissociation were used to separate cells. After centrifugation, cell pellets were resuspended in astrocyte culture media (DMEM + GlutaMax with 10% fetal bovine serum) and quickly plated in 75-mL culture flasks pre-coated with Matrigel. After 7-10 days *in vitro*, astrocytes outgrew other types of brain cells that were further removed by multiple washing with cold PBS and repeated shaking. Surviving astrocytes were detached by 1% Trypsin/EDTA and collected by centrifugation before cryopreservation in astrocyte media containing 9% glycerol. For experimental use, cryopreserved astrocytes were revived and plated in the same way as that of 3T3 cells except the use of astrocyte media.

### Live-cell fluorescence confocal imaging

A Nikon A1R confocal microscope equipped with a 60× (N.A. 1.40) oil objective and excitation lasers with matching optical filters for CNDs (405 nm excitation) or Bchol (488 nm excitation), 496 nm long-pass dichroic filter, and 510/20 nm band-pass emission filter), Nile Red (540 nm excitation, 565 nm long-pass dichroic filter, and 600/40 nm band-pass emission filter), and LysoView 650 (640 nm excitation, 680 nm long-pass dichroic filter, and 710 nm long-pass emission filter). Nikon Elements program was used for image acquisition. Laser power, gain, and offset were chosen to maximize fluorescence dynamic range for every channel and minimize bleed-through of other fluorophores. And those settings were kept the same for the same types of imaging experiments. Cells were loaded with desired probe to obtain a final dye concentration of 1 μM for 1 hour and then incubated in dye-free media for desired periods of time until imaging. Before imaging, cells were loaded with Nile Red and LysoView (0.5 μM) for 1 hour and then washed with dye-free media immediately before placing inside an Oko-Lab weather chamber which was mounted on the microscope stage. The chamber was pre-warmed to 37 °C and filled with 5% CO_2_ and 100% humidity. The 60× objective was also pre-warmed to 37 °C. For every sample, three fields of view were selected randomly and images with a frame size of 1024 x 1024 pixels were acquired. For all z-stack acquisition, the z-axis step size was 0.3 mm. For all time-lapse imaging, the frame interval was 10 seconds, and the duration was 5 minutes. Acquired images were then further analyzed using ImageJ/Fiji (35).

### Particle analysis

For particle analysis, the fluorescence intensity across Z-stack was analyzed and single frame was extracted and saved for further analysis. To generate ROIs for particle analysis, a trainable Weka segmentation was employed (36). The Weka classifier was trained on a series of three noisy (low s/n) and three good (high s/n) from the pool of acquired images. Classified images were converted into masks and ROIs were generated. Upon ROI generation, particle analysis was performed in Fiji/ImageJ with object detection between 0.1 µm^2^ -10 µm^2^. For each ROI, mean pixel intensity and area were obtained. For obtaining the diameter, the area was used, and the particles assumed to be perfect circles. Particle size used for analysis was in the range between 0 – 3 µm. Data was visualized and analyzed using R programming language and RStudio package. For statistical significance testing, Wilcoxon test was employed.

### Colocalization Analysis

In colocalization studies, while Pearson’s correlation coefficient (PCC) is useful for assessing the overall association between two probes in an image and is being reported frequently, it falls short in accurately measuring the fraction of one marker that colocalizes with another, a crucial aspect in cell biology analyses (37). In contrast, the Manders’ colocalization coefficient (MCC) directly measures co-occurrence regardless of signal proportionality. However, to employ MCC effectively, it’s imperative to eliminate background noise from pixel intensities (37). While subtracting a global threshold value is a common method, determining the appropriate threshold can be challenging. Costes *et al.* devised an efficient, robust, and reproducible approach to automatically identify the threshold value by analyzing the pixel values that yield a positive Pearson’s correlation coefficient (PCC) (38). It iteratively measured PCC for different pixel intensity values until reaching values where PCC dropped to or below zero. The corresponding intensity values on the regression line were then utilized as thresholds for each channel. Only pixels surpassing both thresholds were considered colocalized. Consequently, the Costes’ method eliminates human bias. For these reasons, we used it and effectively identified labeled structures from background noise, facilitating accurate Manders’ colocalization coefficient (MCC) in various scenarios. As it is implemented in EzColocalization plugin available for ImageJ (39), we used EzColocalization to determine both, PPC and MCC.

Colocalization was performed between selected channels (e.g. CND vs. LysoView) within each Z-stack on a plane-by-plane basis in a batch mode, with Costes’ thresholding, using EzColocalization plugin in Fiji/ImageJ. PCC and MCC values were obtained from each slice in Z-stack. Data from 1h and 24h was then compared plotted using GraphPad Prism (ver. 10). Statistical analysis was performed using Wilcoxon matched pairs signed rank test.

## RESULTS

### Preparation of CND probes

All fluorescent cholesterol CND analogs can be prepared by reaction of amino acid (linker) with 4-bromo-1,8-naphthalic anhydride to form 4-bromo-1,8-naphthalimide which then is used to esterify cholesterol (Scheme S1). Substitution of bromine at C4 position renders complete CND probe with unique photophysical properties. The order of reaction sequence depends on linker used. As linkers, we have employed glycine, L-serine and β-alanine. Ethanolamine, piperazine, morpholine, imidazole, hydroxyl, 4-hydroxypiperidine and 4-carboxypiperidine were used as head groups in various combinations to make a set of ten different CND probes (**Fig.** 1). The choice of linkers and head groups was supported by the molecular dynamics (MD) simulation analyzing the behavior of cholesterol and its interactions in the membrane. The chemical synthesis of CND analogs was accomplished with moderate yields, in three or four steps, depending on the type of the linker used.

**Fig. 1.**
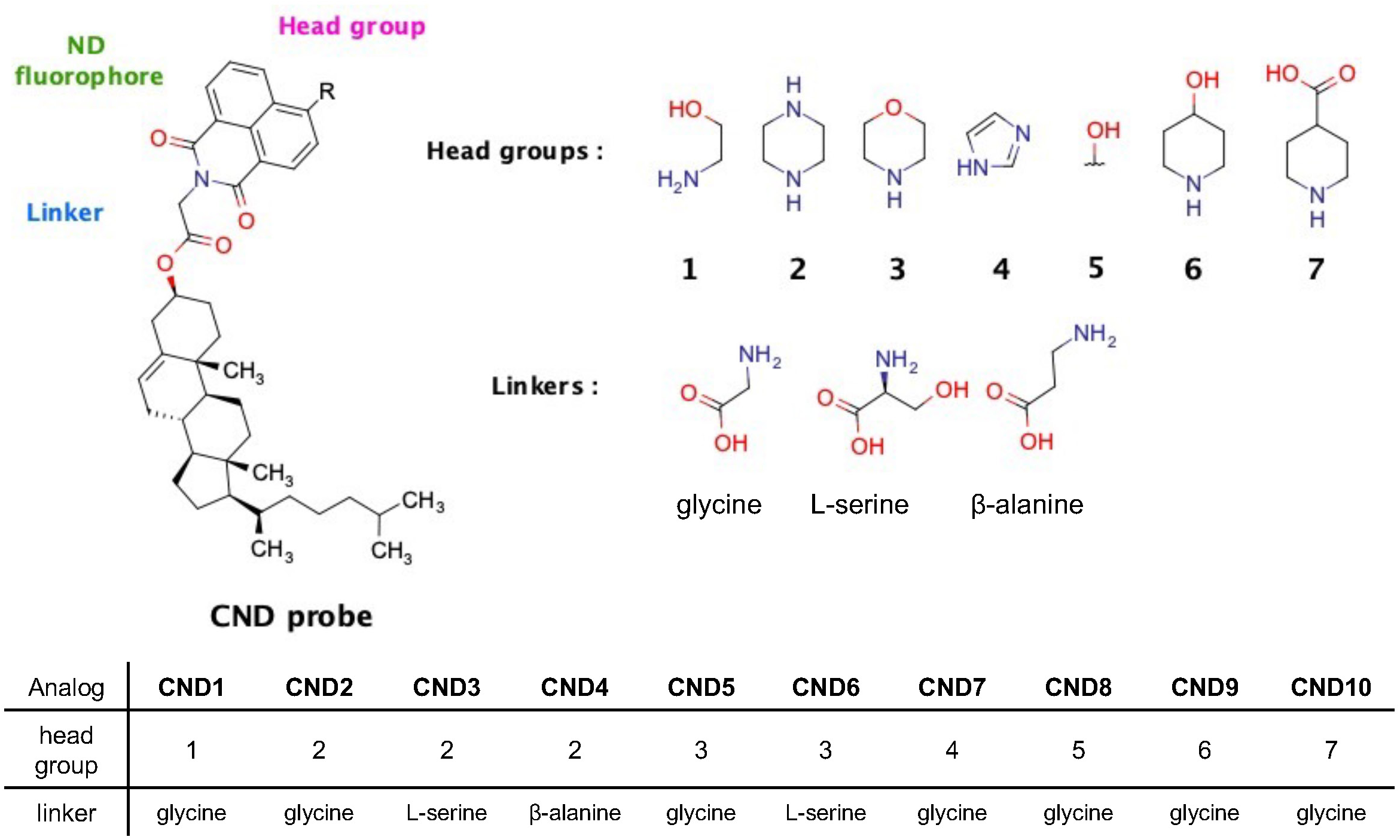
Structural modularity of CND probes. Variation of head groups and linkers used to make CND1-CND10.

### The influence of the head groups and the linkers on CNDs’ photophysical properties

To establish the solvatochromic properties of CNDs, we determined the excitation and emission spectra of the CNDs in various solvents (**Fig.** S1 and Table S1). The absorbance maxima for the analogs with nitrogen substituted aliphatic rings at C4 position (CND2-6, 9, and 10) are proximal to 405 nm which is a common laser line for excitation in fluorescent microscopy. Most of their emission spectra also nicely fall into the commonly used green channel (i.e., 500-550 nm). The substitution at C4 with aliphatic ethanolamine results in obtaining analog (CND1) having red-shifted λ_max_ ∼ 420-440 nm. The solvatochromic properties of the analogs can be illustrated by the Lippert-Mataga plot (**Fig.** S2). To quantify solvatochromic properties, the maximum wavelengths of absorption and emission (λ_abs_/λ_em_) were obtained, and the Stokes’ shifts (Δω) were plotted as a function of solvent orientation polarizability (Δf) to obtain Lippert-Mataga (LM) plot (**Fig.** S3)

(40). The deviation of positive, linear correlations suggests that there are fluorophore-fluorophore or fluorophore-solvent interactions influencing the ICT process in the ground and excited states. CND7 is the only analog that does not exhibit detectable correlation, corresponding to undesirable spectral profile. With its absorbance maxima ranging from 342 nm – 346 nm and emission maxima ranging from 440 nm – 505 nm, CND7 was found to be unsuitable for further microscopy studies as a probe. CND8 is the only probe with hydroxyl group present at the C4 position. It also exhibits unique solvatochromic properties due to its hydroxyl (phenolic) group interacts strongly with polar protic solvents (MeOH and EtOH) through hydrogen bonds (**Fig.** S1). For all CNDs the molar extinction coefficients were determined in DMSO and chloroform (Table S2) ranging from 8000-16000 M^-1^ cm^-1^, depending on the analog and solvent type used. This is in agreement with prior studies on substituted naphthalimides(41,42).

The solvatochromic characteristics of the ND scaffold makes CNDs sensitive to membrane environments (43). To test that, we have obtained fluorescence profiles of CNDs at 1 μM in 1% of octyl glucoside micellar solution (i.e., mimicking membrane environment). In comparison to their spectra in water we have found a head group dependent change (**Fig.** S3). Probes containing ethanolamine (CND1), piperazine, (CND2, CND3, CND4) and hydroxyl group (CND8) have substantial increase in their fluorescence (50 – 400-fold) in the micellar solution. Interestingly, for analogs containing morpholine group (CND5 and 6) there is a strong fluorescence signal in micellar solutions but also a considerable fluorescence in water. Upon close inspection by transmission light, a suspension of particles was observed in the aqueous CND5 and CND6 solutions. The highly fluorescent particles were also observed using fluorescent microscope. We postulate that their considerable fluorescence in water likely results from the phenomenon known as aggregation induced emission (AIE) (44). In fact, AIE has been well documented in some naphthalimide-based probes (45).

The protonation state of the piperazine head group also influences the fluorescence of the ND scaffold. We have successfully used this molecular property to design a membrane and pH-sensitive lipid probe, ND6 (43). Incorporation of piperazine residue in CND2-CND4 analogs resulted in pH sensitivity, which was tested in 1% OG micellar solutions containing Sorensen’s phosphate buffer with variable pH (**Fig.** 2). The protonation of the amino group of piperazine contributes to the inhibition of the photoinduced electron transfer process in the naphthalimide scaffold (46,47), which in turn produces higher (∼2-3 fold) fluorescence at pH 5 than at pH 7.

**Fig. 2.**
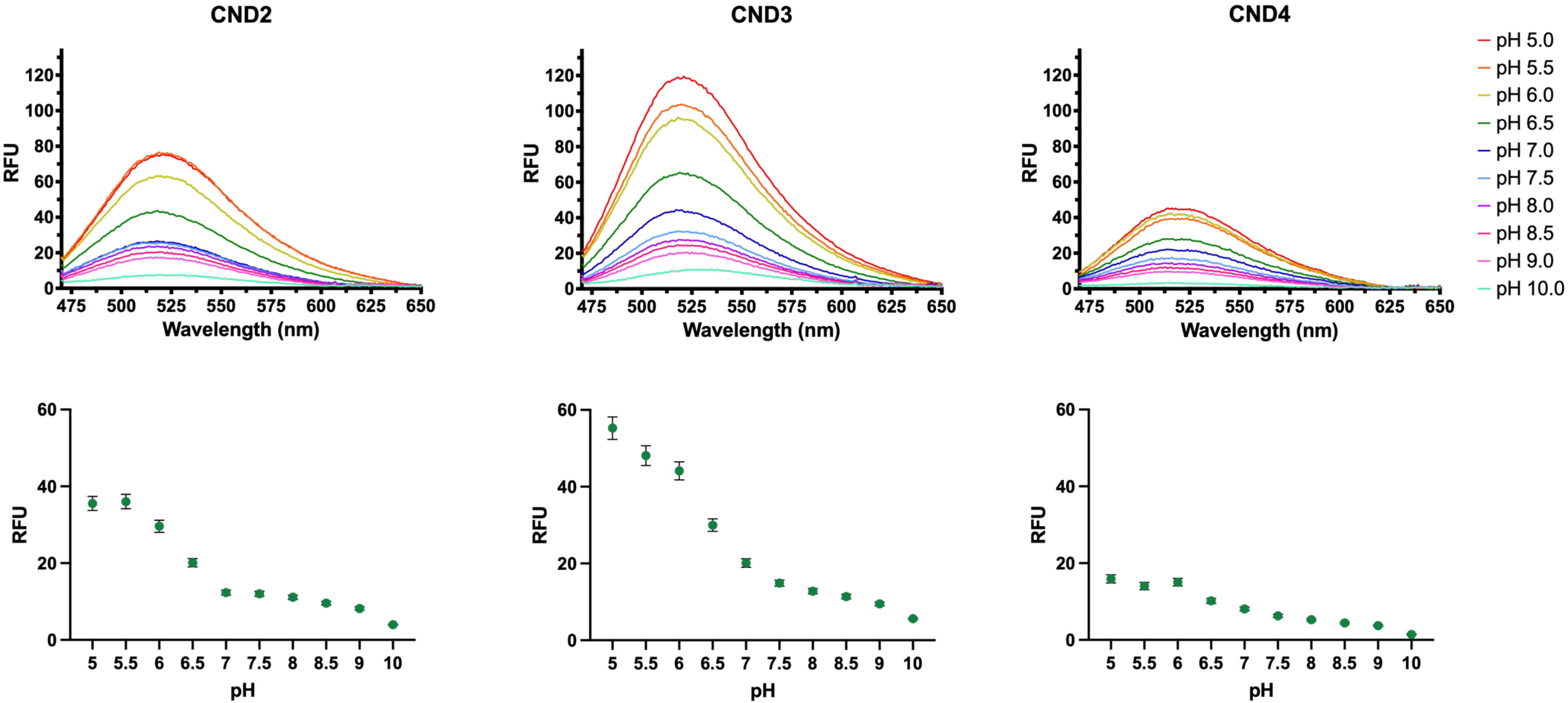
Emission spectra of CND2-CND4 in 1% octyl glucoside micellar solution in Sorensen’s buffer at various pH (._Ex_=405 nm) demonstrating pH sensitivity of these analogs (top row). The fluorescence in the 470 -650 nm range was used to calculate area under the curve to determine the relative fluorescence change (mean ± SEM, bottom row). The fluorescence increases 2-3 fold with pH drop from 7 to 5.

### Lipid partitioning of CND probes

To determine the partitioning preference of CND analogs to L_o_/L_d_ phases, we have employed electroformed GUVs from a mixture of DOPC:SM:Chol (2:2:1) containing 0.1 mol% of L_d_ marker (DiD) (48) and the same amount of individual CNDs (49). Results of lipid partitioning in GUVs is shown in **Fig.** 3A. Notably, all CND analogs demonstrate at least 30% partitioning into the L_o_ phase. The standout performer among these analogs is CND3 with ∼50% partitioning, which is comparable to Bchol (49). These results demonstrate that the introduction of a hydroxyl group into the linker structure significantly enhances the lipid partitioning performance, as evidenced by the improved performance of CND3 compared to its counterpart lacking the hydroxyl group (CND2). This observation is further validated by the noteworthy performance of CND8, which includes an additional hydroxyl group, securing its position as the second-best performing analog. As we predicted, analogs with an -OH group present in the head group (CND1, CND8, CND9) exhibited above-average performance. On the other hand, the results show that the character of the head group also plays significant role in the partitioning process. The neutral head group analogs (morpholine, CND5 and CND6) exhibited the weakest partitioning into L_o_ phase compared with all other analogs.

**Fig. 3.**
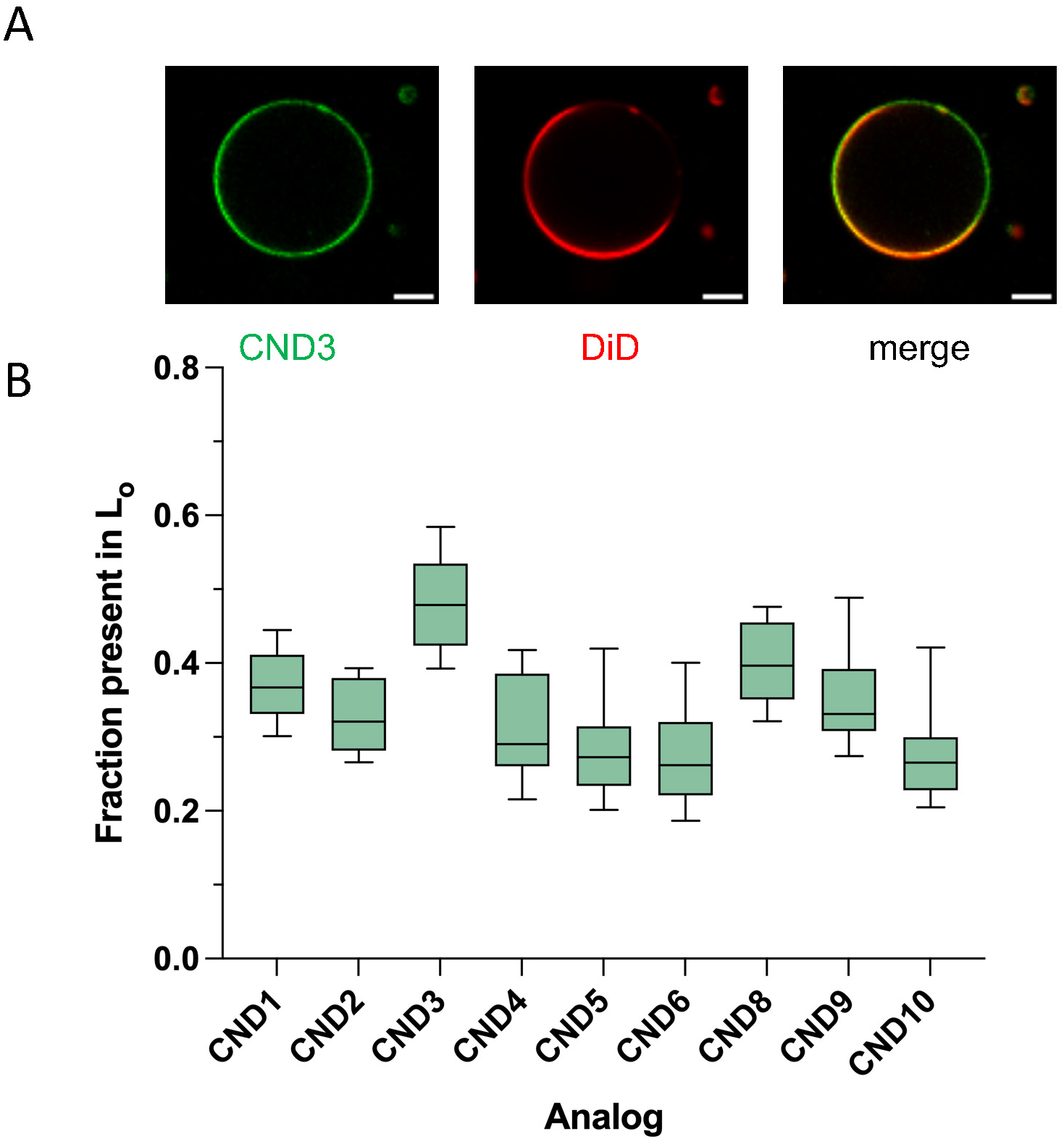
Partitioning of CNDs between Ld/Lo phases in phase-separated GUVs. **A**, representative fluorescent confocal image of CND3 and Ld marker DiD. To determine the partitioning the fluorescence intensity around vesicle perimeter was measured using GUV-AP plugin for ImageJ (see supplementary info). Scale bar, 10 μm. **B**, Fraction of probe partitioning into L_o_ phase for all tested CNDs.

### Molecular dynamics simulations of CNDs in lipid bilayer

To evaluate the molecular interactions between CNDs and lipid membrane, we have conducted a set of molecular dynamics (MD) simulations. In this experiment we also, we included the 22-NBD-Chol, 25-NBD-Chol and 3-Hex-NBD-chol. Five different membrane model systems were tested (Table S3). Four of them had cholesterol and POPC in various amounts. The last one was composed of SM:Chol:POPC (1:1:2), for better mimicking the plasma membrane and L_o_ phase (50,51). Since 25% Chol is commonly found in the plasma membrane, two of the five systems, namely Chol:POPC at 1:3 and SM:Chol:POPC at 1:1:2, were evaluated in detail for all CND analogs as well as NBD analogs.

To efficiently evaluate cholesterol behavior, we focused on two key parameters, i.e., cholesterol’s tilt angle and the immersion depth. The tilt angle reflects how CNDs incorporate into the lipid bilayer and interact with surrounding lipids (52). We selected a vector defined by the positions of C10 and C13 atoms within the cholesterol moiety to determine the cholesterol tilt angle (θ) relative to the membrane normal (Z). The outcomes of tilt angle analysis are shown on **Fig.** 4 (Table S3). Clearly, there was a reduction in the cholesterol tilt angle when cholesterol content increased, particularly in the presence of saturated lipids. This agrees very well with previously published data (52). As cholesterol content influences membrane thickness and organization, it affects the behavior of membrane-embedded probes. With decreased cholesterol tilt angle, lipid packing was increased as well (Table S4). Between 0 and 40% cholesterol, the change is apparent. Also, the addition of SM (saturated lipids) not only increased the membrane thickness but also enhanced lipid condensation which is represented by the diminished cholesterol tilt angle (53).

**Fig. 4.**
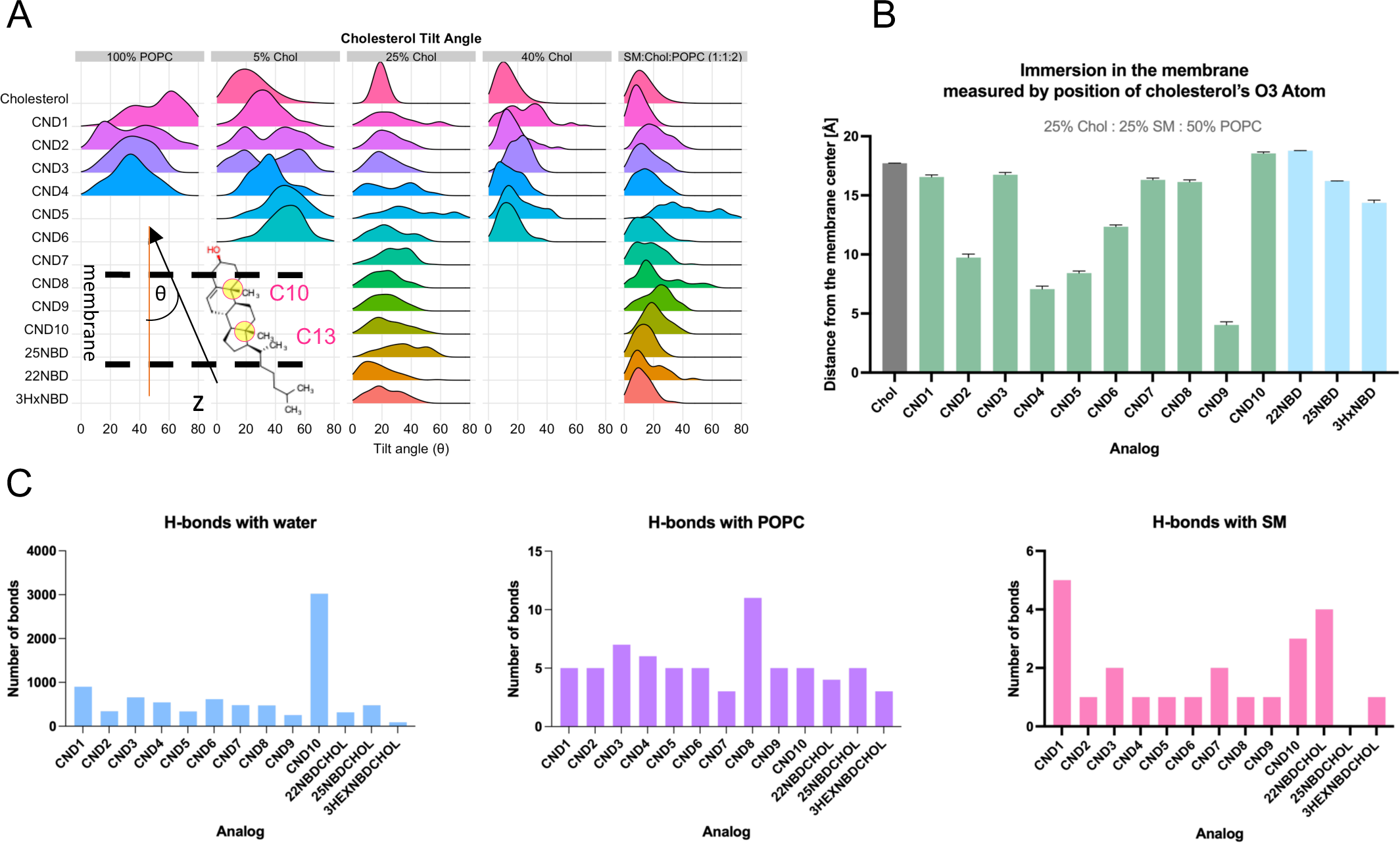
Characterization of probes using molecular dynamics simulations. **A**, analysis of the tilt angle of cholesterol. C10-C13 atoms were selected as selection (inset). **B**, analysis of the depth of probe immersion in the membrane in SM:Chol:POPC system. **C**, analysis of hydrogen bond patterns between probes and surrounding water, POPC and sphingomyelin (SM).

To measure the depth of immersion, the position of cholesterol O3 atom in the Z-plane was used and its distance from the middle point of the membrane bilayer was calculated in both Chol:POPC, 1:3 (25% Chol) and the SM:Chol:POPC, 1:1:2 systems (**Fig.** S4, left column). The immersion depth of other hydroxyl groups within certain CND probes, either in linkers (CND3, CND6) or head groups (CND1, CND8, CND9) was also calculated (**Fig.** S4, right column). Since the presence of SM lipids in the system influences membrane behavior (ordering) and increases the type of in-membrane interactions available for the probes (51), the SM:Chol:POPC system revealed better probe differentiation than the simpler Chol:POPC (1:3) system (comparing the top vs. the bottom plots in **Fig.** S4). In the SM:Chol:POPC system the CND1, CND3 and CND8 were immersed in the membrane very similarly to cholesterol. CND10 is immersed the least in the membrane, higher than cholesterol (**Fig.** 4B), which is likely due to the electrostatic interactions of carboxylate group with choline moiety from SM and POPC as well as the formation of hydrogen bonds with water (**Fig.** 4C). It seems that 4-carboxypiperidine head group position carboxylate at the right height in respect to choline group to maximize these interactions. In terms of hydrogen bond formation with SM, CND1 performed the best (**Fig.** 4C). CND1 ethanolamine head group is water exposed and thus can form multiple hydrogen-bonds with the phosphate groups from phospholipids as well as water, which prevents CND1 from further immersion into the lipid bilayer. The interactions of CND8 and its hydroxyl head group also follows similar pattern, but the effect is less pronounced (**Fig.** 4C) On the contrary, the hydroxyl group of CND9 (4-hydroxypiperadine head group) forms the least number of hydrogen bonds with water due to the very hydrophobic character of this head group (**Fig.** 4C). It is also the reason that CND9 exhibits the deepest immersion among all analogs tested (**Fig.** 4B). The analogs containing piperazine head groups (i.e., CND2, 3 and 4) also show electrostatic interactions with the phosphate groups of phospholipids, although their immersion depth depends on their linkers (Movie S1). CND4 with β-alanine linker shows the second deepest immersion in the membrane (**Fig.** 4), as this linker enables for great flexibility and stretchability between naphthalimide moiety and the cholesterol residue. CND2 with its glycine linker is slightly shorter and more constrained than the beta-alanine one (**Fig.** 4B). CND3 serine linker’s side chain hydroxyl group has significant effect on immersion in the membrane, which allows for additional hydrogen bond interactions with surrounding lipids and water, imitating the behavior of native cholesterol’s hydroxyl group (**Fig.** 4B). The serine linker effect is also evident in case of CND6 when compared to the CND5 analog which share the same (morpholine) head group. Interestingly, the morpholine head group also significantly changes the probes’ behavior compared to piperazine moiety. Together, the results from simulated SM:Chol:POPC membrane system are in qualitative agreement with experimental data on analogs partitioning in GUVs where CND3, CND8 and CND1 showed the most L_o_ partitioning (**Fig.** 3).

### Live-cell imaging of CNDs

We studied the behavior of CNDs *in vitro* using cultured 3T3 fibroblast cells and mouse astrocytes. Bchol was used as reference. In addition, we used fluorescent markers for lipid droplets (Nile red)(54) and lysosomes (LysoView™ 650), spectrally separable from CNDs. Following 1-hour incubation with individual CNDs, cells were rinsed with dye-free media to remove unloaded dyes. Multi-color fluorescence confocal imaging was carried out immediately after the 1-hour incubation and again at 24 hours post-incubation to assess the localization of the compounds. Not surprisingly, different CNDs exhibit different cellular staining patterns. Some were weakly loaded (e.g., CND5) while others were strongly present in cells (e.g., CND1, CND2-4); some were more diffused (e.g., CND1) while others were more punctuated (e.g., CND2-4); some accumulated all over the cells (e.g., CND6, CND9,) while others were more clustered near nuclei (e.g., CND3) as shown in **Fig.** S5.

To generate masks for fluorescent puncta quantification, we used machine learning approach, implemented in Weka trainable segmentation (**Fig.** S7) (55). We quantified the accumulation of probes by counting the fluorescent puncta per cell across both cell types as well as both time points (**Fig.** 5C). Notable patterns emerged. In astrocytes, CNDs with piperazine head groups (CND2, CND3, and CND4) showed a larger number of fluorescent puncta in cells at both 1 and 24 hours (**Fig.** 5C, S7), indicating that the head group may be the primary factor influencing their uptake and accumulation. In 3T3 cells, a similar pattern was observed, although Bchol and CND6 were taken up more efficiently in 3T3 cells than in astrocytes.

**Fig 5.**
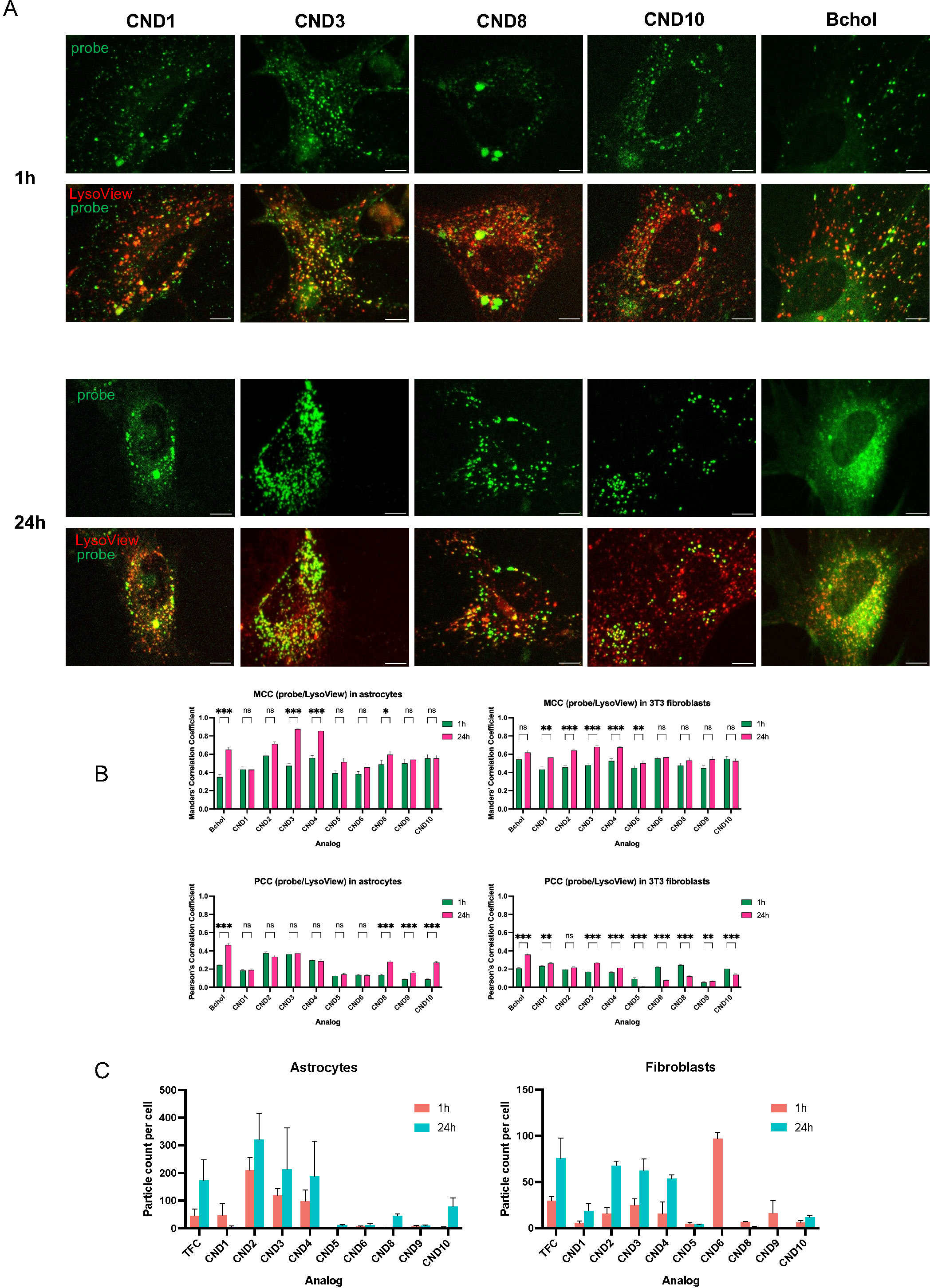
Live cell imaging of CND probes in cultured astrocytes. **A**, sample images for analogs that are neutral (CND1, 8) positively charged (CND3) and negatively charged (CND10) compared with Bchol. Overlay images show co-labelled LysoView (red) overlapping with selected CNDs better than Bchol. Scale bar, 10 μm. **B**, Manders’ and Pearson’s correlation coefficients between probe/LysoView at different time points and cell types. **C**, relative probe uptake and accumulation measured by particle count at different time points in astrocytes and 3T3 fibroblasts.

The role of the linker in probe uptake is evident when examining analogs with morpholine head groups. For both cell lines, CND5, which has a glycine linker, exhibited the least uptake (**Fig.** 5C and S6, S7), likely due to its poor solubility and propensity to aggregate in water. On the other hand, CND6, featuring a serine linker, showed significantly more uptake and intracellular accumulation (**Fig.** 5C and S6, S7). Interestingly, in 3T3 cells, CND6 displayed a high number of fluorescent puncta at 1 hour but decreased dramatically at 24 hours (**Fig.** 5 and S7). This suggests higher efflux or degradation of CND6 by fibroblast cells. Similar but less pronounced patterns were observed in astrocytes.

Neutral CND analogs with hydroxyl-containing head groups (CND1, 8, and 9) generally had lower number of fluorescent puncta compared to those with piperazine head groups, CND2-CND4 (**Fig.** S7). However, CND1, CND8 and CND9 shared a similar uptake behavior as CND6 in 3T3 cells. In astrocytes, the patterns were the opposite, with CND8 and 9 showing more accumulation and CND1 exhibiting less. CND10, with a negatively charged carboxylate head group, exhibited a comparable distribution pattern in both cell types, accumulation after 24h, but the degree of that accumulation was more pronounced in astrocytes.

Analyzing cell-wide fluorescence signal distribution in astrocytes reveals distinct results (**Fig.** S5 and S7). For CND2-4, the fluorescent puncta were scattered across the cell at 1 hour. By the 24-hour mark, puncta appeared to be more perinuclear. In contrast, CND8, 9, and 10 exhibited a different pattern where the puncta not only became more defined after 24 hours but also showed a reduced fluorescence throughout the cells. The CND1 presented yet another distinct pattern, i.e., puncta were prominent at 1 hour and became more widespread by 24 hours. Bchol, also accumulated as fluorescent puncta; was more dispersed, and remained prominently diffused even 24 hours after loading (**Fig.** S5).

Next, we measured the size of fluorescent puncta. While most of them were approximately 1 μm in diameter, there was a significant variation in size as depicted in **Fig.** S8. In 3T3 cells, the puncta sizes were generally large for all CNDs except those with free hydroxyl head groups (CND6, 8, and 9). In astrocytes, however, the largest increase in puncta size was seen with CNDs that have piperazine head groups (CND2-4). This size increase was also noted for hydroxyl-containing head group analogs (CND8 and 9) and the negatively charged CND10. We further compared the accumulation of probes within the fluorescent puncta by assessing the average fluorescence intensity within individual particles (**Fig.** S9). As reference, Bchol exhibited high average intensity in both cell types, slightly more in 3T3 cells. The fluorescent puncta of CNDs with positively charged head groups (CND2-4) exhibited a strikingly higher average intensity.

To determine the nature of those fluorescence puncta, we performed co-labeling with LysoView, a commercially available marker selective for lysosomes and Nile red, typically used for lipid droplet visualization (54). By comparing co-staining between 1-hr and 24-hour time points, Bchol shows accumulation in lysosomes and lipid droplets that increases over time in both cell types (**Fig.** 5). This observation is in with existing literature reports in which Bchol was shown to partition to lipid droplets in CHO and HeLa cells, and this accumulation was attributed to the BODIPY group (6,23,56). There is also report suggesting that Bchol accumulated in lysosomes after prolonged incubation (57). In terms of the CNDs, we observed co-localization with both LysoView and Nile red that increased over time, just like Bchol, although the degrees of co-localizations at different time points vary among different CNDs (**Fig.** 5). While we cannot absolutely exclude the possibility of nonspecific labeling, particularly by Nile red (58), it is very possible that CNDs, just like Bchol enter both lysosomes and lipid droplets. It is also possible that some lipid droplets and lysosomes are too close to be distinguished by the fluorescence imaging.

## Discussion

Our recently published study has demonstrated that 1,8-naphthalimide (ND) scaffold possesses excellent solvatochromic properties and can be used to design membrane-sensitive probes (43). The ND is a push-pull type of fluorophore, exhibiting upon excitation an intramolecular charge transfer fluorescence mechanism. This scaffold is highly sensitive to its solvent environment (59,60). Naphthalimide-conjugated molecules have been used as DNA intercalating agents, RNA sensors, cancer therapeutics, organogels, organic light emitting diodes, enzyme sensors, and cell imaging agents (61–65). We have taken the advantage of the unique structure and fluorescent properties of 1,8-naphthalimide scaffold (42) that provides opportunity for easy chemical modifications to create a series of cholesterol-tagged probes (**Fig.** 1). Large Stokes’ shifts (∼120 nm) and solvatochromic properties of ND scaffold make it very useful for obtaining membrane sensitive probes (42,43). The modular structure of these cholesterol probes allows us to build a toolbox of cholesterol probes with different membrane- and pH-sensitivities. These probes are uniquely capable of mimicking cholesterol or its derivatives, advantageous for studying cholesterol trafficking and distribution in live cells. Recognizing the crucial role of the hydroxyl group in cholesterol’s interactions with different lipids, including sphingomyelin (SM) and proteins, inspired us to incorporate an additional hydroxyl group into the design of the CND probes (into linker or the head group).

We categorized resulting analogs into three groups: neutral (CND5, CND6, CND1, CND8, CND9), and charged (CND2, CND3, CND4, CND10). Each compound was assessed for its solvatochromic behavior in organic solvents and model membranes. The unique photophysical properties of CND probes make them more useful in cell imaging than NBD- and BODIPY-conjugated cholesterol probes. CNDs have much larger Stokes shifts (∼130 nm) than NBD-conjugated probes (∼40 nm) and Bchol (∼10 nm), which is particularly beneficial for co-imaging with other fluorescent probes. The solvatochromic characteristics of the ND scaffold makes CNDs sensitive to membrane environments (43). In contrast, some of the neutral analogs (CND5, CND6, CND9) exhibit aggregation in water due to AIE effect. The observed contrast in aggregation propensities between CND2-CND5 and CND3-CND6 pairs, despite sharing identical linkers but possessing different head groups, is evident. This highlights the substantial influence of the head group on the aggregation (and solubility) of these probes. More specifically, the charged state of the piperazine group under physiological pH induces charge repulsion among fluorophore groups, mitigating aggregation in aqueous solutions. The protonation state of the piperazine head group also influences the fluorescence of the ND scaffold. The exposure of the piperazine (and the naphthalimide core) to aqueous environment is proportional to the fluorescence change and correlates with the type and length of the linker, which was also confirmed by the MD simulations. The pH-sensitivity of CND2-CND4 has great potential in cell biology, since organelles in the endosomal pathway exhibit different pH (e.g., PM is ∼pH 7, endosomes ∼pH 6, and lysosomes ∼pH 4.5) (66). Hence, those CNDs are useful to measure intracellular pH changes as well as to track cholesterol trafficking, similarly to previously reported ND6 probe (67). Moreover, such pH-sensitive cholesterol probes or lipid probes in general are very useful in imaging synaptic vesicles at nerve terminals (68) and endosomal pathway in various cell types, bearing great promise in studying cholesterol-related disorders like Alzheimer’s disease (68–70).

The L_o_/L_d_ partitioning preference of CND analogs were evaluated in phase-resolved GUVs in tertiary lipid mixture of DOPC:SM:Chol (2:2:1). Comparatively to other lipid partitioning studies, with similar conditions, the CND dyes clearly outperform other probes, such as 22- and 25-NBD-Chol (having free hydroxyl group) (49). In the cited study, the 22-NBD-Chol, and 25-NBD-Chol partitioned to L_o_ phase in ∼5% and ∼20% respectively. Interestingly, the ester analog 3-C6-NBD-Chol (abbreviated as 3HxNBD in this work) performed much better, with ∼30% for partitioning coefficient. It was also noted that with the additional extension of aliphatic linker, the partition coefficient dropped significantly to ∼7% (3-C6-NBD-Chol vs. 3-C12-NBD-Chol). These results suggest that further modification of L_o_ partitioning can be achieved by altering the attachment positions for fluorophore and linker.

Our best analog, CND3 with its L_o_ partitioning ∼50%, is comparable to Bchol (49). Our results demonstrate that the introduction of a hydroxyl group into the linker structure significantly enhances the lipid partitioning performance, as evidenced by the improved performance of CND3 compared to its counterpart lacking the hydroxyl group (CND2). Among CNDs with positively charged head group (i.e., CND2-4), the L_o_ partitioning is proportional to their immersion in the membrane and correlated to their pH-sensitivity (**Fig.** 3B). In contrast, the negatively charged head group of CND10 weakens its L_o_ partitioning. Our findings demonstrate that the combination of linkers and head groups collectively determines the resemblance of CNDs to cholesterol in the lipid bilayers.

The behavior of CND probes in model membranes were studied using molecular dynamics simulations. The extensive all-atom molecular dynamics simulations helped us examine how these analogs perform in model membranes compared to cholesterol. Analogs were evaluated based on cholesterol tilt angle, and the depth of immersion in the membrane. As anticipated, these parameters were dependent on the type of the linker and head group used. More flexible linker (β-alanine vs. glycine) enables for deeper immersion in the membrane and results in smaller cholesterol tilt angle. The presence of the hydroxyl group in the linker (serine) constrains the flexibility of the probe and the immersion depth is determined by the nature of the head group and its interactions with phospholipids. Ethanolamine, piperazine and 4-carboxypiperidine bring the probe towards the phospholipid-water layer through interactions with phosphate groups and water, while neutral head group (morpholine) prefer to be in presence of hydrocarbon environment. The hydroxyl group containing analogs – CND8 and CND9 differ dramatically in the depth of immersion. Hydroxyl group at position C4 in CND8 analog, due to its phenolic character is much more polar (and acidic) than the hydroxyl group present in 4-hydroxypiperidine group in CND9. Thus, CND8 is embedded in the membrane similarly to CND1 and CND3 while CND9 drops in. The immersion depth and cholesterol tilt angle of NBD containing cholesterol analogs (22NBD, 25NBD and 3HxNBD) was comparable with that of CND1, CND3 and CND8 and the ester containing 3HxNBD had the deepest immersion in the membrane. The cholesterol tilt angle in 3HxNBD was smaller than in 25NBD and on par with that of 22NBD. Our results are different for previously reported MD data for these analogs (71). In the cited study NBD probes were positioned rather sideways in relation to membrane normal. The difference between these results can be attributed to the difference in the type of forcefield used and perhaps initial orientation of the probes in the membrane. As mentioned in the recent review, the data on the behavior of NBD containing cholesterol probes in the membranes is rather unclear (72).

In our MD simulations we have tested five different membrane models. The presence of SM lipids in the system influences membrane behavior (ordering) and increases the type of in-membrane interactions available for the probes (51). We found that the SM:Chol:POPC system showed better probe differentiation than the simpler Chol:POPC (1:3) system (comparing the top vs. the bottom plots in **Fig.** S4). This system also better reflected the experimental data for probe partitioning into L_o_ phase.

We have tested CND probes *in vitro* using cultured 3T3 fibroblast cells and mouse astrocytes and perform their imaging in live cells. We evaluated the uptake and subcellular distribution of these probes and compared that to BODIPY-cholesterol (Bchol), a well-regarded fluorescent cholesterol analog. Results revealed the differences between CND analogs and their staining patterns. The differences were also found between the types of cells used. The distinct staining patterns suggest that the nature of the head group may be the primary factor determining the uptake and probe accumulation. This was evident for charged analogs (CND2, CND3, CND4) compared to neutral analogs (e.g. CND1, CND6, CND8 and CND9). We were surprised to see CND6 clearance in fibroblasts after 24h. This can be attributed to probe efflux (for example by ABC transporters (73)) or degradation of CND6. Similar, but less pronounced patterns were observed in astrocytes.

Some CNDs (i.e., CND1, CND2-4, CND6, CND8), by better functional resemblance, elucidate intracellular cholesterol trafficking through endosomal pathway(74), possibly *via* interactions with caveolae. These flask-shaped lipid microdomain structures are involved in cholesterol regulation, signaling, proliferation, migration, and endocytosis (75,76). In addition, those CNDs may also be useful to illustrate cholesterol exchange between lipid droplets and endosomes/lysosomes through vesicular and non-vesicular pathways. Taking into account the central role of lysosomes and their role in subcellular cholesterol distribution (77) we have focused on CNDs colocalization analysis with this organelle. We observed co-localization of CNDs with both LysoView and Nile red that increased over time, just like Bchol, although the degrees of co-localizations at different time points vary among different CNDs. While we cannot absolutely exclude the possibility of nonspecific labeling, particularly by Nile red (58), it is very possible that CNDs, just like Bchol, enter both lysosomes and lipid droplets. It is also possible that some lipid droplets and lysosomes are too close to be distinguished by the fluorescence imaging. For example, lipid droplets laden with CNDs (or containing cholesterol) could be processed by autophagy, whereby autophagosomes—double-membraned vesicles—engulf and enclose materials for degradation and recycling in lysosomes (78,79) causing the overlapping of the two organelle markers (80). This hypothesis arises from the particle analysis. This analysis revealed that pH sensitive analogs (CND2-CND4) exhibited a 2 – 3-fold increase in fluorescence intensity after 24 hours compared to other CNDs (**Fig.** S9). Such a substantial surge in intensity could be due to the probes accumulating in autolysosomes, where the acidic environment inhibits the photoinduced electron transfer (PET) process in the naphthalimide group, resulting in amplified fluorescence. It seems rather unlikely that CND2-CND4 intensity increase is caused simply due to the accumulation of these analogs in lipid droplets.

In subsequent studies, the subcellular distribution of CNDs, especially among endosomes, lysosomes, and lipid droplets, can be better elucidated by using highly specific organelle markers as well as different loading methods (e.g. using methyl-β-cyclodextrin or LDL complexes) (57,81).

In summary, in this report we present the development of novel fluorescent cholesteryl probes, featuring a modular design that incorporates a 1,8-naphthalimide scaffold. This bifunctional scaffold is notable for its fluorescence and solvatochromic properties and provide opportunity to incorporate numerous head groups and linkers connecting it with cholesterol moiety. Our probes possess many unique features, making them very useful for studies where fluorescent cholesterol analogs are needed. Recognizing the crucial role of the hydroxyl group in cholesterol’s interactions with different lipids, including sphingomyelin (SM) and proteins, inspired us to incorporate an additional hydroxyl group into the design of the CND probes, either in linker or head group. We also intend to take advantage of the additional -OH group by esterifying it with oleic acid to obtain fluorescent analogs of cholesteryl oleate.

To the best of our knowledge, CND2, CND3, and CND4 represent a novel class of pH-dependent fluorescent cholesteryl probes, readily uptaken by cells. On average, they partition better than NBD-cholesterol analogs and CND3 shows partitioning comparable to that of Bchol. This observation is of interest for those studying lipid and membrane dynamics and partitioning. Additionally, for research into subcellular cholesterol trafficking, analogs such as CND1, CND6, and CND8 are promising due to their specific interactions and behaviors. Additionally, at the concentration tested the CND probes were not toxic to cells (cells showed healthy morphology) and fluorescent puncta were observed even two days after pulse-chase exposure.

The modular nature of our system facilitates the easy synthesis and application of these fluorescent CND analogs for live cell and tissue studies. Our initial experiments, especially those conducted on brain slices using two-photon fluorescence microscopy, have shown promising results. The introduction of these environment-sensitive analogs represents a valuable addition to the current collection of fluorescent cholesterol probes, particularly due to their pH sensitivity. We anticipate that these compounds will be useful not only for live cell imaging but also for areas such as cellular targeting and drug delivery systems, expanding the possibilities for scientific exploration and application.

## Supplemental Data

- Supplementary figures, synthesis procedures and characterization of compounds (PDF).
- **Movie S1**. Visualization of molecular dynamics simulation trajectory of CND2 and CND3 in SM:Chol:POPC system (AVI).
- **Movie S2**. Time-lapse of CND2 in 3T3 fibroblast at 24h after incubation showing the movement of fluorescent puncta containing CND2 (green) and LysoView (red) (AVI).

## Acknowledgments

We would also like to extend our sincere appreciation to the Florida Atlantic University Office of Undergraduate Research and Inquiry for their invaluable support of undergraduate research endeavors.

## Abbreviations

(CND): Cholesterol-naphthalimide probe

