## Supplementary_Data for "Modular Fluorescent Cholesterol Naphthalimide Probes And Their Application For Cholesterol Trafficking Studies In Cells"

### Table of Contents


PAGE

**Supplementary figures and tables referenced in the main text**

Photophysical characterization of compounds S3-S10
Molecular Dynamics related data S11-13
Cell images. Particle masks and particle quantification S14-S39

**NMR and MS spectra of all intermediates and final compounds**

NMR spectra S55
MS spectra S72


**Movie S1.** Visualization of molecular dynamics simulation trajectory of CND2 and CND3 in SM:Chol:POPC system. (AVI)

**Movie S2.** Time-lapse of CND2 in 3T3 fibroblast at 24h after incubation showing the movement of fluorescent puncta containing CND2 (green) and LysoView (red). (AVI)

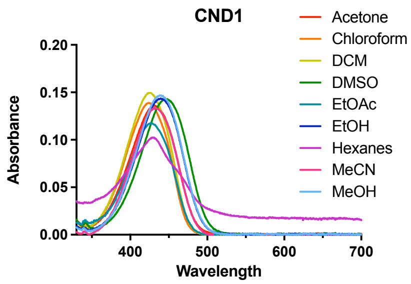
CND1
Absorbance
Wavelength
0.20
0.15
0.10
0.05
0.00
400
500
600
700
Acetone
Chloroform
DCM
DMSO
EtOAc
EtOH
Hexanes
MeCN
MeOH

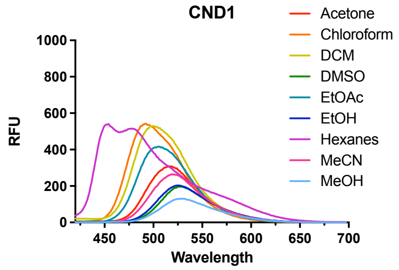
CND1
RFU
Wavelength
1000
800
600
400
200
0
450
500
550
600
650
700
Acetone
Chloroform
DCM
DMSO
EtOAc
EtOH
Hexanes
MeCN
MeOH

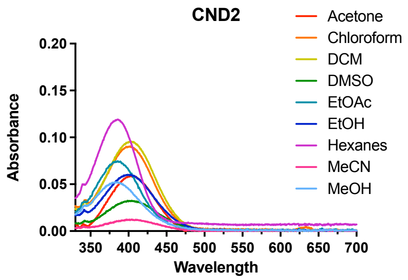
CND2
Absorbance
Wavelength
0.20
0.15
0.10
0.05
0.00
350
400
450
500
550
600
650
700
Acetone
Chloroform
DCM
DMSO
EtOAc
EtOH
Hexanes
MeCN
MeOH

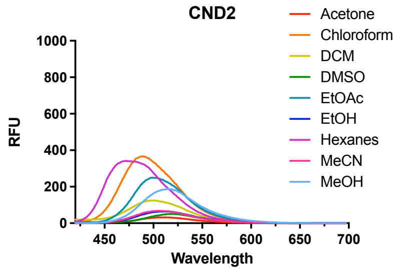
CND2
RFU
Wavelength
1000
800
600
400
200
0
450
500
550
600
650
700
Acetone
Chloroform
DCM
DMSO
EtOAc
EtOH
Hexanes
MeCN
MeOH

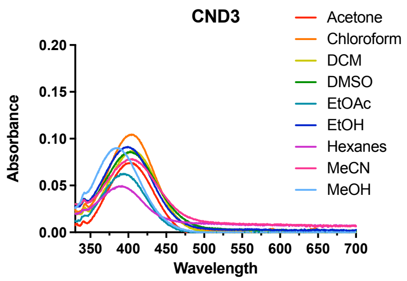
CND3
Absorbance
Wavelength
0.20
0.15
0.10
0.05
0.00
350
400
450
500
550
600
650
700
Acetone
Chloroform
DCM
DMSO
EtOAc
EtOH
Hexanes
MeCN
MeOH

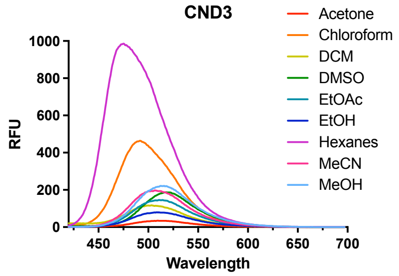
CND3
RFU
Wavelength
1000
800
600
400
200
0
450
500
550
600
650
700
Acetone
Chloroform
DCM
DMSO
EtOAc
EtOH
Hexanes
MeCN
MeOH

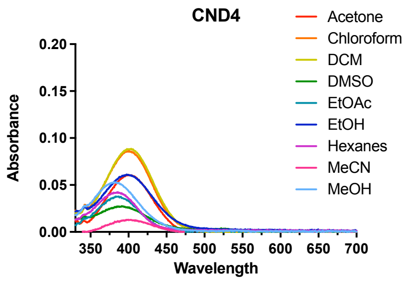
CND4
Absorbance
Wavelength
0.20
0.15
0.10
0.05
0.00
350
400
450
500
550
600
650
700
Acetone
Chloroform
DCM
DMSO
EtOAc
EtOH
Hexanes
MeCN
MeOH

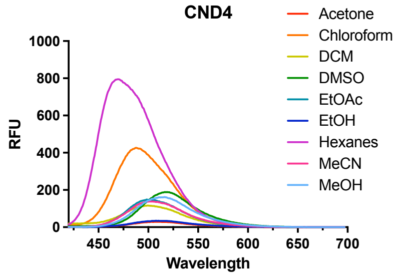
CND4
RFU
Wavelength
1000
800
600
400
200
0
450
500
550
600
650
700
Acetone
Chloroform
DCM
DMSO
EtOAc
EtOH
Hexanes
MeCN
MeOH

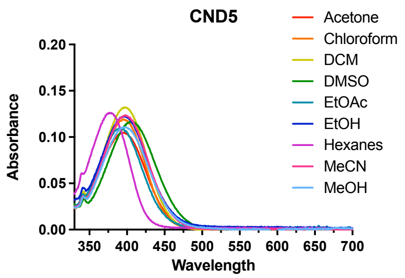
CND5
Absorbance
Wavelength
0.20
0.15
0.10
0.05
0.00
350
400
450
500
550
600
650
700
Acetone
Chloroform
DCM
DMSO
EtOAc
EtOH
Hexanes
MeCN
MeOH

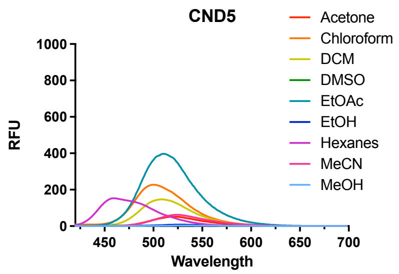
CND5
RFU
Wavelength
1000
800
600
400
200
0
450
500
550
600
650
700
Acetone
Chloroform
DCM
DMSO
EtOAc
EtOH
Hexanes
MeCN
MeOH

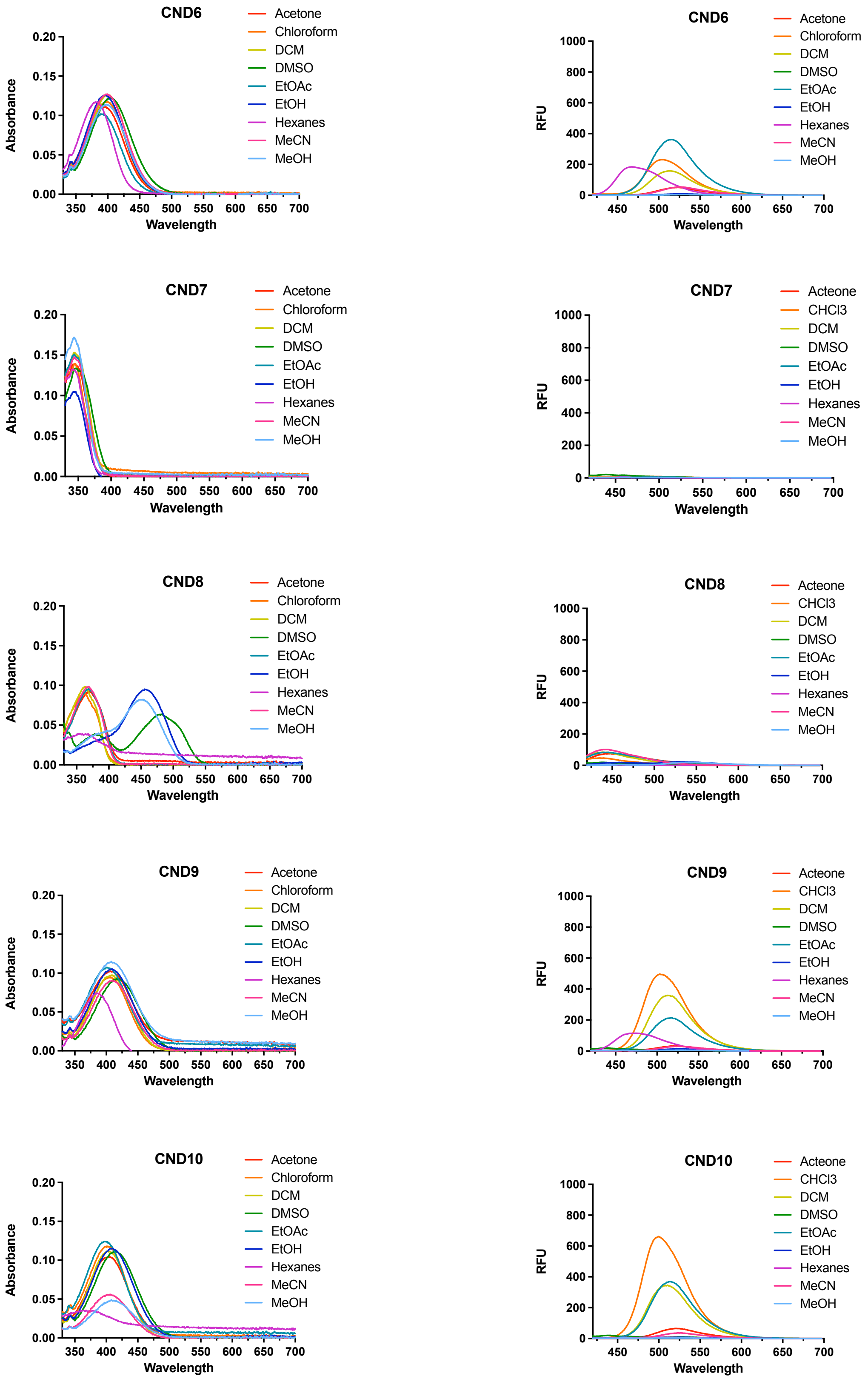


**Fig. S1**. The absorbance and emission spectra of CND analogs in various solvents.

**Table S1.** Absorbance (abs), emission (em) maxima and Stokes' shift (Δλ) for all CND analogs. Values provided in nanometers and wavenumbers. Values not obtained are denoted as n/a.

| CND1 | | | | | | |
|---|---|---|---|---|---|---|
| Solvent | λ abs (nm) | λ em (nm) | Δλ (nm) | ν abs ($cm^{-1}$) | ν em ($cm^{-1}$) | Δν ($cm^{-1}$) |
| Acetone | 431 | 519 | 88 | 23202 | 19268 | 3934 |
| Chloroform | 425 | 492 | 67 | 23529 | 20325 | 3204 |
| DCM | 425 | 499 | 74 | 23529 | 20040 | 3489 |
| MeCN | 431 | 519 | 88 | 23202 | 19268 | 3934 |
| DMSO | 446 | 528 | 82 | 22422 | 18939 | 3482 |
| EtOH | 440 | 525 | 85 | 22727 | 19048 | 3680 |
| Hexanes | 429 | 454, 478 | 25, 49 | 23310 | 22026, 20920 | 1284, 2390 |
| MeOH | 440 | 530 | 90 | 22727 | 18868 | 3859 |
| EtOAc | 427 | 506 | 79 | 23419 | 19762 | 3656 |

| CND2 | | | | | | |
|---|---|---|---|---|---|---|
| Solvent | λ abs (nm) | λ em (nm) | Δλ (nm) | ν abs ($cm^{-1}$) | ν em ($cm^{-1}$) | Δν ($cm^{-1}$) |
| Acetone | 403 | 512 | 109 | 24814 | 19531 | 5283 |
| Chloroform | 402 | 490 | 88 | 24876 | 20408 | 4467 |
| DCM | 402 | 501 | 99 | 24876 | 19960 | 4916 |
| MeCN | 406 | 505 | 99 | 24631 | 19802 | 4829 |
| DMSO | 403 | 519 | 116 | 24814 | 19268 | 5546 |
| EtOH | 400 | 511 | 111 | 25000 | 19569 | 5431 |
| Hexanes | 386 | 472 | 86 | 25907 | 21186 | 4720 |
| MeOH | 386 | 515 | 129 | 25907 | 19417 | 6489 |
| EtOAc | 386 | 499 | 113 | 25906 | 20040 | 5866 |

| CND3 | | | | | | |
|---|---|---|---|---|---|---|
| Solvent | λ abs (nm) | λ em (nm) | Δλ (nm) | ν abs ($cm^{-1}$) | ν em ($cm^{-1}$) | Δν ($cm^{-1}$) |
| Acetone | 401 | 510 | 109 | 24938 | 19608 | 5330 |
| Chloroform | 401 | 492 | 91 | 24938 | 20325 | 4612 |
| DCM | 406 | 503 | 97 | 24631 | 19881 | 4750 |
| MeCN | 404 | 507 | 103 | 24752 | 19724 | 5029 |
| DMSO | 403 | 518 | 115 | 24814 | 19305 | 5509 |
| EtOH | 399 | 509 | 110 | 25063 | 19646 | 5416 |
| Hexanes | 392 | 475 | 83 | 25510 | 21053 | 4458 |
| MeOH | 386 | 513 | 127 | 25907 | 19493 | 6414 |
| EtOAc | 395 | 511 | 116 | 25316 | 19569 | 5746 |

| CND4 | | | | | | |
|---|---|---|---|---|---|---|
| Solvent | λ abs (nm) | λ em (nm) | Δλ (nm) | ν abs ($cm^{-1}$) | ν em ($cm^{-1}$) | Δν ($cm^{-1}$) |
| Acetone | 399 | 503 | 104 | 25063 | 19881 | 5182 |
| Chloroform | 401 | 488 | 87 | 24938 | 20492 | 4446 |
| DCM | 400 | 499 | 99 | 25000 | 20040 | 4960 |
| MeCN | 399 | 504 | 105 | 25063 | 19841 | 5221 |
| DMSO | 391 | 517 | 126 | 25575 | 19342 | 6233 |
| EtOH | 398 | 508 | 110 | 25126 | 19685 | 5441 |
| Hexanes | 385 | 470 | 85 | 25974 | 21053 | 4921 |
| MeOH | 381 | 517 | 136 | 26247 | 19342 | 6904 |
| EtOAc | 383 | 499 | 116 | 26109 | 20040 | 6069 |

| CND5 | | | | | | |
|---|---|---|---|---|---|---|
| Solvent | λ abs (nm) | λ em (nm) | Δλ (nm) | ν abs ($cm^{-1}$) | ν em ($cm^{-1}$) | Δν ($cm^{-1}$) |
| Acetone | 396 | 520 | 124 | 25253 | 19231 | 6022 |
| Chloroform | 397 | 500 | 103 | 25189 | 20000 | 5189 |
| DCM | 397 | 509 | 112 | 25189 | 19646 | 5543 |
| MeCN | 398 | 524 | 126 | 25126 | 19084 | 6042 |
| DMSO | 405 | 528 | 123 | 24691 | 18939 | 5752 |
| EtOH | 397 | 528 | 131 | 25189 | 18939 | 6250 |
| Hexanes | 378 | 459 | 81 | 26455 | 21786 | 4669 |
| MeOH | 395 | 530 | 135 | 25316 | 18868 | 6449 |
| EtOAc | 390 | 510 | 120 | 25641.03 | 19607 | 6033 |

| CND6 | | | | | | |
|---|---|---|---|---|---|---|
| Solvent | λ abs (nm) | λ em (nm) | Δλ (nm) | ν abs ($cm^{-1}$) | ν em ($cm^{-1}$) | Δν ($cm^{-1}$) |
| Acetone | 396 | 524 | 128 | 25253 | 19084 | 6169 |
| Chloroform | 398 | 505 | 107 | 25126 | 19802 | 5324 |
| DCM | 401 | 513 | 112 | 24938 | 19493 | 5444 |
| MeCN | 399 | 525 | 126 | 25063 | 19048 | 6015 |
| DMSO | 405 | 528 | 123 | 24691 | 18939 | 5752 |
| EtOH | 397 | 525 | 128 | 25189 | 19048 | 6141 |
| Hexanes | 384 | 467 | 83 | 26042 | 21413 | 4628 |
| MeOH | 398 | 529 | 131 | 25126 | 18904 | 6222 |
| EtOAc | 389 | 515 | 126 | 25706 | 19417 | 6289 |

| CND7 | | | | | | |
|---|---|---|---|---|---|---|
| Solvent | λ abs (nm) | λ em (nm) | Δλ (nm) | ν abs ($cm^{-1}$) | ν em ($cm^{-1}$) | Δν ($cm^{-1}$) |
| Acetone | 345 | 460 | 115 | 28986 | 21739 | 7246 |
| Chloroform | 345 | 501 | 156 | 28986 | 19960 | 9025 |
| DCM | 343 | 499 | 156 | 29155 | 20040 | 9114 |
| MeCN | 344 | 499 | 155 | 29070 | 20040 | 9030 |
| DMSO | 346 | 440 | 94 | 28902 | 22727 | 6174 |
| EtOH | 343 | 505 | 162 | 29155 | 19802 | 9353 |
| Hexanes | 342 | 459 | 117 | 29240 | 21786 | 7453 |
| MeOH | 344 | 460 | 116 | 29070 | 21739 | 7331 |
| EtOAc | 344 | 492 | 148 | 29070 | 20325 | 8745 |

| CND8 | | | | | | |
|---|---|---|---|---|---|---|
| Solvent | λ abs (nm) | λ em (nm) | Δλ (nm) | ν abs ($cm^{-1}$) | ν em ($cm^{-1}$) | Δν ($cm^{-1}$) |
| Acetone | 368 | 448 | 80 | 27174 | 22321 | 4852 |
| Chloroform | 362 | 435 | 73 | 27624 | 22989 | 4636 |
| DCM | 363 | 438 | 75 | 27548 | 22831 | 4717 |
| MeCN | 369 | 443 | 74 | 27100 | 22573 | 4527 |
| DMSO | 481 | n/a | n/a | 20790 | n/a | n/a |
| EtOH | 456 | 533 | 77 | 21930 | 18762 | 3168 |
| Hexanes | 354 | 460 | 106 | 28249 | 21739 | 6509 |
| MeOH | 453 | 541 | 88 | 22075 | 18484 | 3591 |
| EtOAc | 370 | 440 | 70 | 27027 | 22727 | 4300 |

| CND9 | | | | | | |
|---|---|---|---|---|---|---|
| Solvent | λ abs (nm) | λ em (nm) | Δλ (nm) | ν abs ($cm^{-1}$) | ν em ($cm^{-1}$) | Δν ($cm^{-1}$) |
| Acetone | 406 | 526 | 120 | 24631 | 19011 | 5619 |
| Chloroform | 407 | 503 | 96 | 24570 | 19881 | 4689 |
| DCM | 409 | 513 | 104 | 24450 | 19493 | 4957 |
| MeCN | 406 | 525 | 119 | 24631 | 19048 | 5583 |
| DMSO | 419 | n/a | n/a | 23866 | n/a | n/a |
| EtOH | 409 | 530 | 121 | 24450 | 18868 | 5582 |
| Hexanes | 387 | 472 | 85 | 25840 | 21186 | 4653 |
| MeOH | 408 | 528 | 120 | 24510 | 18939 | 5570 |
| EtOAc | 401 | 517 | 116 | 24938 | 19342 | 5595 |

| CND10 | | | | | | |
|---|---|---|---|---|---|---|
| Solvent | λ abs (nm) | λ em (nm) | Δλ (nm) | ν abs ($cm^{-1}$) | ν em ($cm^{-1}$) | Δν ($cm^{-1}$) |
| Acetone | 399 | 522 | 123 | 25063 | 19157 | 5906 |
| Chloroform | 401 | 500 | 99 | 24938 | 20000 | 4938 |
| DCM | 401 | 511 | 110 | 24938 | 19569 | 5368 |
| MeCN | 402 | 523 | 121 | 24876 | 19120 | 5755 |
| DMSO | 416 | n/a | n/a | 24038 | n/a | n/a |
| EtOH | 409 | 527 | 118 | 24450 | 18975 | 5475 |
| Hexanes | 362 | 493 | 131 | 27624 | 20284 | 7340 |
| MeOH | 408 | n/a | n/a | 24510 | n/a | n/a |
| EtOAc | 397 | 512 | 115 | 25189 | 19531 | 5658 |

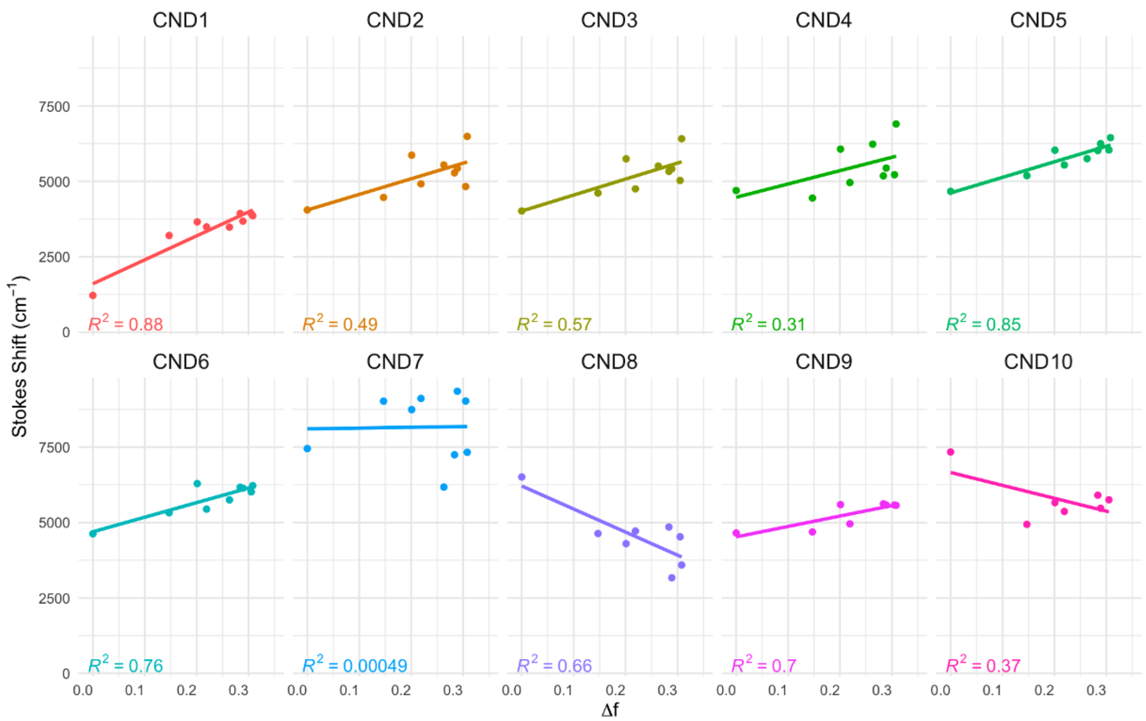


**Fig. S2.** Lippert-Mataga plot for CND series in selected organic solvents. Solvents tested in increasing order of orientation polarizability: hexanes, CHCl3, EtOAc, DCM, DMSO, acetone, EtOH, MeCN, MeOH. For CND8, CND9, and CND10 DMSO was omitted due to no detectable fluorescence. For CND10, MeOH was also omitted. Compounds were tested at 1 μM.

***Table S2***. Molar extinction coefficients of the CND series with maximum wavelength of absorption for $CHCl_3$ and DMSO.

| **Solvent** | **$CHCl_3$** | | **DMSO** | |
|---|---|---|---|---|
| *Analog* | ε ($M^{-1}$ $cm^{-1}$) | $\lambda_{abs}$ (nm) | ε ($M^{-1}$ $cm^{-1}$) | $\lambda_{abs}$ (nm) |
| *CND1* | 12850 | 425 | 16590 | 446 |
| *CND2* | 10170 | 402 | 9398 | 403 |
| *CND3* | 9455 | 401 | 10810 | 403 |
| *CND4* | 9228 | 401 | 8839 | 391 |
| *CND5* | 12000 | 397 | 11470 | 405 |
| *CND6* | 12510 | 398 | 11450 | 405 |
| *CND7* | 14830 | 345 | 13310 | 346 |
| *CND8* | 10240 | 362 | 12170 | 481 |
| *CND9* | 10420 | 407 | 11460 | 419 |
| *CND10* | 12840 | 401 | 10680 | 416 |

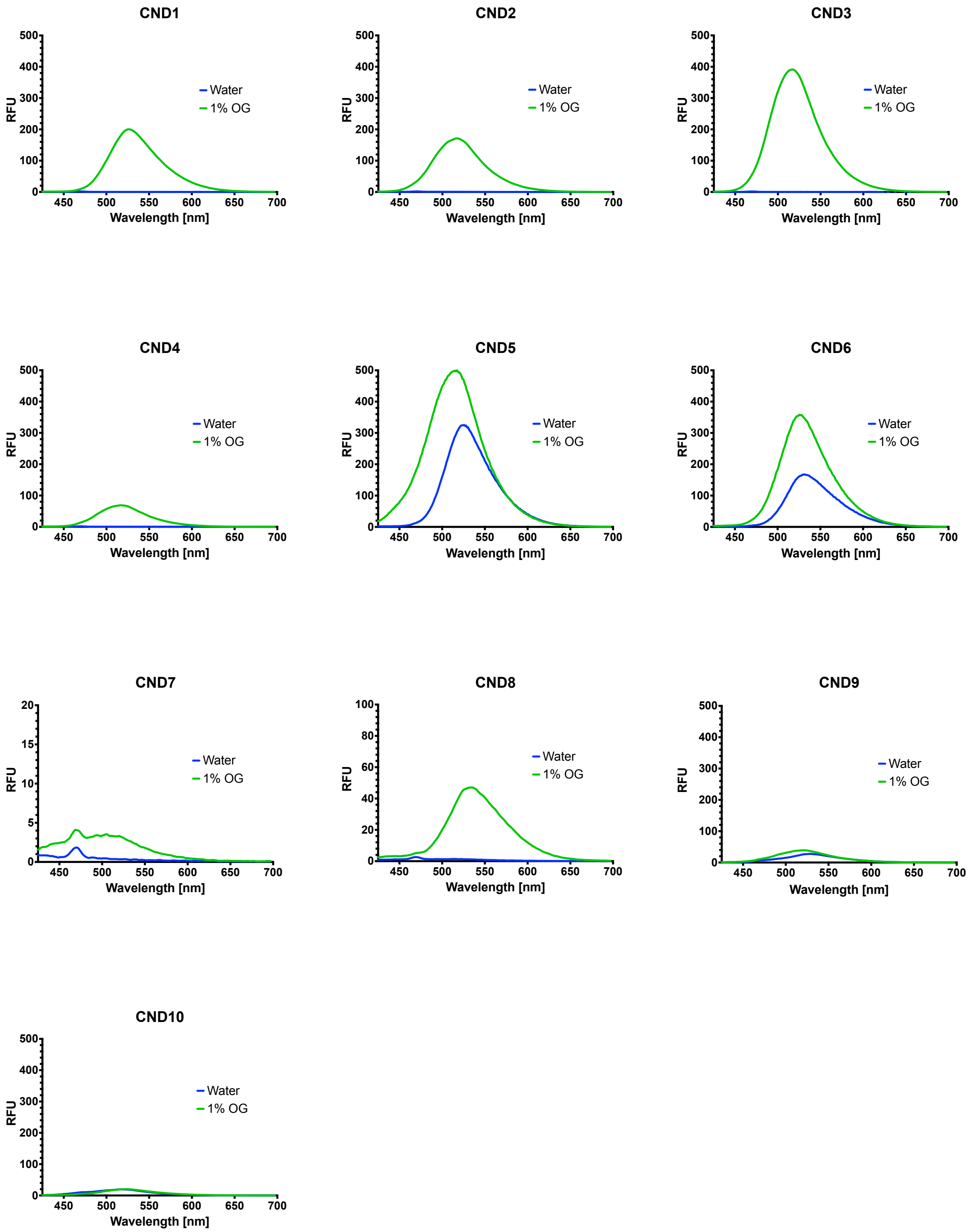


**Fig. S3.** Fluorescence emission spectra of CND analogs in 1% octyl glucoside micellar solution (green) and water (blue) showing membrane sensing and aggregation properties. Compounds were present at 1μM concentration.

**Table S3.** Lipid bilayer composition used for molecular dynamics simulations in this study.

| System Name | POPC (mol %) | Chol (mol %) | SM (mol %) |
|---|---|---|---|
| 100% POPC | 100 | 0 | 0 |
| 5% Chol | 95 | 5 | 0 |
| 25% Chol | 75 | 25 | 0 |
| 40% Chol | 60 | 40 | 0 |
| SM:Chol:POPC (1:1:2) | 50 | 25 | 25 |

**Table S4.** Cholesterol tilt angle analysis from molecular dynamics trajectories. Average sterol tilt angles with standard error of mean for cholesterol and all tested fluorescent cholesterol probes.

| Analog | 100% POPC | 5% Chol | 25% Chol | 40% Chol | SM:Chol:POPC |
|---|---|---|---|---|---|
| Cholesterol | - | 25 ± 0.23 | 19 ± 0.07 | 14 ± 0.06 | 15 ± 0.05 |
| CND1 | 56 ± 2.10 | 33 ± 1.38 | 34 ± 2.58 | 25 ± 1.52 | 10 ± 0.60 |
| CND2 | 47 ± 3.08 | 41 ± 2.38 | 24 ± 1.08 | 17 ± 1.06 | 21 ± 1.01 |
| CND3 | 36 ± 1.50 | 36 ± 2.24 | 22 ± 1.03 | 21 ± 0.81 | 15 ± 0.93 |
| CND4 | 37 ± 1.88 | 38 ± 2.33 | 37 ± 4.34 | 15 ± 0.92 | 16 ± 0.92 |
| CND5 | - | 47 ± 1.53 | 40 ± 2.10 | 21 ± 1.21 | 42 ± 1.94 |
| CND6 | - | 47 ± 1.29 | 25 ± 1.35 | 15 ± 0.83 | 15 ± 1.00 |
| CND7 | - | - | 28 ± 1.15 | - | 17 ± 1.04 |
| CND8 | - | - | 22 ± 1.15 | - | 23 ± 1.63 |
| CND9 | - | - | 22 ± 1.14 | - | 23 ± 1.06 |
| CND10 | - | - | 25 ± 1.35 | - | 21 ± 0.89 |
| 22NBD | - | - | 17 ± 1.23 | - | 16 ± 1.15 |
| 25NBD | - | - | 34 ± 1.51 | - | 14 ± 0.71 |
| 3HxNBD | - | - | 22 ± 1.25 | - | 12 ± 0.74 |

Table S5. Average membrane thickness [Å] ± SEM derived from last 80 ns of MD simulation trajectory.

| Analog | 100% POPC | 5% Chol | 25% Chol | 40% Chol | SM:Chol:POPC |
|---|---|---|---|---|---|
| CND1 | 39.8 ± 0.08 | 40.5 ± 0.06 | 44.8 ± 0.06 | 46.3 ± 0.05 | 47.2 ± 0.04 |
| CND2 | 39.4 ± 0.08 | 40.5 ± 0.07 | 45.2 ± 0.06 | 46.2 ± 0.05 | 47.4 ± 0.07 |
| CND3 | 39.1 ± 0.09 | 40.3 ± 0.05 | 44.7 ± 0.05 | 46.6 ± 0.04 | 47.1 ± 0.06 |
| CND4 | 39.6 ± 0.07 | 40.6 ± 0.06 | 44.8 ± 0.05 | 46.8 ± 0.04 | 47.1 ± 0.04 |
| CND5 | - | 40.8 ± 0.07 | 45.1 ± 0.05 | 46.3 ± 0.04 | 47.1 ± 0.05 |
| CND6 | - | 40.5 ± 0.06 | 44.5 ± 0.06 | 46.5 ± 0.04 | 47.1 ± 0.04 |
| CND7 | - | - | 45.0 ± 0.08 | - | 47.5 ± 0.05 |
| CND8 | - | - | 44.8 ± 0.06 | - | 47.0 ± 0.04 |
| CND9 | - | - | 44.4 ± 0.06 | - | 47.1 ± 0.08 |
| CND10 | - | - | 45.0 ± 0.08 | - | 47.3 ± 0.05 |
| 22NBD | - | - | 44.6 ± 0.06 | - | 48.0 ± 0.07 |
| 25NBD | - | - | 44.7 ± 0.06 | - | 47.0 ± 0.07 |
| 3HxNBD | - | - | 44.9 ± 0.05 | - | 47.0 ± 0.05 |

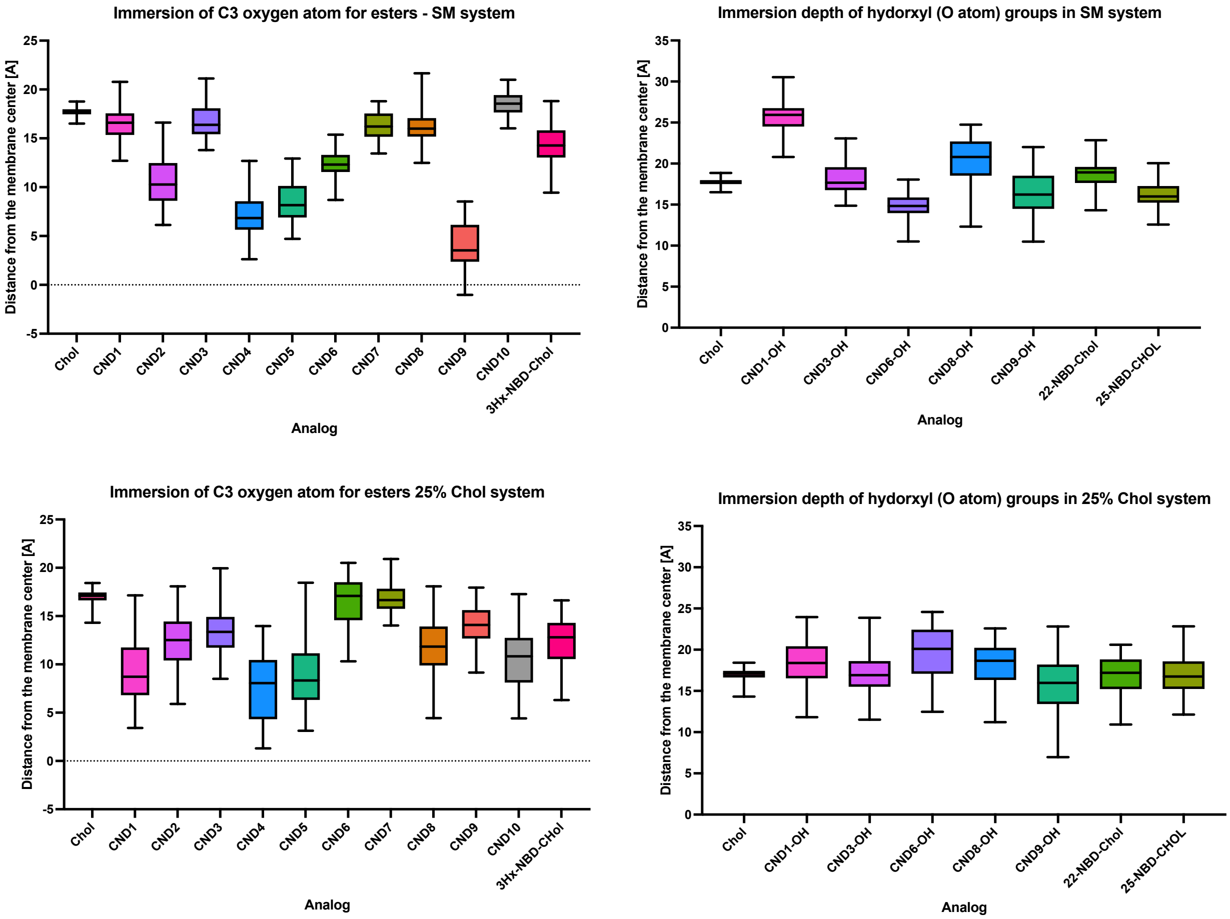


**Fig. S4**. Comparison of the immersion depth for all analogs in Chol:POPC, 1:3 (25% Chol) and the SM:Chol:POPC, 1:1:2 systems measured as position of C3-oxygen atom (left column). Position of available other hydroxyl groups present in some analogs in respect to cholesterol was also determined (right column). Different behavior of probes, depending on lipid composition is evident.

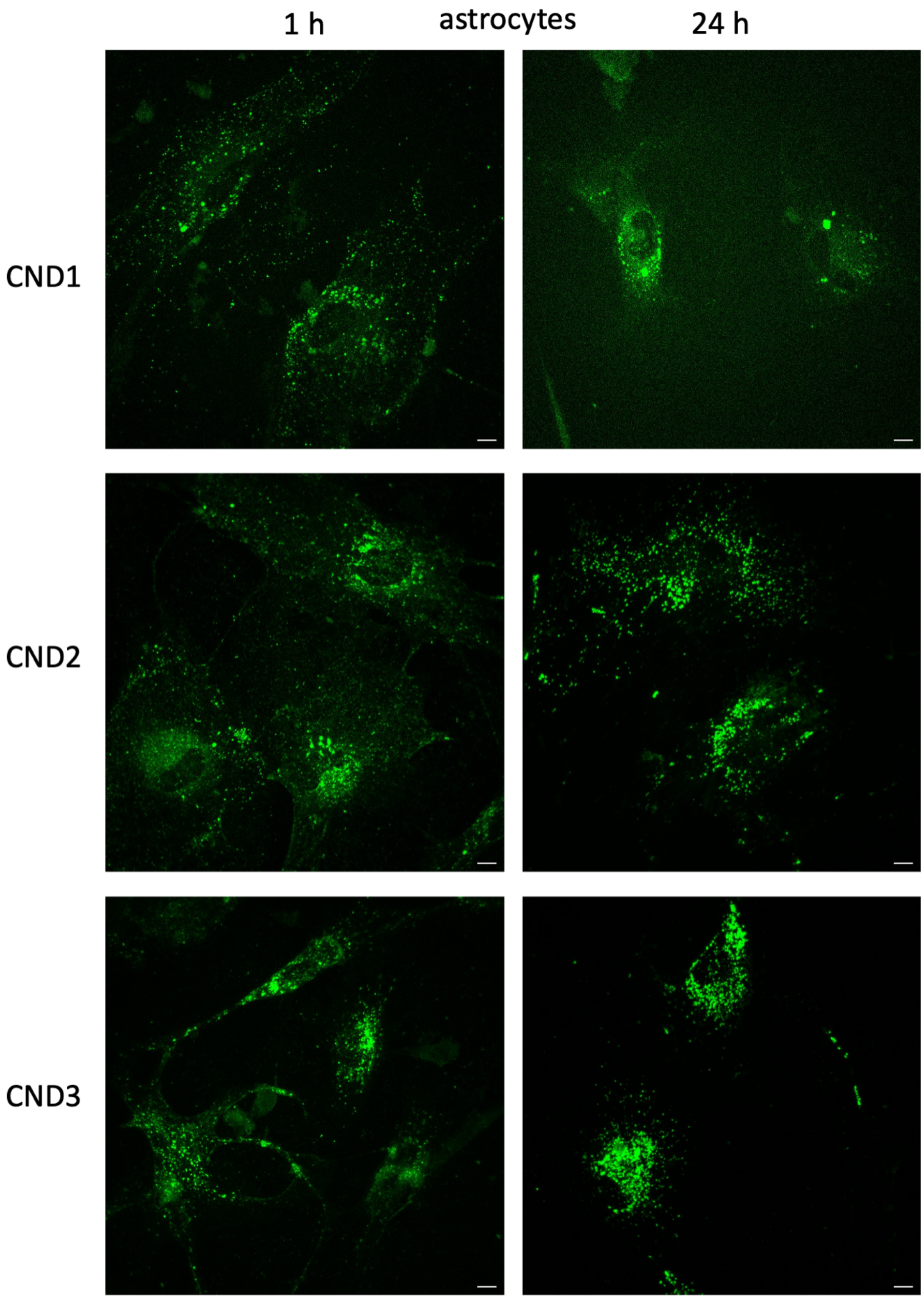
1 h
astrocytes
24 h
CND1
CND2
CND3

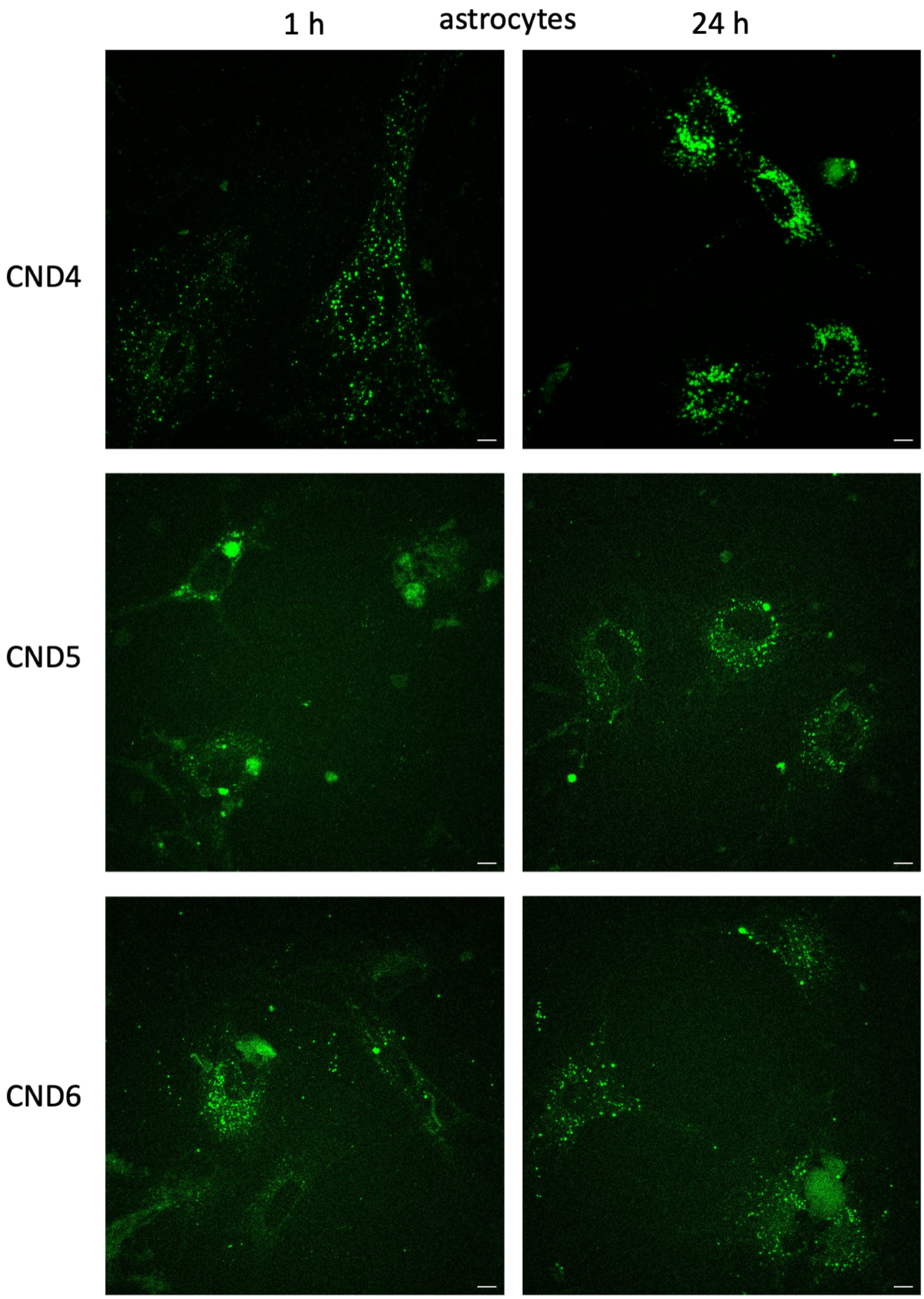
1 h
astrocytes
24 h
CND4
CND5
CND6

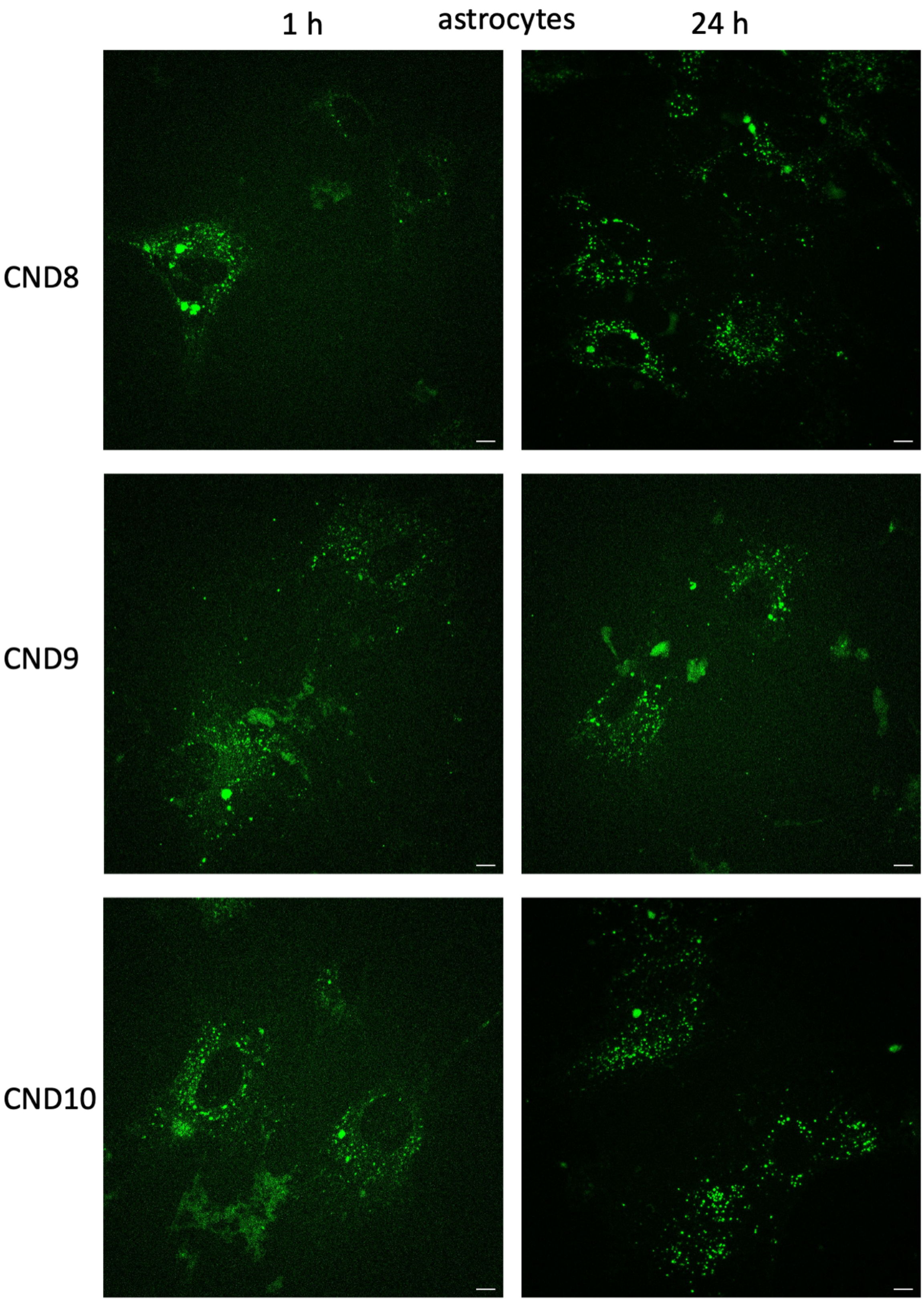
1 h
astrocytes
24 h
CND8
CND9
CND10

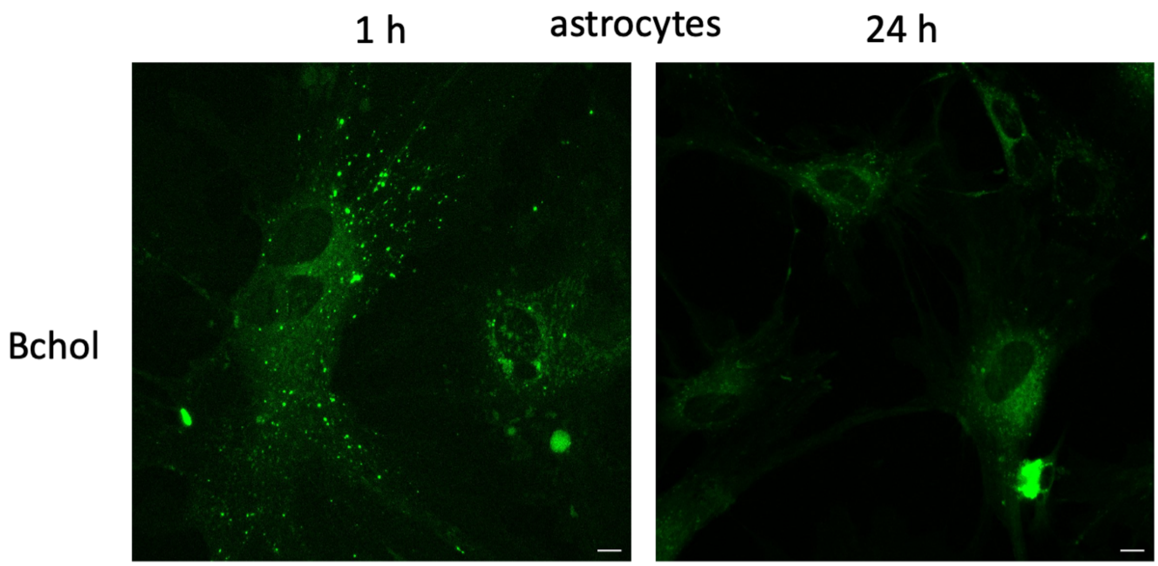
1 h
astrocytes
24 h
Bchol

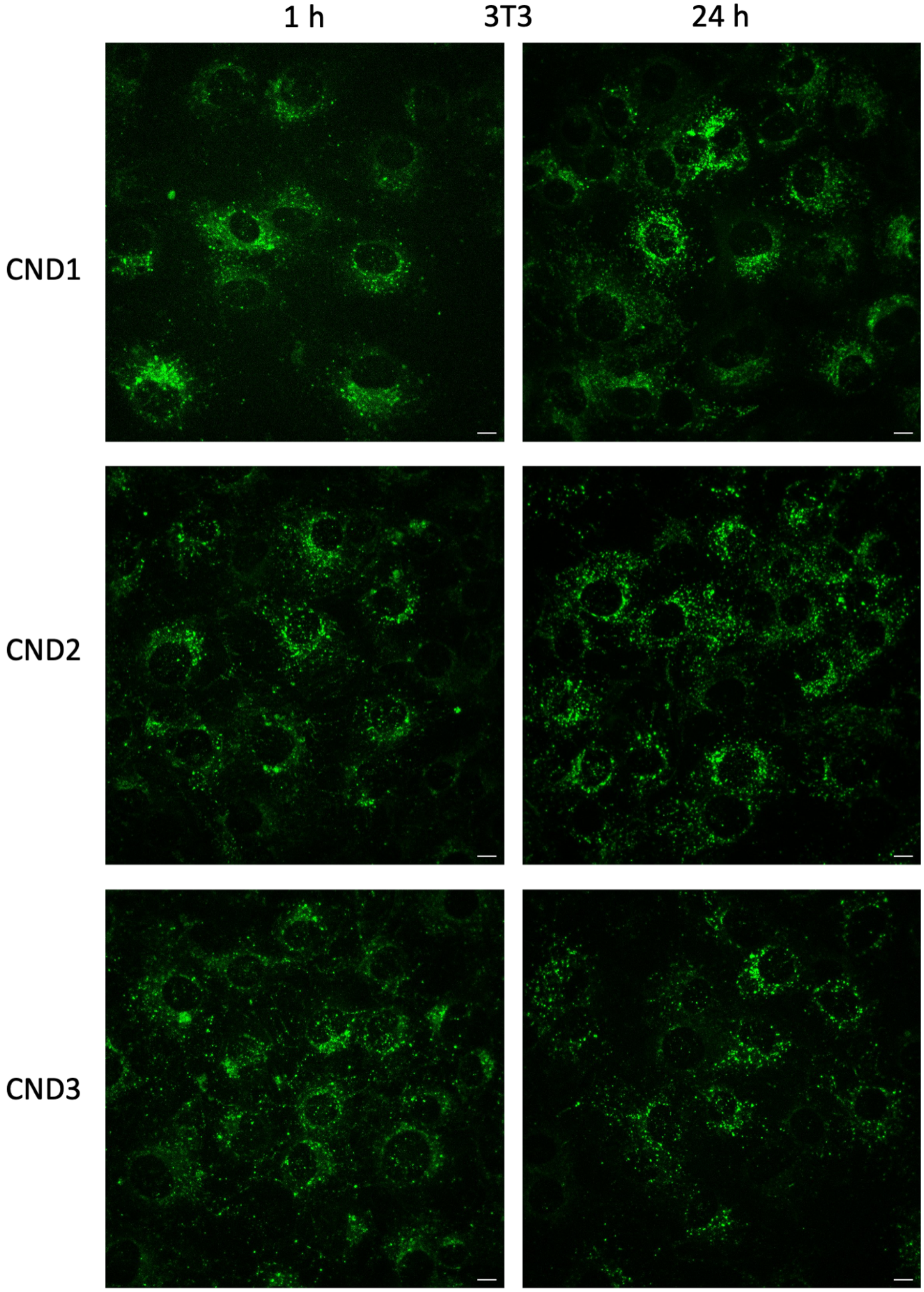
1 h
3T3
24 h
CND1
CND2
CND3

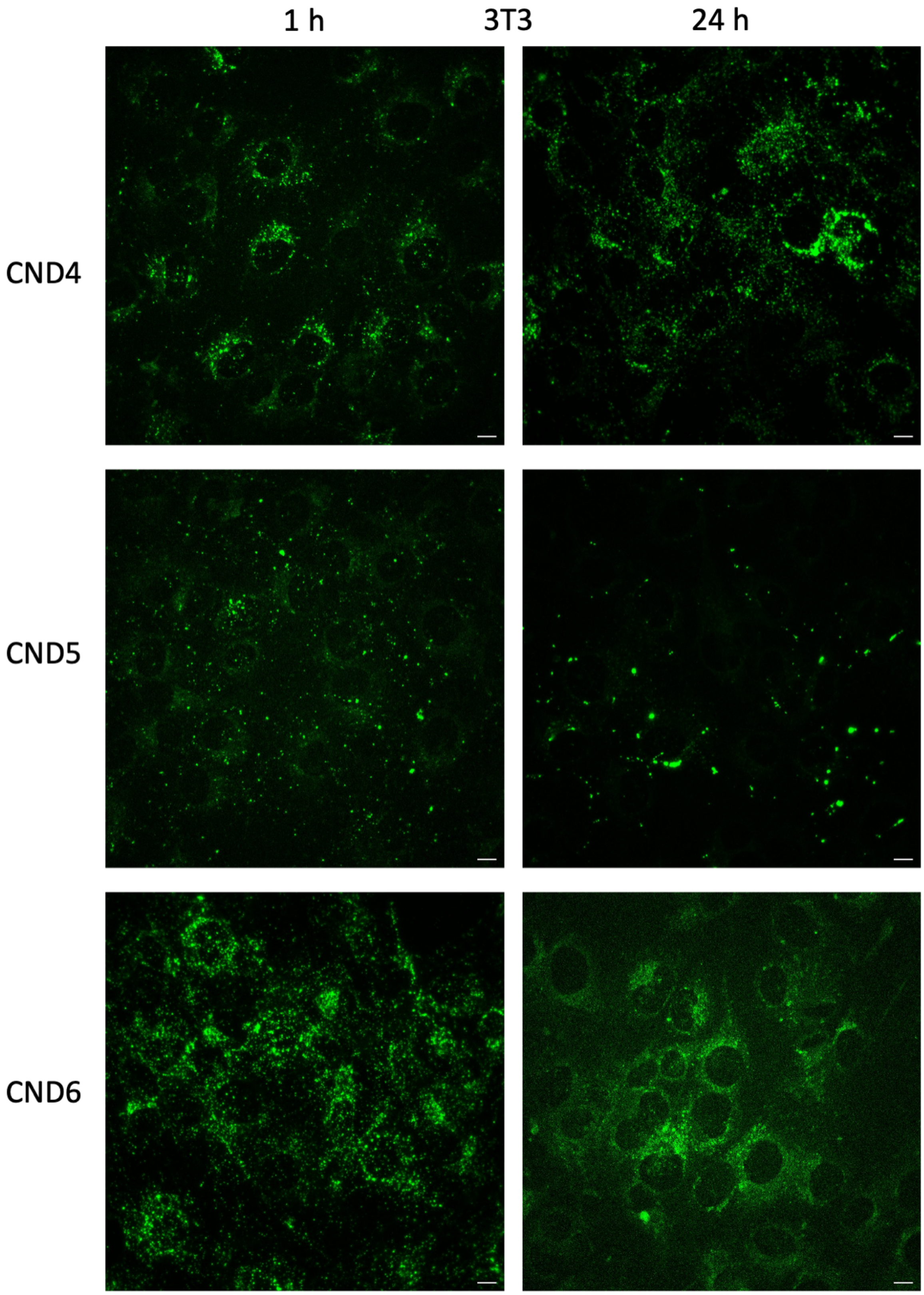

1 h
3T3
24 h
CND4
CND5
CND6

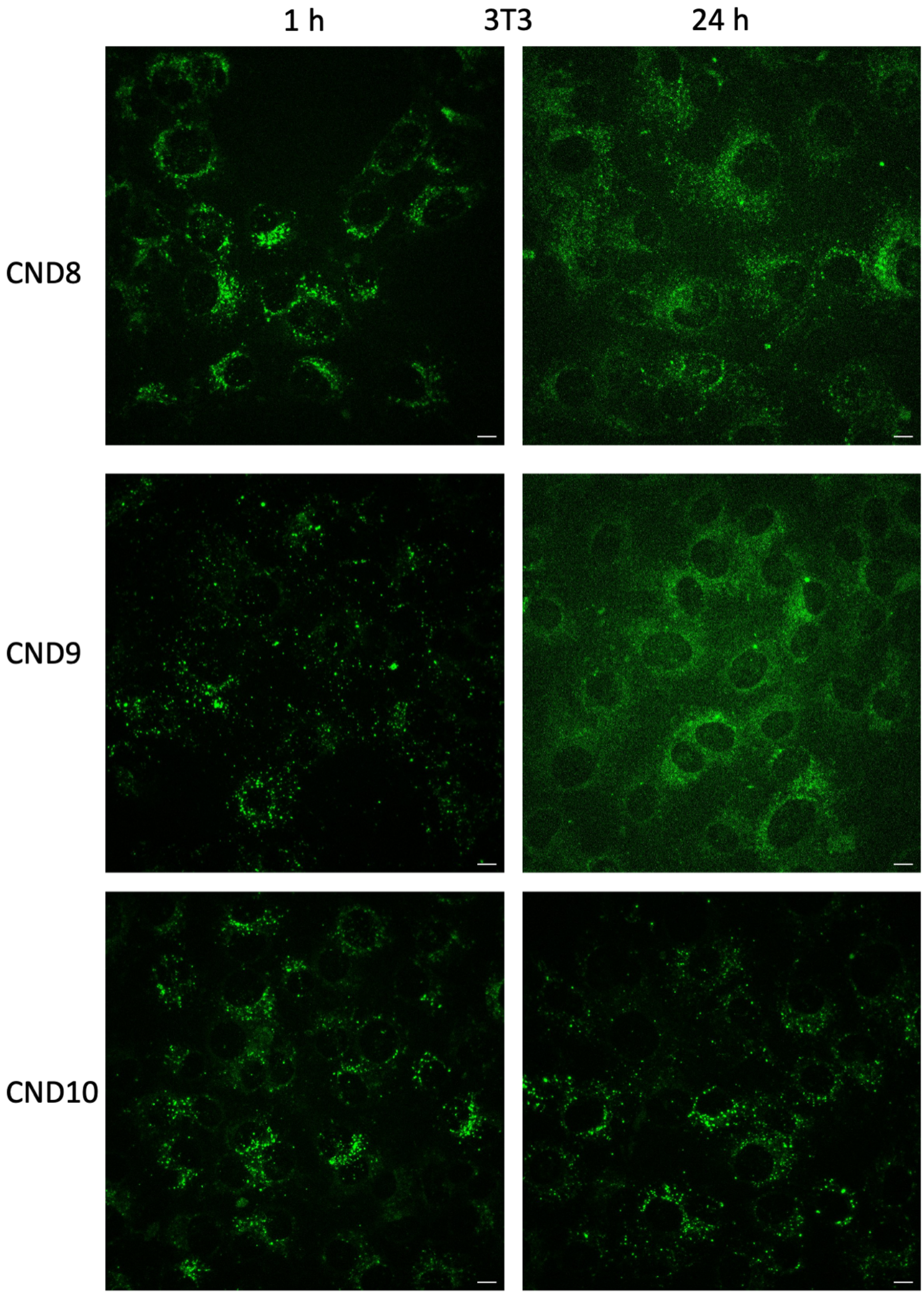
1 h
3T3
24 h
CND8
CND9
CND10

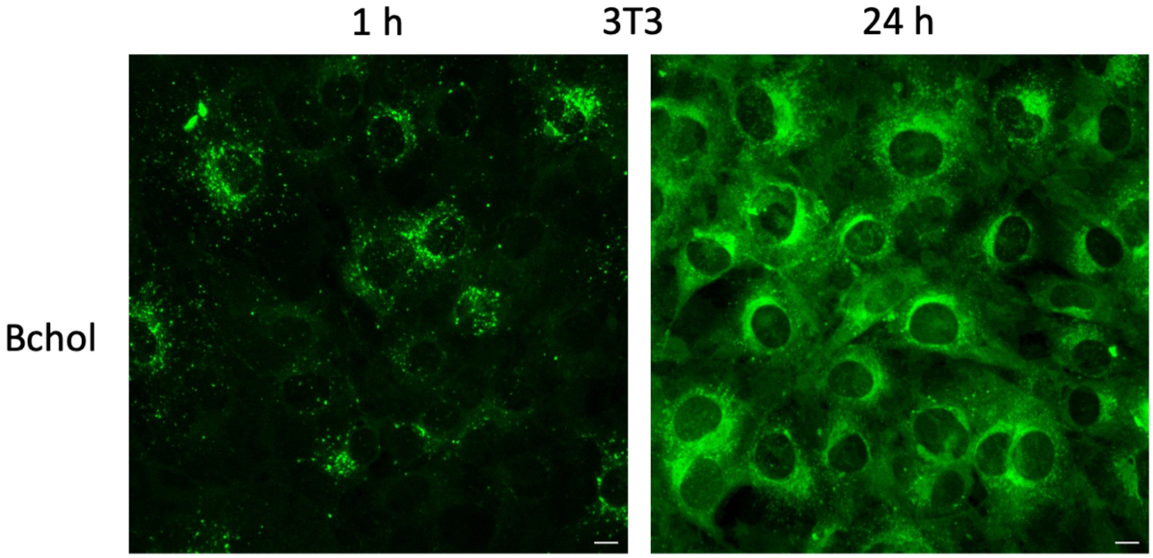


**Fig. S5**. Representative examples of fluorescent confocal images of green channel representing tested probes in astrocytes and 3T3 fibroblasts pulse chased after 1h and 24h. Images depict average intensity of Z-stack. Bar size – 10 μm.

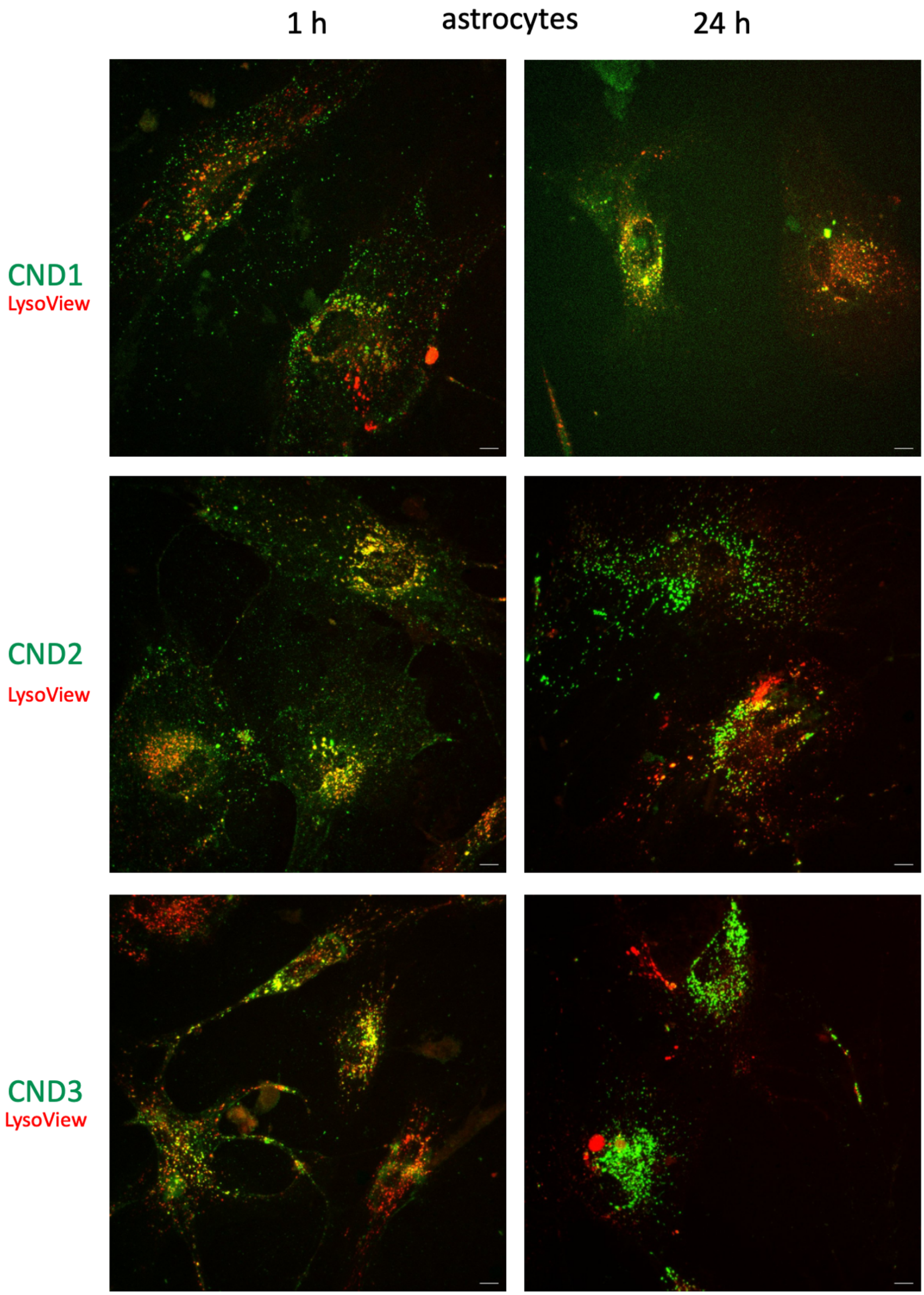
1 h
astrocytes
24 h
CND1
LysoView
CND2
LysoView
CND3
LysoView

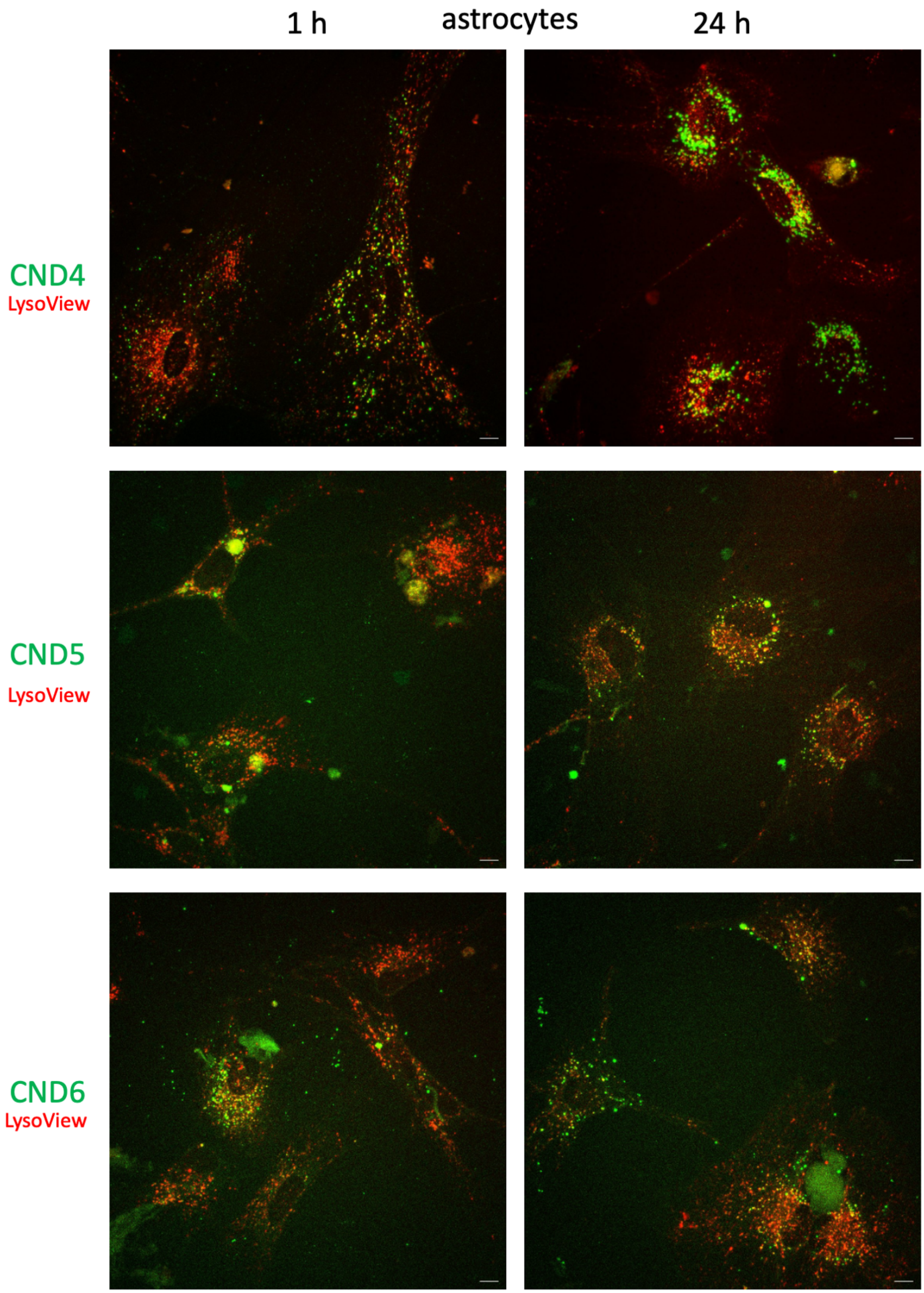
1 h
astrocytes
24 h
CND4
LysoView
CND5
LysoView
CND6
LysoView

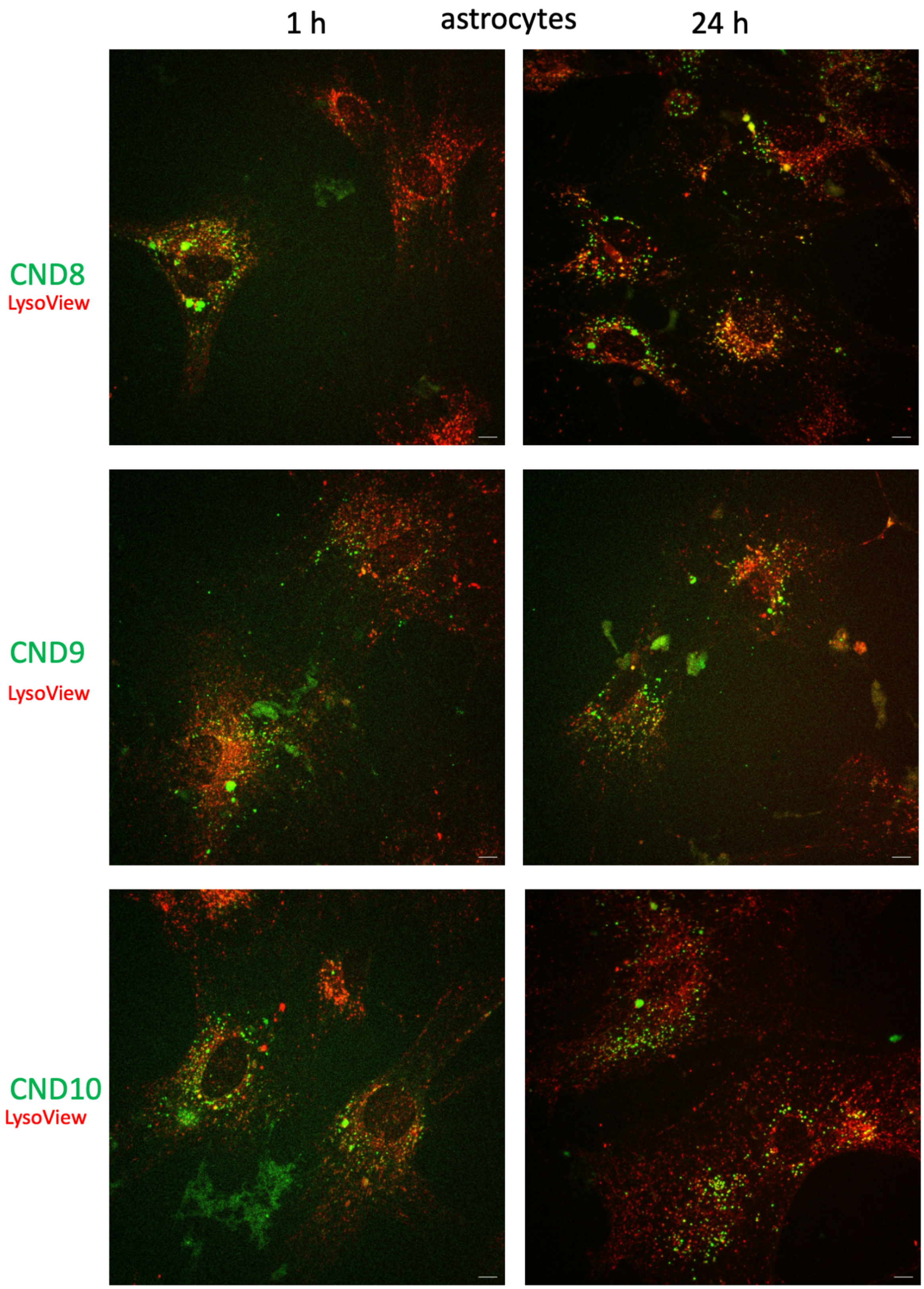
1 h
astrocytes
24 h
CND8
LysoView
CND9
LysoView
CND10
LysoView

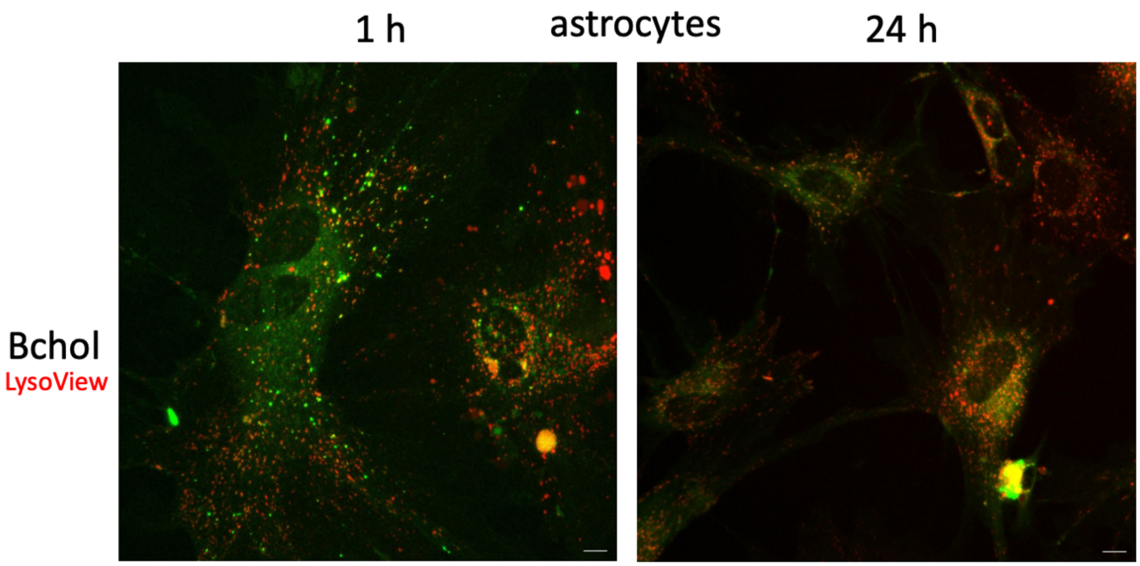
1 h
astrocytes
24 h
Bchol
LysoView

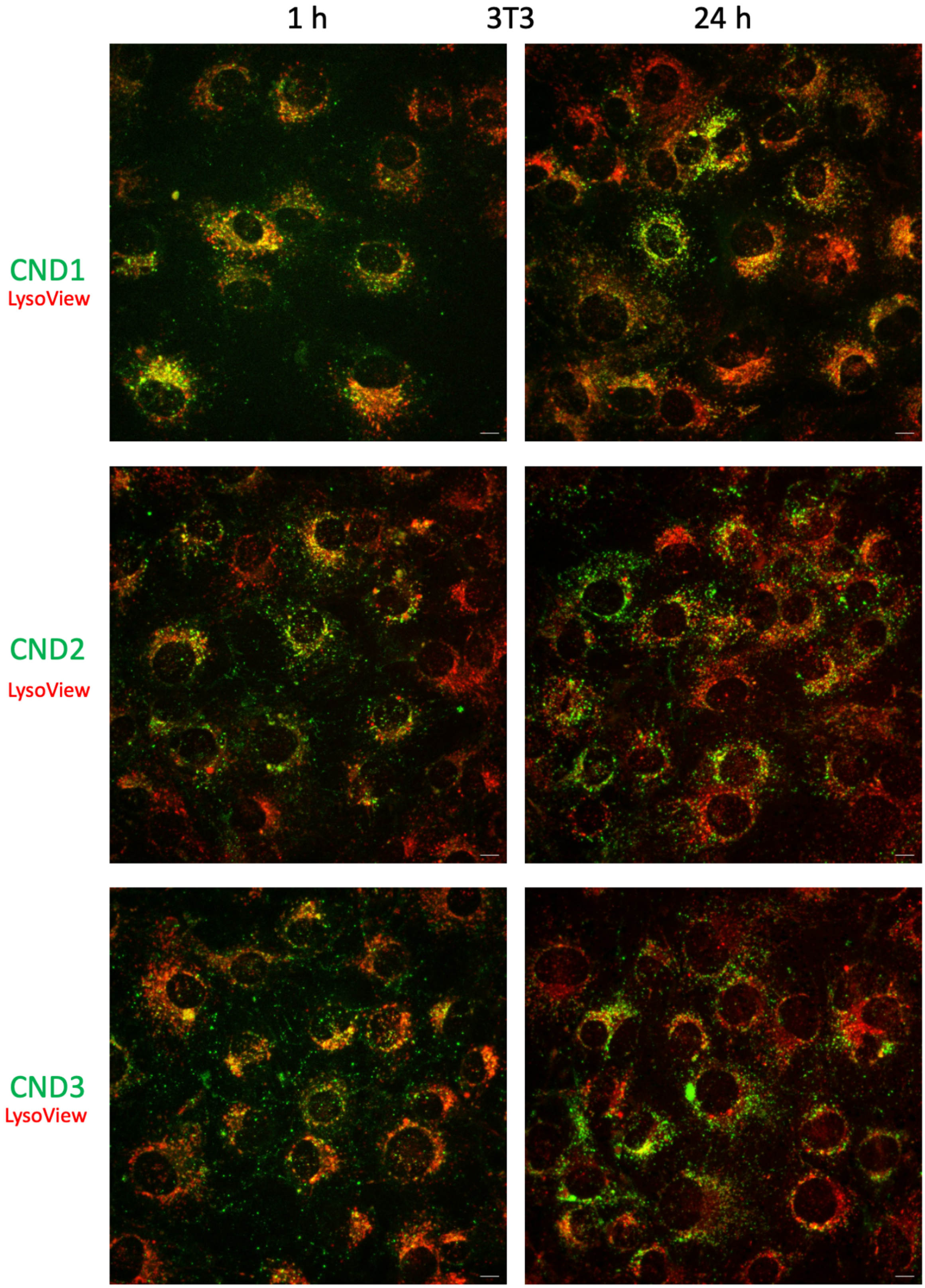
1 h
3T3
24 h
CND1
LysoView
CND2
LysoView
CND3
LysoView

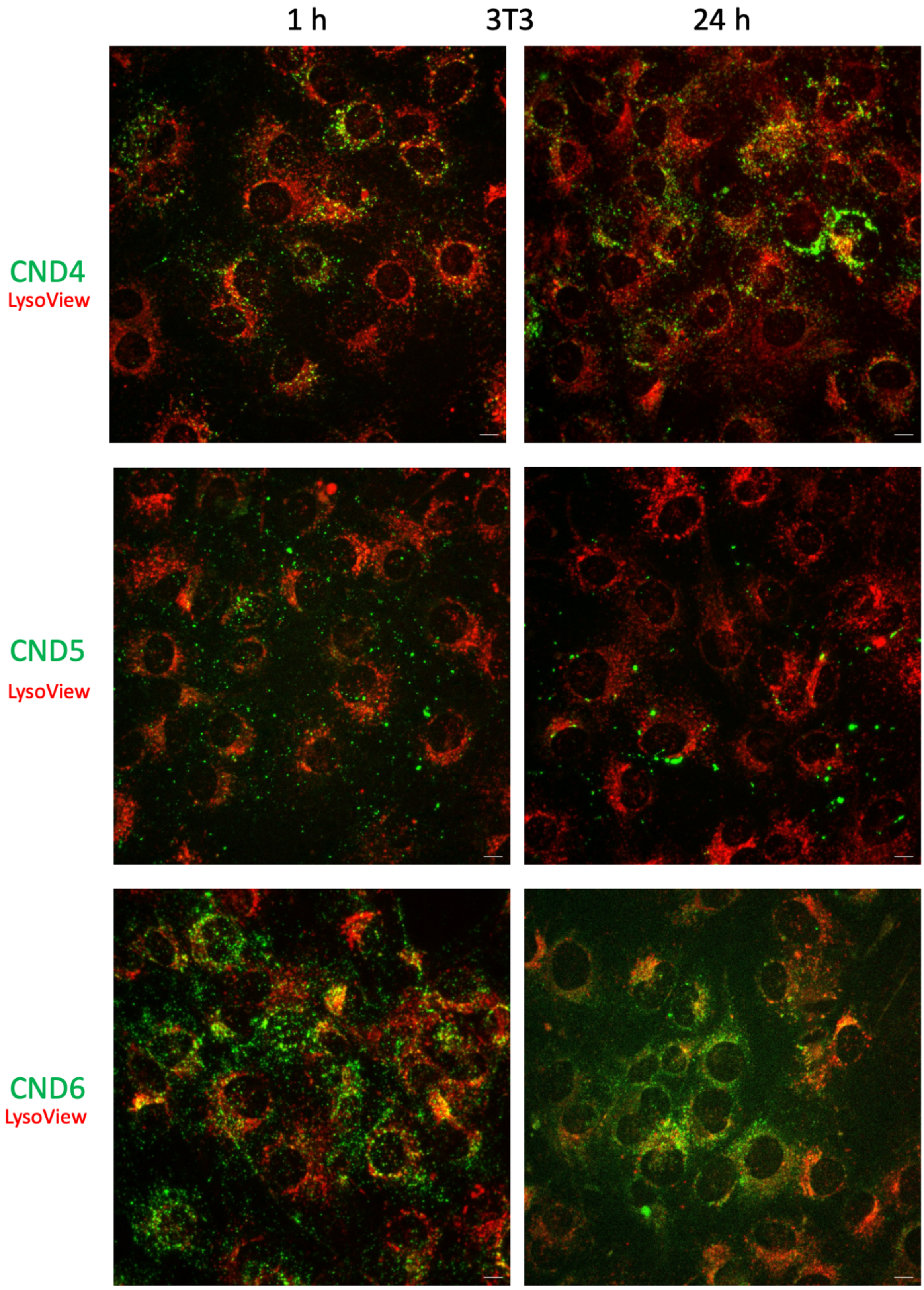

1 h
3T3
24 h
CND4
LysoView
CND5
LysoView
CND6
LysoView

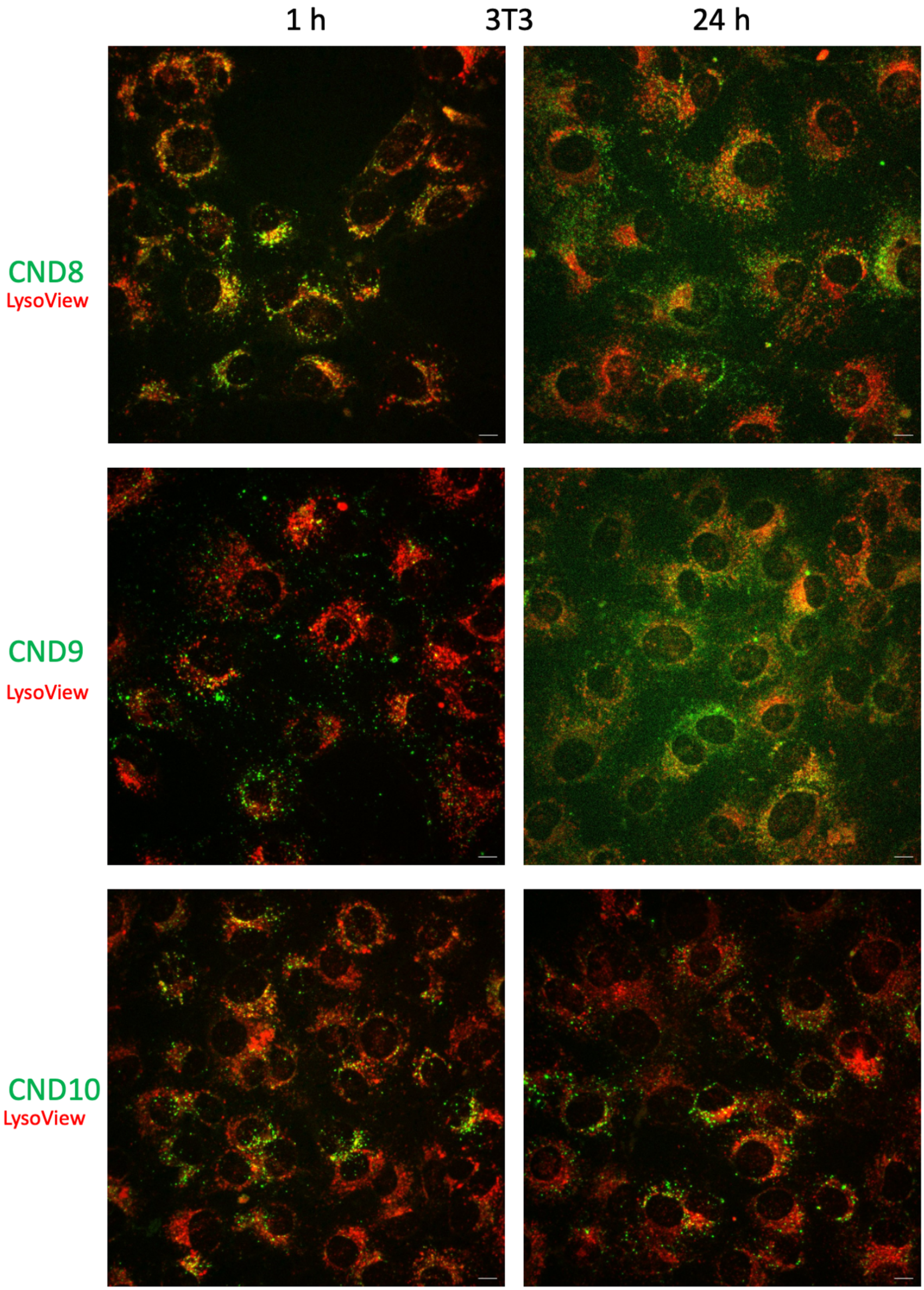
1 h
3T3
24 h
CND8
LysoView
CND9
LysoView
CND10
LysoView

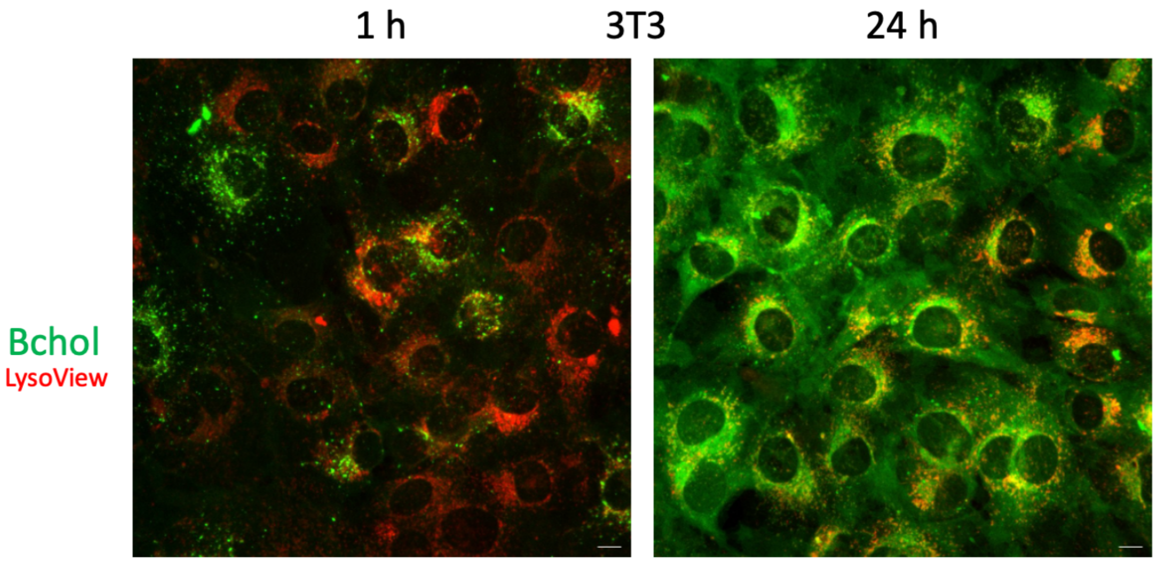


**Fig. S6.** Representative examples of fluorescent confocal composite images of analogs in astrocytes and 3T3 fibroblasts pulse chased for 1h and 24h. Tested probes are depicted in green, LysoView marker is in red. Images in each channel depict average intensity of Z-stack. Bar size – 10 μm.

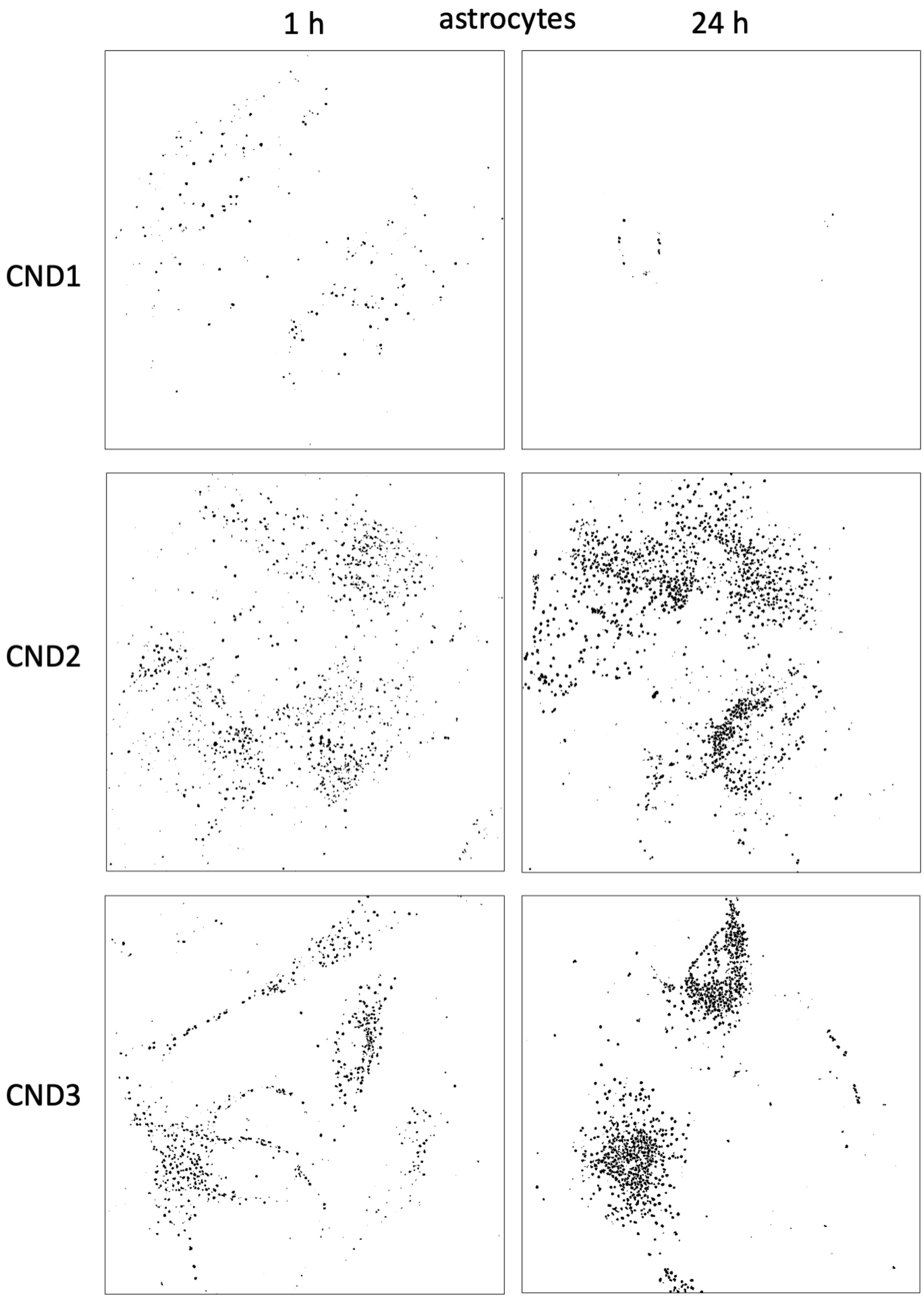
1 h
astrocytes
24 h
CND1
CND2
CND3

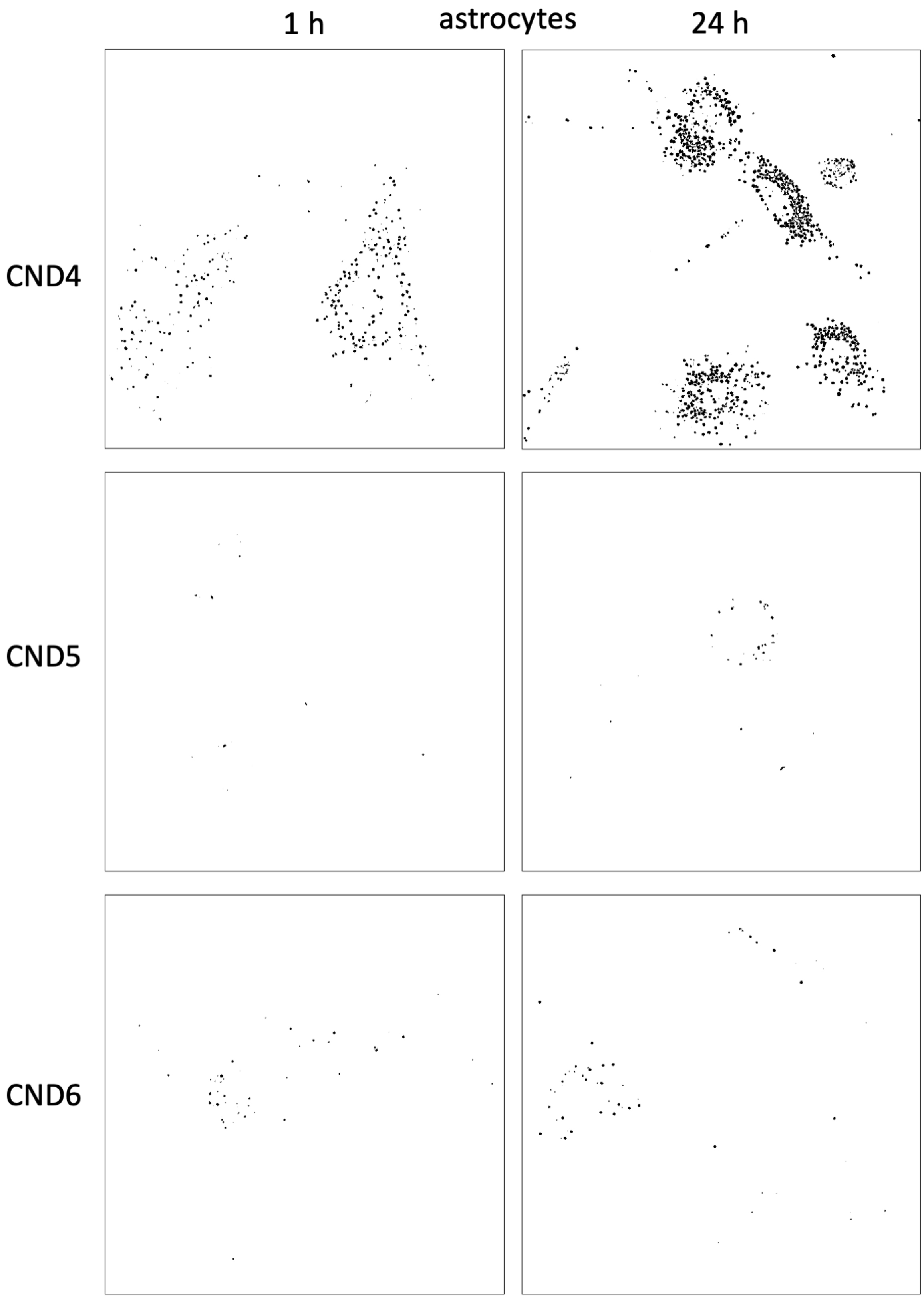
1 h
astrocytes
24 h
CND4
CND5
CND6

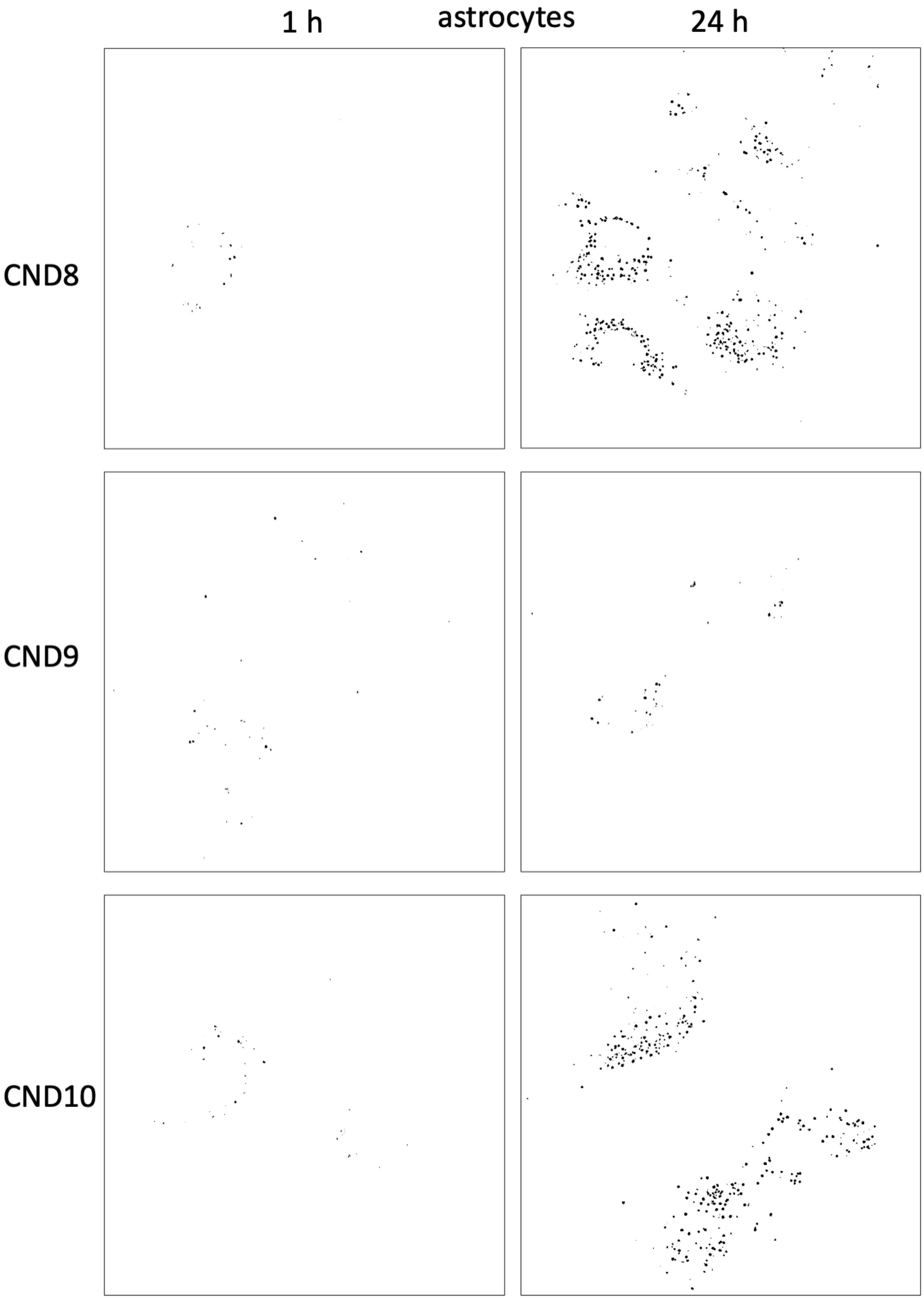
1 h
astrocytes
24 h
CND8
CND9
CND10

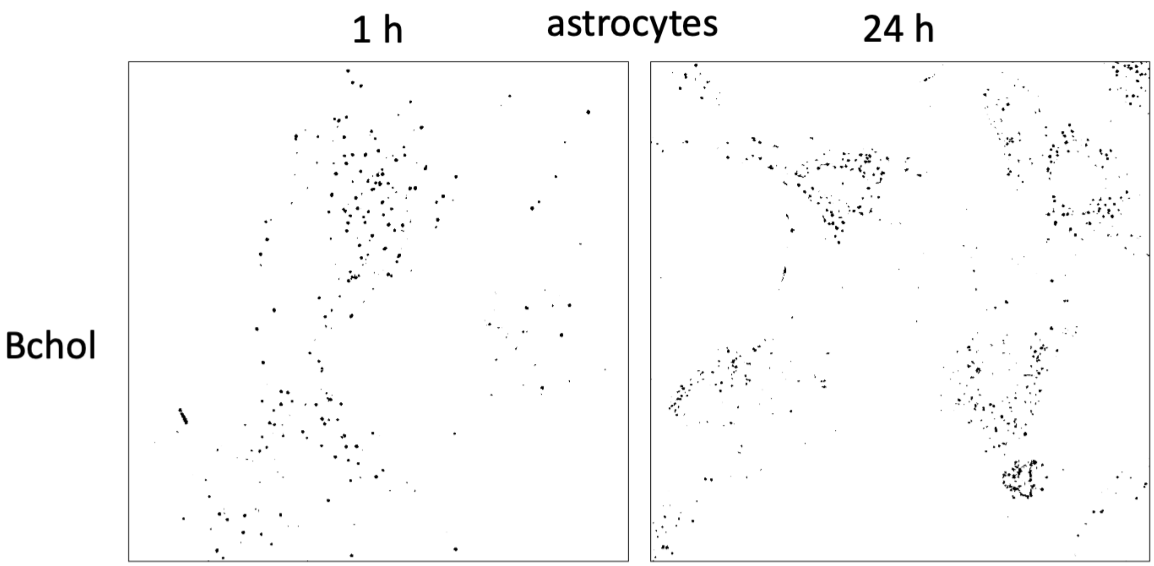
1 h
astrocytes
24 h
Bchol

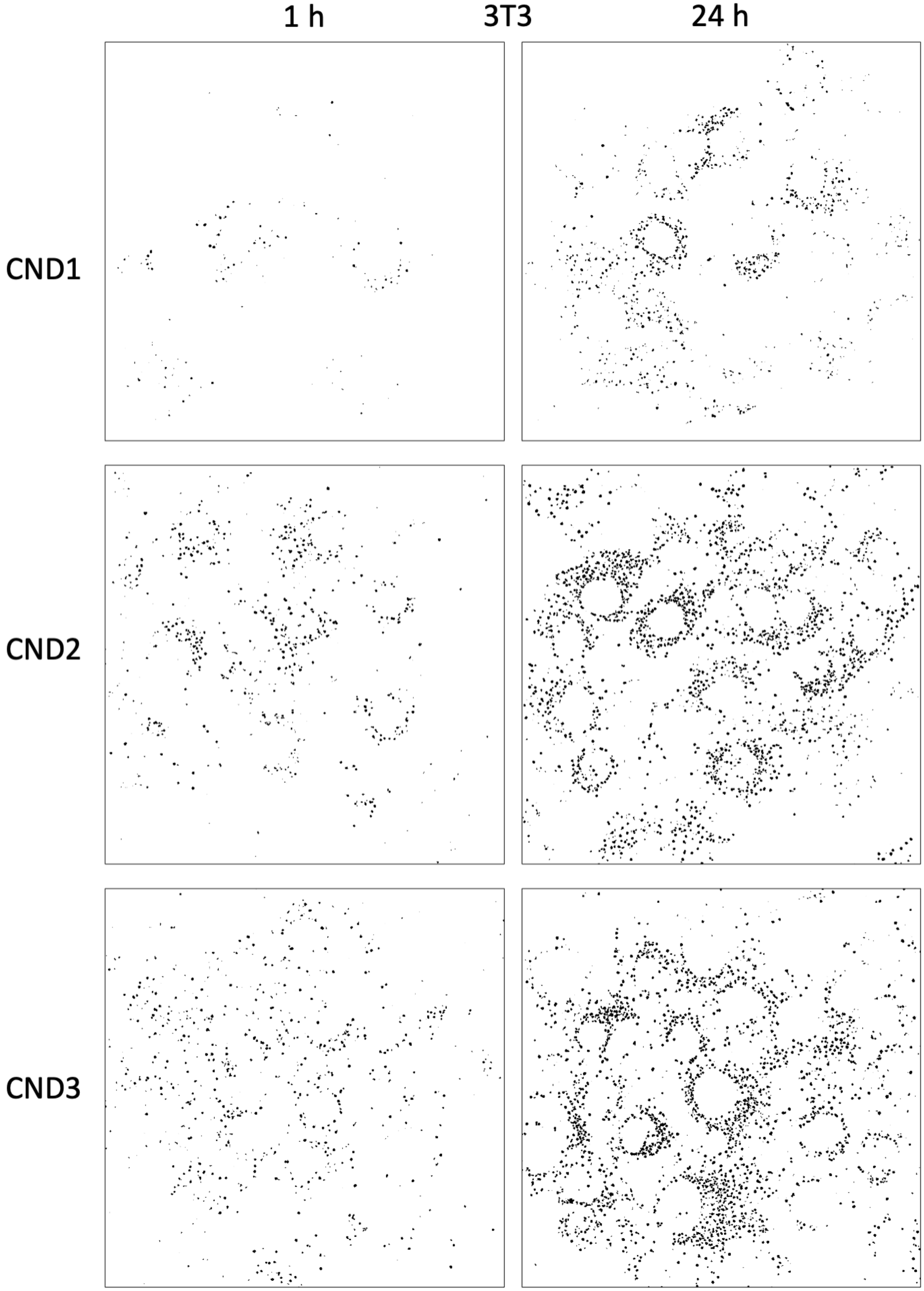
1 h
3T3
24 h
CND1
CND2
CND3

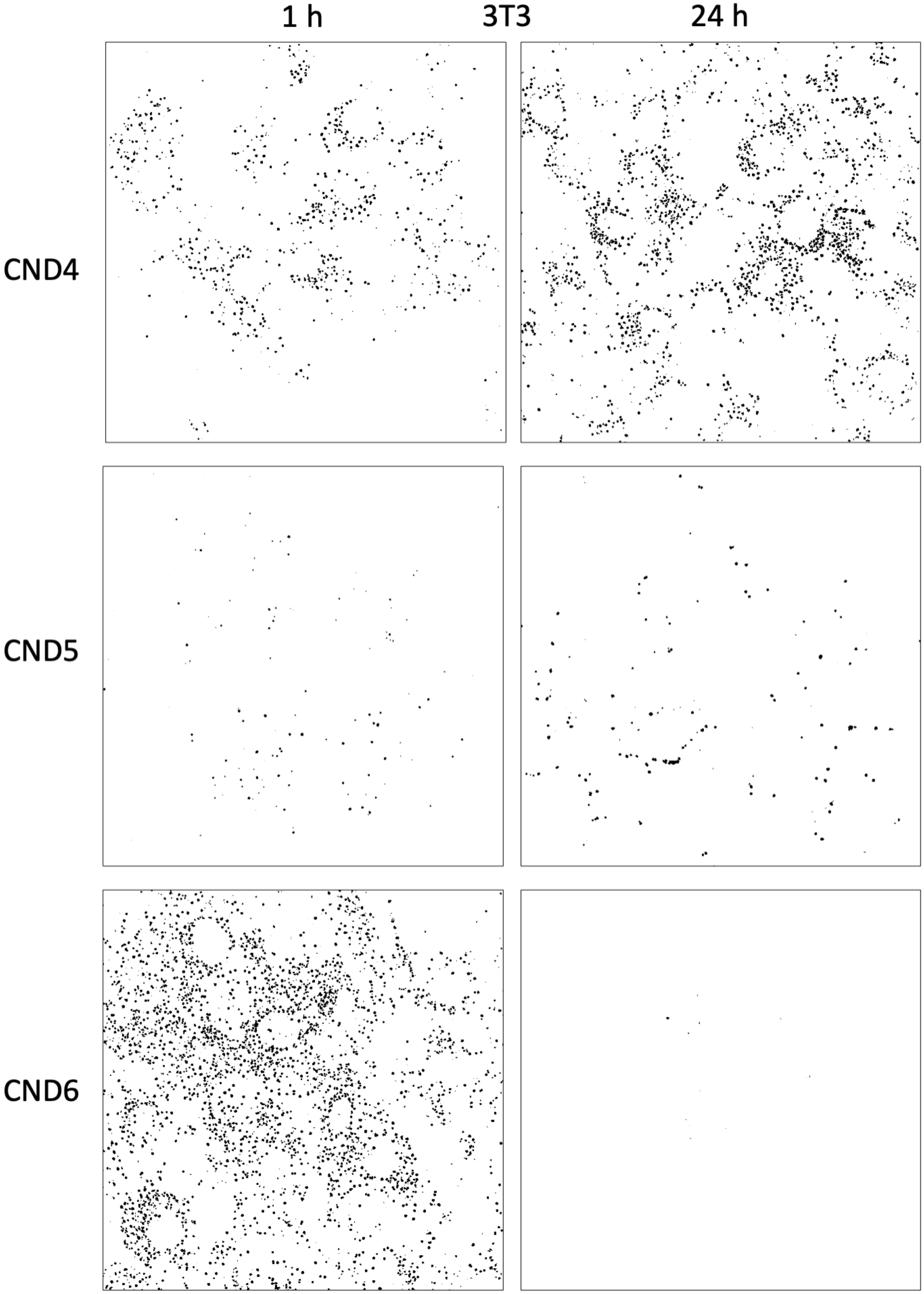

1 h
3T3
24 h
CND4
CND5
CND6

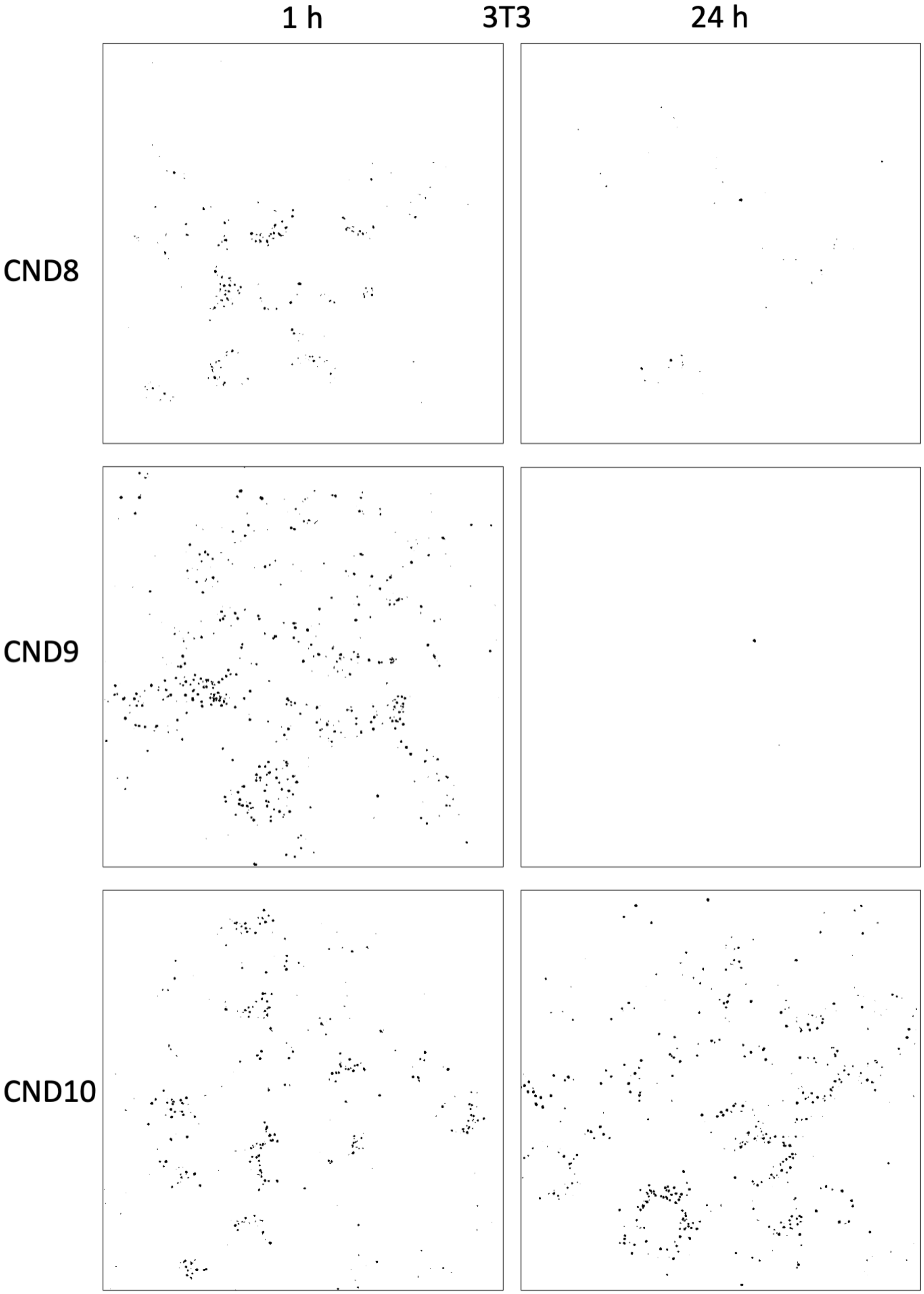
1 h
3T3
24 h
CND8
CND9
CND10

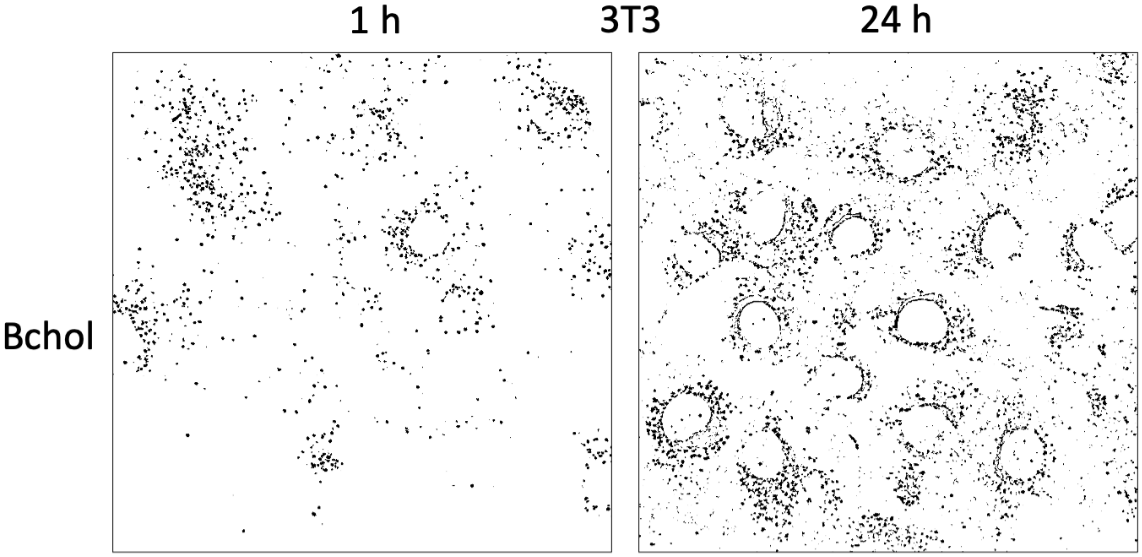


**Fig. S7.** Representative examples of fluorescent puncta masks, obtained through Weka trainable segmentation used to determine particle masks/ROIs. These masks were generated from a single-frame image of the highest intensity profile frame in the green channel in the Z-stack image.

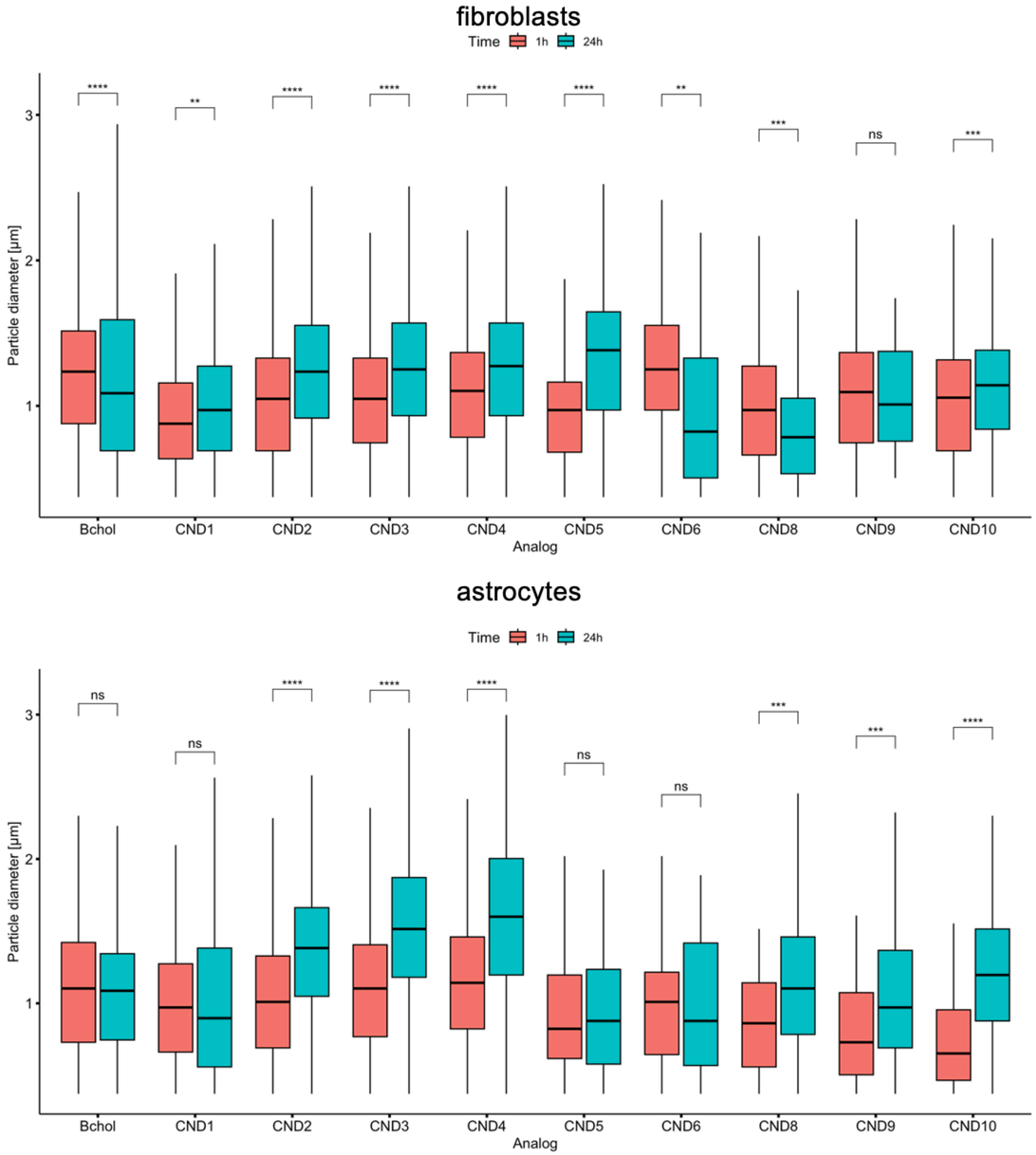


**Fig. S8**. Change in the particle size (diameter) for all tested CND analogs and compared to Bchol at 1h and 24h for both cell lines. Particles were identified in a range between 0-3μm. Statistical analysis was performed using a t-test (rstatix package in R). p-values: **** ($p<0.0001$), *** ($p<0.001$), ** ($p<0.01$), *($p<0.05$), ns – not significant.

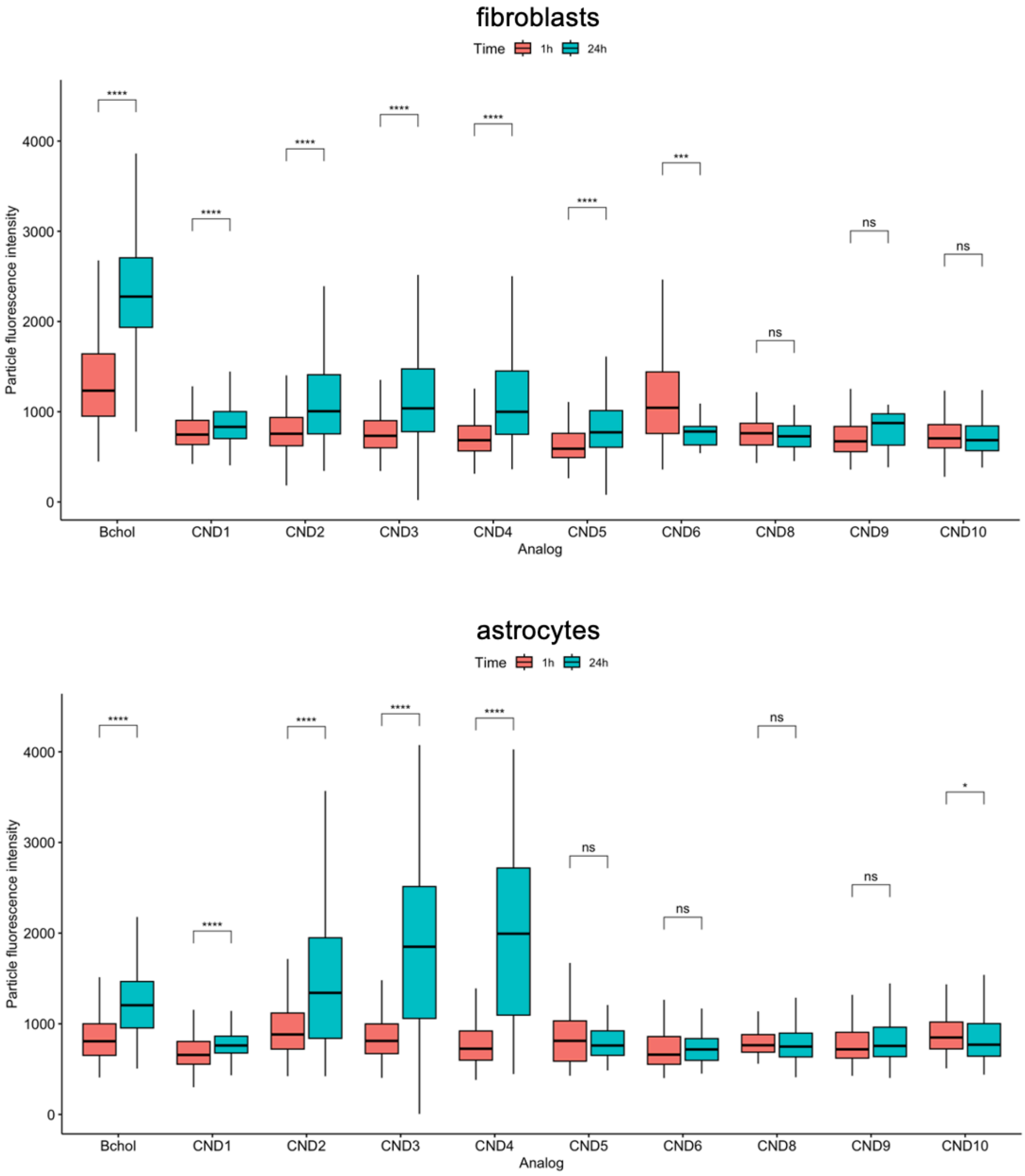


**Fig. S9.** Change in the particle intensity for all tested CND analogs and compared to Bchol at 1h and 24h for both cell lines. Particles were identified in a range between 0-3μm. Statistical analysis was performed using a t-test (rstatix package in R). p-values: **** ($p<0.0001$), *** ($p<0.001$), ** ($p<0.01$), *($p<0.05$), ns – not significant.

### Chemical synthesis of CND probes

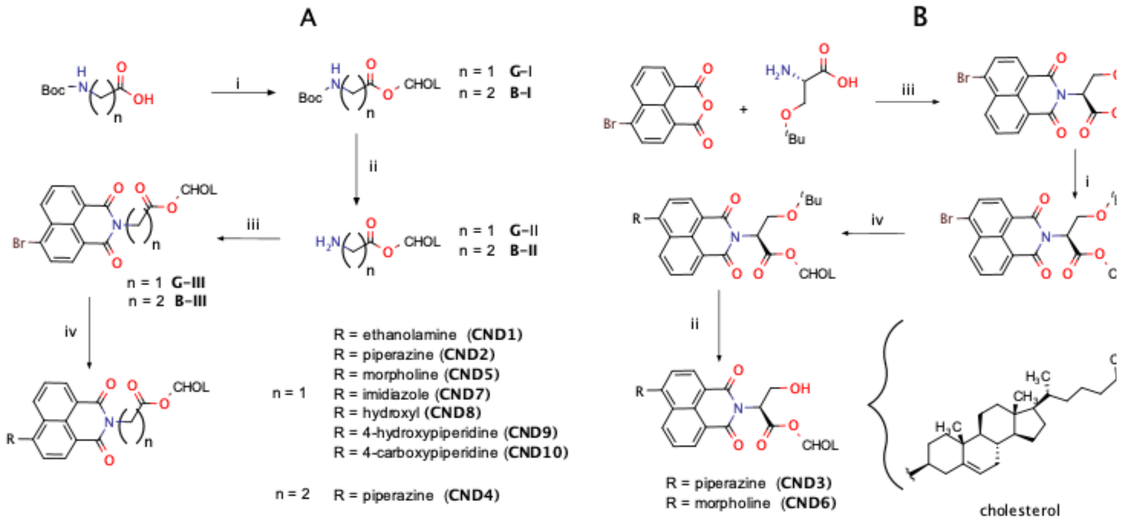


**Scheme S1.** Synthetic scheme for glycine and beta alanine linkers (**A**) and serine linker (**B**). Reagents and conditions: (i) DCC, DMAP, DCM, $N_2$, 0°C, (ii) TFA, TIS, DCM, rt, (iii) 1,8-naphthalic anhydride, EtOH, reflux, (iv) amine (headgroup) DMSO, 100 °C, catalyst (CND7: metformin, CND8:N-hydroxyphthalimide)

### Synthesis of Glycine linker Analogs: CND1, CND2, CND5, CND7 - CND10

Cholesteryl Boc-glycinate and cholesteryl glycinate were synthesized as previously described and this method was adapted for esterification of all carboxylic acid containing intermediates.[1]

(**G-I**) *Cholesteryl Boc-glycinate:* In a round bottom flask, cholesterol (500 mg, 1.30 mmol), boc-glycine (343 mg 1.95 mmol), and DMAP (16 mg 0.130 mmol) were dissolved in anhydrous DCM (10 mL). The solution was stirred and purged with $N_2$ and maintained at 0°C in an ice bath. DCC (430 mg, 2.08 mmol) was dissolved in anhydrous DCM (5 mL) and added to the reaction mixture slowly. The reaction stirred for 30 minutes on the ice bath before being removed and stirring at room temperature overnight. Dicyclohexylurea was filtered off and the filtrate was collected. The filtrate was washed three times with 10% HCl (20 mL) and the organic layer was dried over $Na_2SO_4$ and then was removed by vacuum to give white crystals. Boc-Gly-Chol was purified on silica column in 10% EtOAc in hexanes (1:9 v:v). Solvent was dried over $Na_2SO_4$ and removed by vacuum to obtain 536 mg of a white solid (yield = 76%). $^1$H-NMR (400 MHz $CDCl_3$): δ 5.39 (s, 1H), 5.04 (s, 1H), 4.70 (m, 1H), 3.88 (d, 2H), 2.35 (d, 2H) 0.64 – 2.1 (m, 50H) $^{13}$C-NMR (400 MHz $CDCl_3$): δ 169.7, 155.6, 139.3, 122.9, 79.8, 75.1, 56.6, 56.1, 50.0, 42.6, 42.3, 39.7, 39.5, 38.0, 36.5, 36.1, 35.7, 31.89, 31.85, 28.3, 28.2, 28.0, 27.7, 24.2, 23.8, 22.8, 22.5, 21.0, 19.2, 18.7, 11.8. $C_{34}H_{57}NO_4$ HRMS-ESI: $[M+Na]^+$ $C_{34}H_{57}NO_4$ calcd. m/z = 566.4180 Da, obs. m/z = 566.4188 Da

(**G-II**) *Cholesteryl glycinate:* In a round bottom flask, 536 mg (0.99 mmol) **G-I** was dissolved in 20 mL DCM, 270 µL (1.30 mmol) triisopropylsilane and 3 mL TFA at room temperature. Reaction was stirred for 2 hours before being quenched with 50 mL $H_2O$ and saturated sodium bicarbonate to remove TFA. The organic layer is then evaporated and then purified on silica (MeOH:EtOAC 1:1 v:v) to yield 420 mg of a white solid (yield = 96%). $^1$H-NMR (400 MHz $CDCl_3$): δ 5.40 (s, 1H) 4.69 (m, 1H), 3.41 (s, 2H), 2.35 (d, 2H), 0.64 – 2.1 (m, 41H) $^{13}$C-NMR (400 MHz $CDCl_3$): δ 173.7, 139.4, 122.8, 74.5, 56.6, 56.1, 50.0, 44.2, 42.3, 39.7, 39.5, 38.1, 36.9, 36.5, 36.1, 35.7, 31.9, 31.8, 28.2, 28.0, 27.7, 24.2, 23.8, 22.8, 22.5, 21.0, 19.2, 18.7, 11.8. $C_{29}H_{49}NO_2$ HRMS-ESI: calcd. m/z = 466.3662 Da, obs. m/z = 466.3656

(**G-III**) *4-bromo-1,8-naphthalimide cholesteryl glycinate:* In a round bottom flask, 34mg (0.077 mmol) of **G-II** and 21mg (1 equivalent) of 4-bromo-1,8-naphthalic anhydride was dissolved in ethanol and refluxed overnight. After reaction is complete, ethanol is removed by vacuum. The crude product is purified on silica in (10:10:1 $CHCl_3$:Hexanes:EtOAc) and dried on $Na_2SO_4$. Solvent was removed by vacuum to yield 24 mg of a white solid (yield = 44%). $^1$H-NMR (400 MHz $CDCl_3$): δ 8.66 (d, 1H), 8.58 (d, 1H), 8.42 (d, 1H), 8.06 (d, 1H), 7.86 (t, 1H), 5.39 (s, 1H), 4.92 (s, 2H), 4.73 (m, 1H), 2.4 (d, 2H) 0.64 – 2.1 (m, 41H) $^{13}$C-NMR (400 MHz $CDCl_3$): δ 167.2, 163.2, 139.4, 133.6, 132.3, 131.5, 131.1, 130.7, 129.1, 128.1, 122.8, 122.8, 122.6, 121.83, 75.5, 56.7, 56.1, 50.0, 42.3, 41.6, 39.7, 39.5, 37.9, 36.9, 36.5, 36.1, 35.7, 31.9, 31.8, 28.2. 28.0, 27.7, 24.2, 23.8, 22.8, 22.5, 21.0, 19.3, 18.7, 11.8, $C_{41}H_{52}BrNO_4$ , HRMS-ESI: calcd. m/z = 726.2929 Da, obs. m/z = 726.2929 Da

(**CND1**) *4-ethanolamine-1,8-naphthalimide cholesteryl glycinate:* In a round bottom flask, 58 mg (0.0827 mmol) of **G-III** was dissolved in 2 mL of DMSO. 3 equivalents of neat ethanolamine (15 µL) was added to reaction and was stirred and heated to 100°C. The reaction ran for two hours and was then diluted with 250 mL of water to give < 1% DMSO solution. The aqueous layer was extracted three times with EtOAc (75mL). 35 mg of crude was used for purification on a CombiFlash®Rf+ system in a DCM:MeOH gradient. 100% DCM was allowed to run for three minutes and then the gradient was increased to 5% MeOH over a period of 20 minutes. The gradient ran isocratically at 5% MeOH for an additional 5 minutes before increasing to 100% MeOH over the next 25 minutes. The solvent was evaporated to give an orange solid in a yield of 34 mg (yield = 58%): $^1$H-NMR (400 MHz $CDCl_3$): δ 8.08 (dd, 2H), 7.92 (d, 1H), 7.08 (t, 1H), 6.43 (d, 1H), 6.35 (t, 1H) 5.46 (s, 1H), 4.89 (m, 3H), 3.96 (s, 2H), 3.74 (s, 1H), 3.47 (q, 2H), 2.53 (d, 2H), 0.64 – 2.1 (m, 41H) $^{13}$C-NMR (400 MHz $CDCl_3$): δ 169.4, 164.3, 150.7, 139.5, 134.5, 130.8, 128.9, 127.1, 124.3, 122.9, 121.0, 119.8, 108.3, 104.0, 75.9, 60.6, 56.7, 56.2, 50.1, 46.6, 42.3, 41.4, 39.7, 39.5, 38.1, 37.0, 36.6, 36.2, 35.7, 31.9, 31.8, 28.2, 28.0, 27.7, 24.2, 23.8, 22.7, 22.5, 21.0, 19.3, 18.7, 11.8. $C_{43}H_{58}N_2O_5$ HRMS-ESI: $[M+Na]^+$ calcd. m/z = 705.4238 Da, obs. m/z = 705.4206 Da

(**CND2**) *4-piperazinyl -1,8-naphthalimide cholesteryl glycinate:* In a round bottom flask, 100 mg (0.142 mmol) of **G-III** was suspended in 2 mL of DMSO. 122 mg (1.420 mmol, 10 equiv) of piperazine was added to the reaction mixture and the reaction was heated at 100°C with stirring for three hours. The reaction was cooled to room temperature and diluted with water to give < 1% DMSO solution. The crude product was then extracted with three 50 mL washes of EtOAc. The crude product was purified on a silica column in a gradient of 100% $CHCl_3$ to 5:1 $CHCl_3$:IPA in 1% triethylamine. The pure fractions were washed with 10% HCl to remove the triethylamine. The organic layer was collected and washed with a saturated solution of sodium bicarbonate followed by a wash with deionized water. The organic layer was collected and evaporated to obtain 56 mg of a yellow solid (yield = 55%). $^1$H-NMR (400MHz $CDCl_3$): δ 8.60 (d, 1H), 8.54 (d, 1H), 8.45 (d, 1H), 7.72 (t, 1H), 7.24 (d, 1H), 5.38 (s, 1H), 4.92 (s, 2H), 4.73 (m, 1H), 3.23 (dt, 8H), 2.39 (d, 2H) 0.64 – 2.1 (m, 41H) $^{13}$C-NMR (400 MHz $CDCl_3$): δ 167.6, 164.1, 163.6, 156.7,139.5, 132.9, 131.4, 130.7, 130.1, 126.2, 125.6, 122.8, 122.7, 116.1, 114.9, 75.2, 56.69, 56.1, 54.3, 49.9, 46.1, 42.3, 41.5, 39.7, 39.5, 37.9, 36.9, 36.5, 36.1, 35.7, 31.9, 31.8, 28.2, 28.0, 27.7, 24.2, 23.8, 22.8, 22.5, 21.0, 19.3, 18.7, 11.8. $C_{45}H_{61}N_3O_4$ HRMS-ESI: $[M+H]^+$ calcd. m/z = 708.4735 Da, obs. m/z = 708.4708 Da

(**CND5**) *4-morpholine-1,8-naphthalimide cholesteryl glycinate*: In a round bottom flask, 50 mg of **G-III** was dissolved in 2 mL of DMSO, stirred and heated to 100°C. To the heated solution, 62 μL (0.713 mmol, 10 equiv) of morpholine was added to the reaction and stirred for 4 hours. After the reaction, the solution was cooled to room temperature and diluted with 200mL of water. The aqueous layer was extracted with EtOAc (3x50mL) and collected. The organic layer was then evaporated to obtain a crude yellow solid. The crude product was purified in 20:1 DCM:MeOH on a silica column to yield a bright yellow solid 33 mg (yield = 65%). $^1$H-NMR (400 MHz $CDCl_3$): δ 8.63 (d, 1H), 8.56 (d, 1H), 8.47 (d, 1H), 7.73 (t, 1H), 7.27 (d, 1H), 5.39 (d, 1H), 4.9 (s, 2H), 4.73 (m, 1H), 4.04 (t, 4H), 3.30 (t, 4H), 2.40- 0.69 (m, 43H). $^{13}$C-NMR (400 MHz $CDCl_3$): δ 167.5, 164.0, 163.5, 155.9, 139.5, 132.9, 131.5, 130.4, 130.1, 126.2, 125.8, 122.9, 122.7, 116.7, 114.9, 75.3, 66.9, 56.7, 56.1, 53.4, 50.0, 42.3, 41.5, 39.7, 39.5, 37.9, 36.9, 36.5, 36.1, 35.7, 31.9, 31.8, 28.2, 28.0, 27.7, 24.2, 23.8, 22.8, 22.5, 21.0, 19.3, 18.7, 11.8. $C_{45}H_{60}N_2O_5$ HRMS-ESI: $[M+Na]^+$ calcd. m/z = 731.4394 Da, obs. m/z = 731.4361 Da

(**CND7**) *4-imidazole-1,8-naphthalimide-cholesteryl-glycinate*: 50 mg of **G-III**, 10mg of imidazole, 42 mg of $Cs_2CO_3$, 1.6 mg of metformin, and 4.9 mg of CuI were suspended in 1 mL DMF. The suspended reaction solution was stirred and heated to 110 °C for 24 hours. After the 24 hours, the reaction was cooled to room temperature and diluted in 100 mL of water and acidified. The aqueous solution was extracted with EtOAc (3x50mL) and the organic layer was collected and evaporated. The crude solid was purified in a gradient of 100% $CHCl_3$ to 99:1 $CHCl_3$:TEA to yield 26 mg of an off white/yellow solid (yield = 53%). $^1$H-NMR (400 MHz $CDCl_3$): δ 8.73 (m, 2H), 8.13 (d, 1H), 7.88 (m, 2H), 7.77 (d, 1H), 7.40 (d 2H), 5.40 (s, 1H), 4.96 (s, 2H), 4.75 (m, 1H), 2.42 – 0.69 (m, 43H). $^{13}$C-NMR (400 MHz $CDCl_3$): δ 167.2, 163.4, 162.9, 139.4, 139.4, 132.5, 131.4, 130.6, 129.3, 129.2, 128.4, 127.6, 124.1, 122.9, 122.7, 122.6, 75.6, 56.6, 56.1, 49.9, 42.3, 41.7, 39.7, 39.5, 37.9, 36.9, 36.5, 36.1, 35.7, 31.9, 31.8, 28.2, 28.0, 27.7, 24.2, 23.8, 22.8, 22.5, 21.0, 19.3, 18.7, 11.8. $C_{44}H_{55}N_3O_4$ HRMS-ESI: $[M+H]^+$ calcd. m/z = 690.4252 Da, obs. m/z = 690.4265 Da

(**CND8**) *4-hydroxy-1,8-naphthalimide-cholesteryl-glycinate*: 50 mg of **G-III** (0.071 mmol), 58 mg of *N*-Hydroxyphthalimide (5 equivalents), 59 mg of $K_2CO_3$ (5 equivalents) were suspended in 2 mL DMSO. The reaction was heated to 100 °C and spun for 3 hours. After the 3 hours, the reaction was cooled to room temperature, diluted with

200 mL water, and acidified with 10% HCl. The aqueous layer was extracted with EtOAc (3x25mL) and then it was evaporated. The crude solid was purified on a silica column in a gradient of 100% $CHCl_3$ to 9:1 $CHCl_3$:EtOAc to yield 29 mg of a yellow solid (yield = 64% ). $^1$H-NMR (400 MHz $CDCl_3$): δ 8.46, (d, 1H), 7.98 (d, 1H), 7.94 (d, 1H), 7.38 (t, 1H), 6.72 (d, 1H), 5.47 (s, 1H), 4.99 (s, 1H), 4.89 (m, 1H), 2.54 (d, 2H), 0.71 – 2.06 (m, 42H). $^{13}$C-NMR (400 MHz $CDCl_3$): δ 170.8, 164.3, 163.3, 158.9, 139.2, 134.0, 131.5, 129.0, 125.4, 123.2, 122.0, 121.1, 113.3, 109.8, 56.7, 56.1, 50.0, 42.3, 41.4, 39.7, 39.5, 37.9, 36.9, 36.6, 36.1, 35.8, 31.9, 31.8, 28.2, 28.0, 27.7, 24.3, 22.8, 22.5, 21.0, 19.3, 18.7, 11.8. $C_{41}H_{53}NO_5$ HRMS-ESI: $[M+Na]^+$ calcd. m/z = 662.3816 Da, obs. m/z = 662.3783 Da

(**CND9**) *4-(4-hydroxy-piperadinyl)-1,8-naphthalimide-cholesteryl-glycinate*: 50 mg of **G-III** (0.071 mmol) and 36 mg of 4-hydroxypiperadine (5 equivalents) was dissolved in 1 mL DMSO, stirred, and heated to 100 °C for four hours. After the four hours, the reaction was cooled to room temperature and diluted with 100 mL of water. The aqueous layer was then extracted with EtOAc (3x25mL) and the organic layer was collected and evaporated. The crude solid was then purified on silica column in a gradient of 1:1 $CHCl_3$:hexanes to 9:1 $CHCl_3$:EtOAc to yield 42 mg of a yellow/orange solid (yield = 82%). $^1$H-NMR (400 MHz $CDCl_3$): δ 8.60 (d, 1H), 8.52 (d, 1H), 8.41 (d, 1H), 7.72 (t, 1H), 7.24 (d, 1H), 5.38 (s, 1H), 4.92 (s, 2H), 4.73 (m, 1H), 4.04 (m, 1H), 3.53 (m, 2H), 3.10 (t, 2H), 0.68-2.39 (m, 48H). $^{13}$C-NMR (400 MHz $CDCl_3$): δ 167.6, 164.2, 163.6, 156.7, 139.5, 132.9, 131.4, 130.7, 130.1, 126.3, 125.6, 122.8, 122.7, 115.9, 115.0, 75.2, 67.3, 56.6, 56.1, 50.7, 49.9, 42.3, 41.5, 39.7, 39.5, 37.9, 36.9, 36.5, 36.1, 35.7, 34.6, 31.9, 31.8, 28.2, 28.0, 27.7, 24.2, 23.8, 22.8, 22.5, 21.0, 19.3, 18.7, 11.8. $C_{46}H_{62}N_2O_5$ HRMS-ESI: $[M+H]^+$ calcd. m/z = 723.4731 Da, obs. m/z = 723.4707 Da

(**CND10**) *4-(4-carboxy-piperadinyl)-1,8-naphthalmide-choleseryl-glycinate*: 100 mg of **G-III** (0.142 mmol), 100 mg of tert-Butyl piperadine-4-carboxylate (3.8 equiv), was suspended in 3 mL of DMSO. The reaction was heated to 110 °C and ran for four hours and then cooled to room temperature. The reaction mixture was then diluted in 300 mL water and the aqueous layer was acidified. The aqueous layer was extracted with EtOAc (3x50 mL) and evaporated. The crude yellow solid was solubilized in 6 mL DCM and 80 µL TIS. The reaction mixture was spun at room temperature and then 2 mL of TFA was added to the reaction. The reaction ran for 2 hours and then the reaction was quenched with 30 mL $H_2O$ and TFA was neutralized with saturated $NaHCO_3$. The reaction was then acidified with 10% HCl and then extracted with EtOAc (3x30mL). The organic layer was collected and evaporated to obtain a crude yellow solid. 64 mg of the crude yellow solid was purified in a gradient of 50:1 $CHCl_3$:IPA to 20:1 $CHCl_3$:IPA to obtain 36 mg of yellow solid. (yield = 56%). $^1$H-NMR (400 MHz $CDCl_3$): δ 8.62 (d, 1H), 8.54 (d, 1H), 8.42 (d, 1H), 7.74 (t, 1H), 7.25 (d, 1H), 5.38 (s, 1H), 4.93 (s, 2H), 4.74 (m, 1H), 3.61 (d, 2H), 3.04 (t, 2H), 0.69 – 2.68 (m, 48H). $^{13}$C-NMR (400 MHz $CDCl_3$): δ 179.3, 167.6, 164.1, 163.6, 156.7, 139.5, 132.9, 131.5, 130.6, 130.1, 126.4, 125.7, 122.8, 122.7, 116.2, 115.1, 75.3, 56.6, 56.1, 52.8, 49.9, 42.3, 41.5, 40.4, 39.7, 39.5, 37.9, 36.9, 36.5, 36.1, 35.8, 31.9, 31.8, 28.3, 28.0, 27.7, 24.2, 23.8, 22.8, 22.5, 21.0, 19.3, 18.7, 11.8. $C_{47}H_{62}N_2O_6$ HRMS-ESI: $[M+Na]^+$ calcd. m/z = 773.4500 Da, obs. m/z = 773.4467 Da

**Synthesis of β-alanine Linker Analog: CND4**

(**B-I**) *Cholesteryl-(Boc-β-alanine) ester:* In a round bottom flask, 100mg of cholesterol (0.259 mmol), 73.5 mg of *Boc-β-alanine* (0.388 mmol, 1.5 equivalents), and 5 mg of DMAP (0.0259 mmol) were dissolved in anhydrous DCM (10 mL). The solution was stirred and purged with $N_2$ and maintained at 0°C in an ice bath. 85 mg of DCC (0.414 mmol ) was dissolved in 2 mL of anhydrous DCM and added to the reaction mixture slowly. The reaction was allowed to stir for 30 minutes on the ice bath before being removed and stirring at room temperature overnight. Dicyclohexylurea was filtered off and the filtrate was collected. The filtrate was washed three times with 20 mL of 10% HCl and the organic layer was dried over $Na_2SO_4$ and then was evaporated to give 86 mg of white crystals (yield = 60%) $^1$H-NMR (400 MHz $CDCl_3$): δ 5.40 (s, 1H), 5.02 (s, 1H), 4.66 (m, 1H), 3.40 (q, 2H), 2.51 (t, 2H), 2.34 (d, 2H), 2.01 – 0.69 (m, 50H) $^{13}$C-NMR (400 MHz $CDCl_3$): δ 171.9, 155.7, 139.5, 122.7, 79.3, 74.3, 56.7, 56.1, 50.0, 42.3, 39.7, 39.5, 38.1, 36.9, 36.6, 36.1, 35.7, 34.9, 31.9, 31.8, 28.4, 28.2, 28.0, 27.7, 24.2, 23.8, 22.8, 22.5. 21.0, 19.3, 18.7, 11.8. $C_{35}H_{59}NO_4$ HRMS-ESI: $[M+Na]^+$ calcd. m/z = 580.4336 Da, obs. m/z = 580.4347 Da

(**B-II**) *Cholesteryl-β-alanine ester:* In a round bottom flask, 86 mg **B-I** was dissolved in 10 mL DCM, and 3 mL TFA at room temperature. Reaction was stirred for 2 hours before being quenched with 50mL $H_2O$ and saturated sodium bicarbonate to neutralize TFA. The product is then extracted with EtOAc and then organic layer is collected, and the organic solvent is evaporated. Attempts to purify βAla-Chol and NMR experiments fail due to the hydrolysis of the β-Alanine cholesterol ester to β-Alanine and cholesterol. After extraction step, the crude product is sufficient to perform the next reaction. The crude product is a wax-like substance. $C_{30}H_{51}NO_2$ HRMS-ESI: $[M+H]^+$ calcd. m/z = 458.3993 Da, obs. m/z = 458.4013 Da

(**B--III**) *4-bromo-1,8-naphthalimide-cholesteryl-β-alanine*: In a round bottom flask, **B-II** and 72 mg (1 equivalent) of 4-bromo-1,8-naphthalic anhydride was dissolved in ethanol and refluxed overnight. After reaction is complete, ethanol is removed by vacuum. The solid was purified on silica gel on 10:10:1 $CHCl_3$:Hx:EtOAc (v:v:v) to obtain 117 mg of a white solid (yield = 63%) $^1$H-NMR (400MHz $CDCl_3$): δ 8.68, (d, 1H), 8.59 (d, 1H), 8.43 (d, 1H), 8.06 (d, 1H), 7.86 (t, 1H), 5.36 (s, 1H), 4.63 (m, 1H), 4.50 (t, 2H), 2.76 (t, 2H), 0.50-2.50 (m, 43H). $^{13}$C-NMR (400Mhz $CDCl_3$): δ 170.5, 163.3, 139.6, 133.4, 132.1, 131.3, 131.1, 130.6, 130.4, 129.0, 128.0, 122.9, 122.6, 122.1, 74.3, 56.6, 56.1, 50.0, 42.3, 39.7, 39.5, 38.0, 36.9, 36.5, 36.3, 36.1, 35.7, 32.9, 31.9, 31.8, 28.2, 28.0, 27.6, 24.2, 23.8, 22.8, 22.5, 21.0, 19.2, 18.7, 11.8. $C_{42}H_{54}BrNO_4$ HRMS-ESI: $[M+Na]^+$ calcd. m/z = 740.3117 Da obs. m/z = 740.3097 Da

(**CND4**) *4-piperizinyl-1,8-naphthalimide-cholesteryl-β-alanate* 117mg of **B-III** was dissolved in 4mL of DMSO and 100mg of piperazine (7 equivalents, 1.148mmol), was heated to 100° C for 1 hour. After 1 hour the reaction was diluted in $H_2O$ and extracted with DCM. DCM was evaporated and the crude was purified on silica gel in 7:3 DCM:IPA 0.2% triethylamine. The organic layer was washed with 10% HCl to remove triethylamine. The organic layer was then washed with saturated sodium bicarbonate, followed by a wash with deionized $H_2O$ to obtain 50 mg of a yellow solid (yield = 42%). $^1$H-NMR (400MHz $CDCl_3$): δ 8.61 (d,1H), 8.55 (d, 1H), 8.44 (d, 1H), 7.71 (t, 1H), 7.25 (d, 1H), 5.36 (s, 1H),

4.64 (m, 1H), 4.49 (t, 2H), 3.25 (dt, 8H), 2.75 (t, 2H), 2.3 – 0.5 (m, 43H). $^{13}$C-NMR (400Mhz $CDCl_3$): δ 170.7, 164.2, 163.7, 156.3, 139.7, 132.6, 131.2, 130.3, 129.9, 126.2, 125.7, 123.1, 122.5, 116.6, 115.0, 74.2, 56.6, 56.1, 54.1, 50.0, 46.0, 42.3, 39.7, 39.5, 38.0, 36.9, 36.5, 36.1, 36.0, 35.7, 33.0, 31.9, 31.8, 28.2, 28.0, 27.6, 24.2, 23.8, 22.8, 22.5, 21.0, 19.2, 18.7, 11.8. $C_{46}H_{63}N_3O_4$ HRMS-ESI: $[M+H]^+$ calcd. m/z = 722.4891 Da, obs. m/z = 722.4878 Da

**Synthesis of Serine Linker Analogs: CND3 and CND6**

(**S-I**) *4-bromo-1,8-naphthalimide-(O-tertbutyl)-L-serine:* In a round bottom flask, 100 mg (0.361 mmol) of 4-bromo-1,8-naphthalic anhydride and 70 mg (1.2 equivalents, 0.433mmol) of O-tert-butyl-L-Serine was dissolved in 10mL of ethanol refluxed overnight. The reaction solution was concentrated on the rotavapor and then diluted in water and acidified (pH = 1). The aqueous solution was then extracted with EtOAc (3x 50mL) and the organic layer was collected and evaporated. The crude brown solid was purified on a silica column in 20:1 DCM:MeOH (v:v) and dried on sodium sulfate to yield 130 mg of a white solid (yield = 86%). $^1$H-NMR (400MHz $CDCl_3$): δ 8.69 (d, 1H), 8.63 (d, 1H), 8.45 (d, 1H) 8.08 (d, 1H), 7.88 (t, 1H), 5.92 (t, 1H), 4.36 (t, 1H), 3.88 (t, 1H), 1.25 (s, 9H) $^{13}$C-NMR (400MHz $CDCl_3$): δ 169.8, 163.2, 133.7, 132.5, 131.7, 131.2, 130.8, 130.7, 129.2, 128.1, 122.6, 121.7, 75.6, 70.5, 59.1, 52.5, 29.6, 27.3. $C_{19}H_{18}BrNO_5$ HRMS-ESI: $[M+Na]^+$ calcd. m/z = 444.0243 Da obs. m/z = 444.0263 Da

(**S-II**) *4-bromo-1,8-naphthalimide-cholesteryl-O-terbutyl-L-serine:* In a round bottom flask, 210 mg of cholesterol (0.542 mmol, 1.5 equivalents) 5 mg of DMAP (0.1 equivalents, 0.036 mmol) and **S-I** were dissolved in 10 mL of anhydrous DCM at 0° C under an $N_2$ atmosphere. Subsequently, 120 mg of DCC (1.6 equivalents, .577mmol) dissolved in 3mL of anhydrous DCM was added to the reaction. The reaction proceeded overnight and dicyclohexaurea was filtered off and washed with DCM while the filtrate was collected. The filtrate was extracted with 25 mL of 10% HCl and then subsequently by deionized $H_2O$. The organic layer was evaporated and then was purified on silica gel in 10:10:1 $CHCl_3$:hexanes:EtOAc (v:v:v) to obtain 180 mg of white crystals (yield = 63%). $^1$H-NMR (400 MHz $CDCl_3$): δ 8.69 (d, 1H), 8.62 (d, 1H), 8.45 (d, 1H), 8.08 (d, 1H), 7.88 (t, 1H), 5.90 (q, 1H), 5.38 (s, 1H), 4.75 (m, 1H), 4.20 (q, 1H), 4.07 (t, 1H), 0.6 - 2.5 (m, 52H). $^{13}$C-NMR (400 MHz $CDCl_3$): δ 167.8, 163.2, 139.5, 133.4, 132.3, 131.4, 131.1, 130.7, 130.4, 129.2, 128.1, 122.9, 122.7, 122.0, 75.0, 75.0, 73.3, 59.4, 56.6, 56.1, 54.2, 49.9, 42.3, 39.7, 39.5, 37.9, 37.8, 36.9, 36.8, 36.5, 36.1, 35.7, 31.8, 31.8, 28.2, 28.0, 27.5, 27.4, 24.2, 23.8, 22.8, 22.5, 21.0, 19.2, 18.7, 11.8. $C_{46}H_{62}BrNO_5$ HRMS-ESI: $[M+Na]^+$ calcd. m/z = 812.3694 Da, obs. m/z = 812.3709 Da

*(***CND3***) 4-piperazynl-1,8-naphthalimide-Cholesteryl-L-serinate:* 180 mg of **S-II** was suspended in 8 mL of DMSO and was stirred with 138 mg of piperazine (1.603mmol, 7 equivalents) and heated at 100°C for 1 hour. After 1 hour, the orange reaction solution was diluted in $H_2O$ ( < 1% DMSO) and extracted with DCM (3x100mL). The organic layer was collected and evaporated to obtain the crude 164 mg of yellow/orange solid. 164 mg of crude product was dissolved in 10mL DCM and 70 μL of triisopropylsilane (TIS) and stirred. To the stirred solution, 2mL of TFA was added and stirred at room temperature for 1 hour. After 1 hour, the reaction was diluted in 50mL $H_2O$ and TFA was quenched

with a solution of saturated sodium bicarbonate. The organic layer was collected and evaporated. The crude product was purified on a silica column in 7:3 DCM:IPA 0.2% triethylamine. The organic layer was washed with 10% HCl to remove triethylamine. The organic layer was then washed with saturated sodium bicarbonate, followed by a wash with deionized $H_2O$ to obtain 42 mg of an orange/yellow solid (yield = 25% ). $^{1}$H-NMR (400MHz $CDCl_3$): δ 8.61 (d, 1H), 8.53 (d, 1H), 8.45 (d, 1H), 7.73 (t, 1H), 7.23 (d, 1H), 5.79 (t, 1H), 5.39 (s, 1H) 4.78 (m, 1H), 4.42 (q, 1H), 4.00 (q, 1H), 3.66 (s, 1H), 2.40- 0.69 (m, 43H).$^{13}$C-NMR (400Mhz $CDCl_3$): δ 169.1, 164.2, 163.7, 156.8, 139.4, 133.2, 131.6, 130.8, 130.1, 126.2, 125.7, 122.8, 116.0, 115.0, 75.2, 70.5, 61.1, 56.6, 56.1, 54.5, 54.3, 49.9, 46.1, 42.3, 39.7, 39.5, 37.9, 37.8, 36.9, 36.8, 36.5, 36.1, 35.7, 31.8, 31.8, 28.2, 28.0, 27.6, 27.5, 24.2, 23.8, 22.8, 22.5, 21.0, 19.2, 18.7, 11.8. $C_{46}H_{63}N_3O_5$ HRMS-ESI: $[M+H]^+$ calcd. m/z = 738.484 Da, obs. m/z = 738.4837 Da

(**CND6**) *4-Morpholinyl-1,8-napthalimide Cholesteryl Serinate*: In a round bottom flask, 34 mg of **S-II** was suspended in 1 mL of DMSO. To the reaction mixture, 75 μL (20 equivalents, 0.0864 mmol) of morpholine was added and the reaction mixture was heated to 100°C for 3 hours. After the 3 hours, the reaction was cooled to room temperature and diluted in 100 mL of $H_2O$ and acidified with 10% HCl in water (v:v). The aqueous layer was then extracted with EtOAc (3x25mL) and collected. The crude yellow solid was resolubilized in 5 mL DCM and stirred. To the stirred solution, 1 mL TFA and 35 μL of TIS was added at room temperature. After 1 hour, the reaction was diluted in 25mL $H_2O$ and TFA was quenched with a solution of saturated sodium bicarbonate. The organic layer was collected, evaporated, and purified on a silica column in a gradient of 10:1 DCM:EtOAC, and 4:1 DCM:EtOAc. The organic layer was evaporated to a bright yellow solid with a yield of 18 mg (yield = 55%). $^{1}$H-NMR (400MHz $CDCl_3$): δ 8.62 (d, 1H), 8.57 (d, 1H), 8.48 (d, 1H), 7.74 (t, 1H), 7.27 (d, 1H), 5.79 (t, 1H), 5.39 (s, 1H), 4.78 (m, 1H) 4.42 (q, 1H), 4.04 (t, 4H), 4.00 (m, 1H), 3.30 (t, 4H), 2.93 (s, 1H), 2.40- 0.69 (m, 43H). $^{13}$C-NMR (400MHz $CDCl_3$): δ 169.0, 164.1, 163.6, 156.1, 139.4, 133.1, 130.6, 131.7, 130.1, 126.1, 125.9, 122.8, 122.8, 116.5, 115.0, 75.3, 66.9, 61.1, 56.6, 56.1, 54.5, 53.4, 49.9, 42.2, 39.7, 39.5, 37.9, 36.8, 36.5, 36.1, 35.7, 31.8, 31.8, 28.2, 28.0, 27.5, 24.2, 23.8, 22.8, 22.5, 21.0, 19.2, 18.7, 11.8. $C_{46}H_{62}N_2O_6$ , HRMS-ESI: $[M+Na]^+$ calcd. m/z = 761.45 Da, obs. m/z = 761.4478 Da

NMR spectra of all intermediates and final compounds

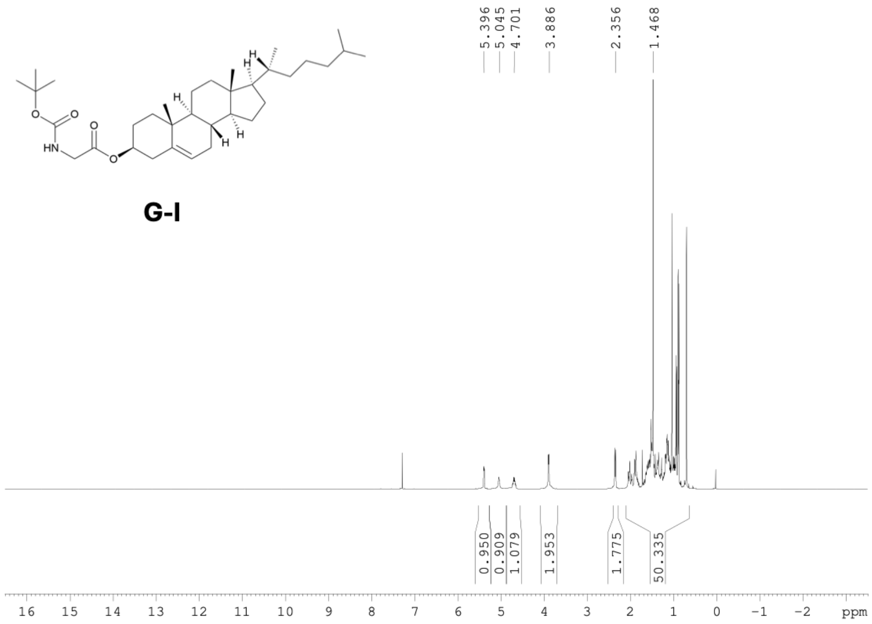


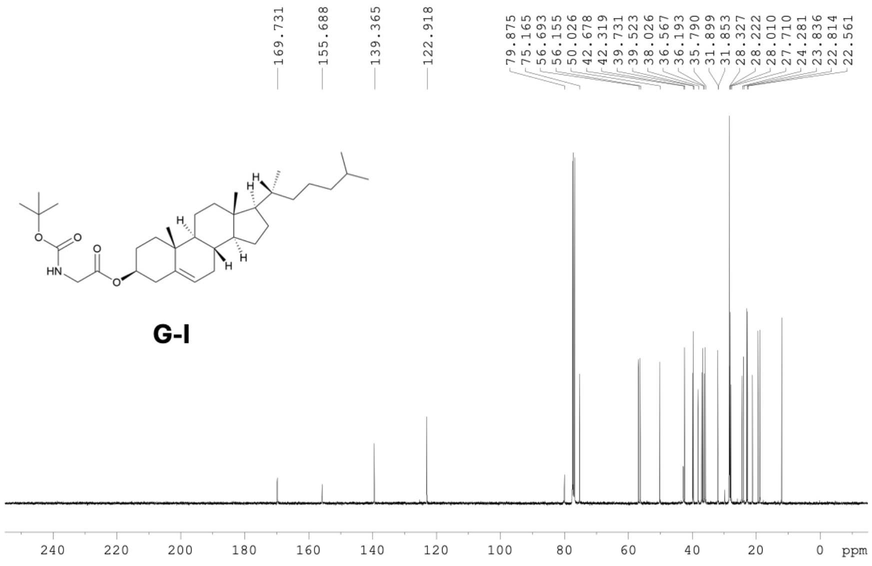

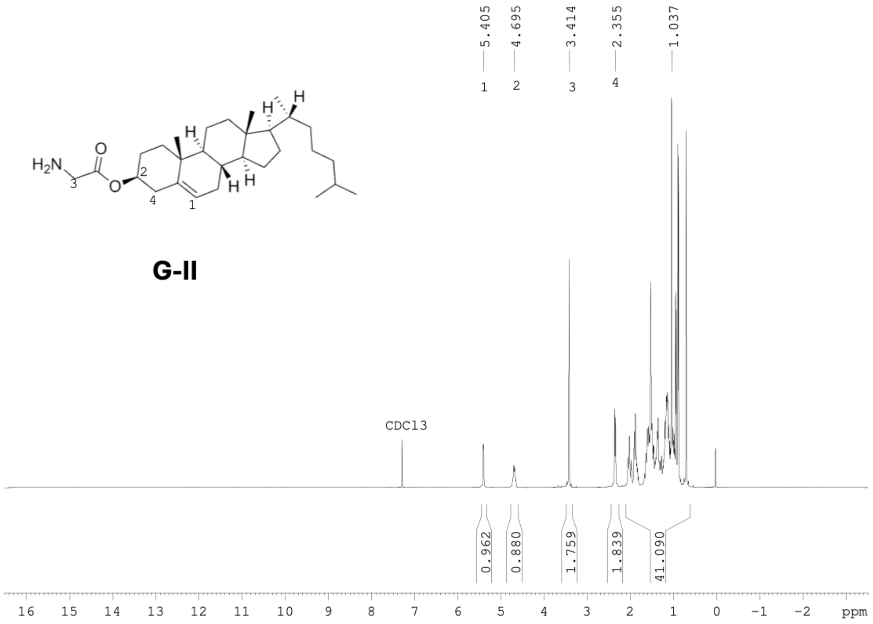
5.405
4.695
3.414
2.355
1.037
1 2 3 4
$H_2N$
O
H
G-II
CDCl3
0.962
0.880
1.759
1.839
41.090
16 15 14 13 12 11 10 9 8 7 6 5 4 3 2 1 0 −1 −2 ppm

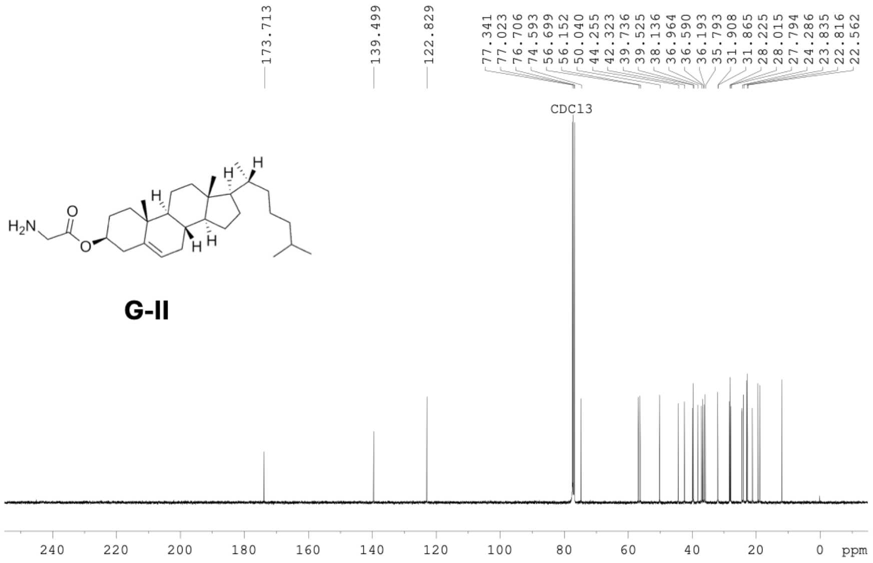
173.713
139.499
122.829
77.341
77.023
76.706
74.593
56.699
56.152
50.040
44.255
42.323
39.736
39.525
38.136
36.964
36.590
36.193
35.793
31.908
31.865
28.225
28.015
27.794
24.286
23.835
22.816
22.562
CDCl3
$H_2N$
O
H
G-II
240 220 200 180 160 140 120 100 80 60 40 20 0 ppm

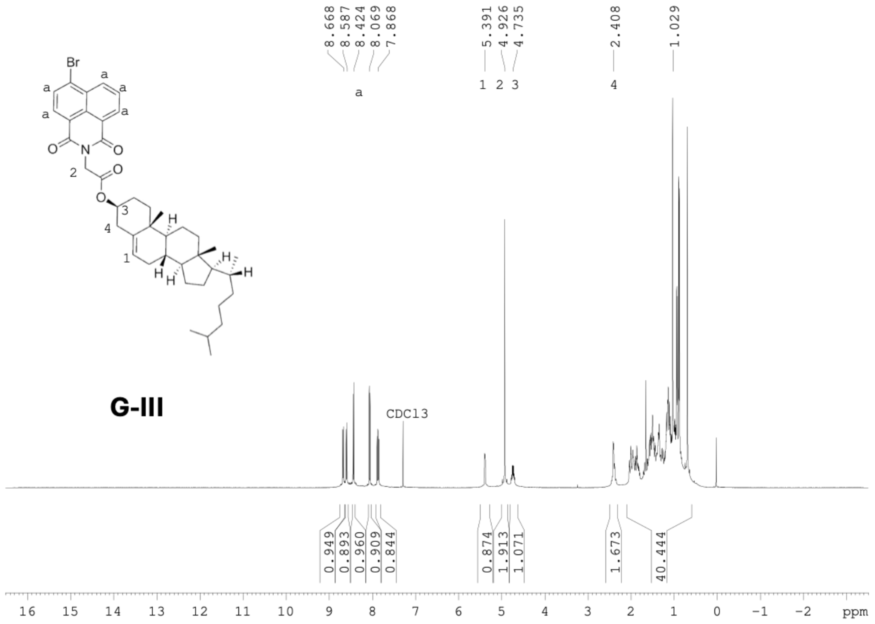
8.668
8.587
8.424
8.069
7.868
5.391
4.926
4.735
2.408
1.029
a
1 2 3
4
Br
N
O
G-III
CDCl3
0.949
0.893
0.960
0.909
0.844
0.874
1.913
1.071
1.673
40.444
16 15 14 13 12 11 10 9 8 7 6 5 4 3 2 1 0 -1 -2 ppm

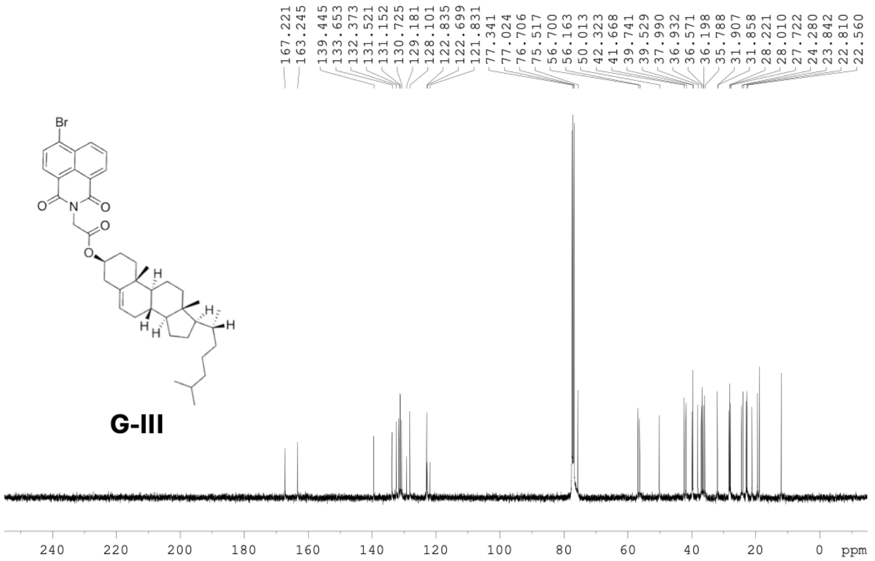
167.221
163.245
139.445
133.653
132.373
131.521
131.152
130.725
129.181
128.101
122.835
122.699
121.831
77.341
77.024
76.706
75.517
56.700
56.163
50.013
42.323
41.668
39.741
39.529
37.990
36.932
36.571
36.198
35.788
31.907
31.858
28.221
28.010
27.722
24.280
23.842
22.810
22.560
Br
G-III
240 220 200 180 160 140 120 100 80 60 40 20 0 ppm

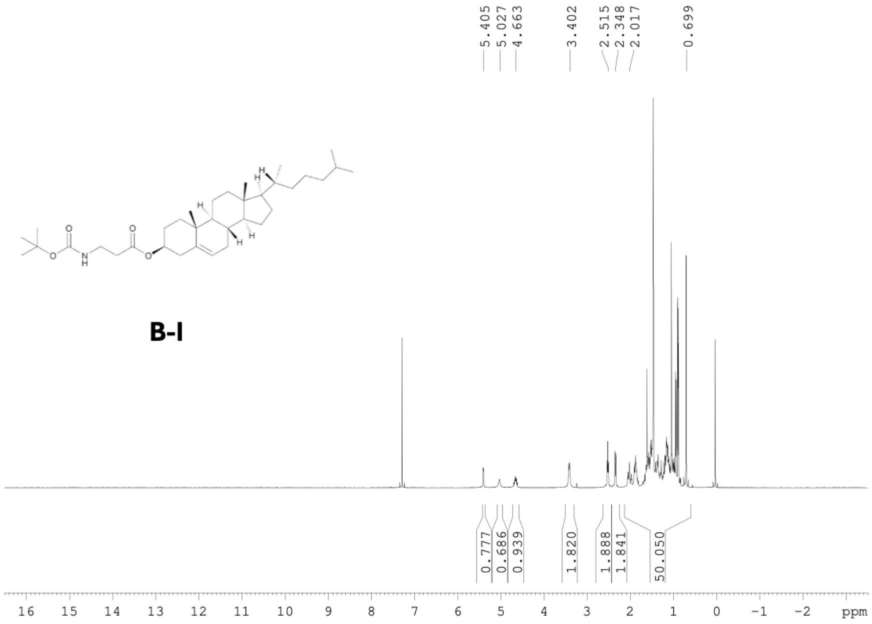
5.405
5.027
4.663
3.402
2.515
2.348
2.017
0.699
B-I
0.777
0.686
0.939
1.820
1.888
1.841
50.050
16 15 14 13 12 11 10 9 8 7 6 5 4 3 2 1 0 −1 −2 ppm

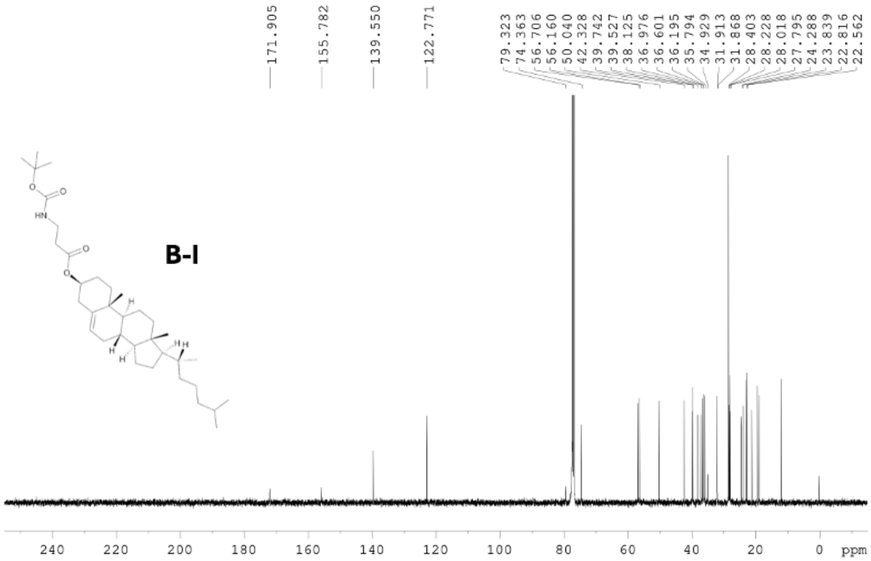
171.905
155.782
139.550
122.771
79.323
74.363
56.706
56.160
50.040
42.328
39.742
39.527
38.125
36.976
36.601
36.195
35.794
34.929
31.913
31.868
28.403
28.228
28.018
27.795
24.288
23.839
22.816
22.562
B-I
240 220 200 180 160 140 120 100 80 60 40 20 0 ppm

B-III

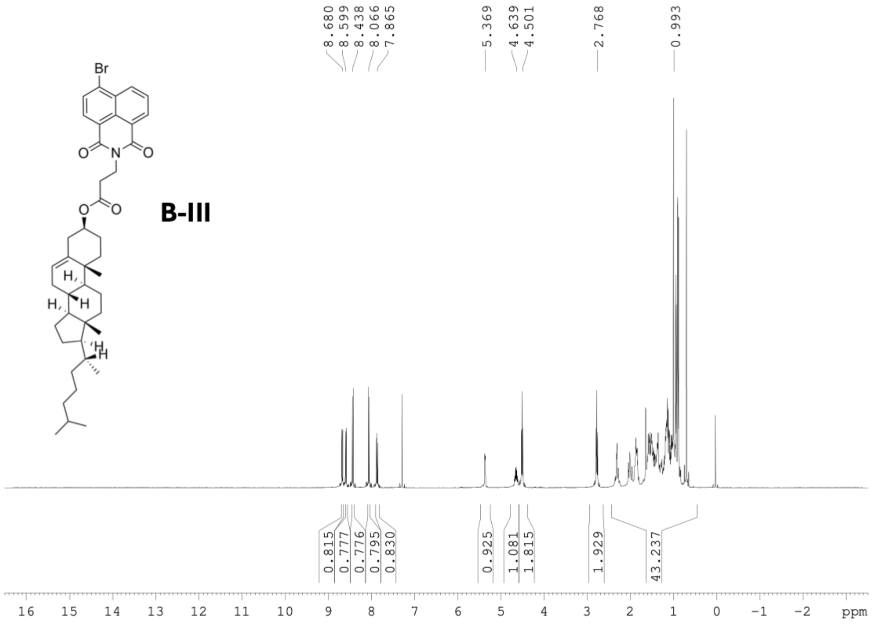

Br
O
N
O
O
O
H
H
H
H
8.680
8.599
8.438
8.066
7.865
5.369
4.639
4.501
2.768
0.993
0.815
0.777
0.776
0.795
0.830
0.925
1.081
1.815
1.929
43.237
16 15 14 13 12 11 10 9 8 7 6 5 4 3 2 1 0 −1 −2 ppm


B-III

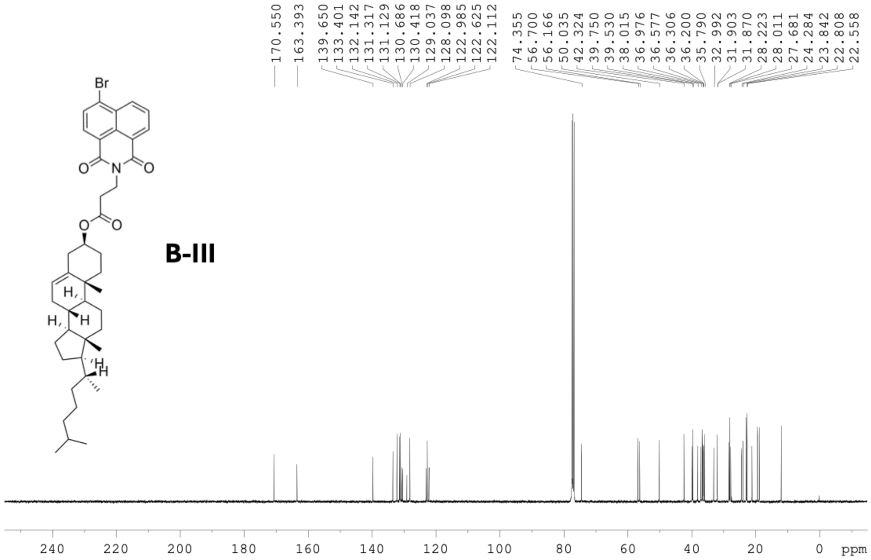

Br
O
N
O
O
O
H
H
H
H
170.550
163.393
139.650
133.401
132.142
131.317
131.129
130.686
130.418
129.037
128.098
122.985
122.625
122.112
74.355
56.700
56.166
50.035
42.324
39.750
39.530
38.015
36.976
36.577
36.306
36.200
35.790
32.992
31.903
31.870
28.223
28.011
27.681
24.284
23.842
22.808
22.558
240 220 200 180 160 140 120 100 80 60 40 20 0 ppm

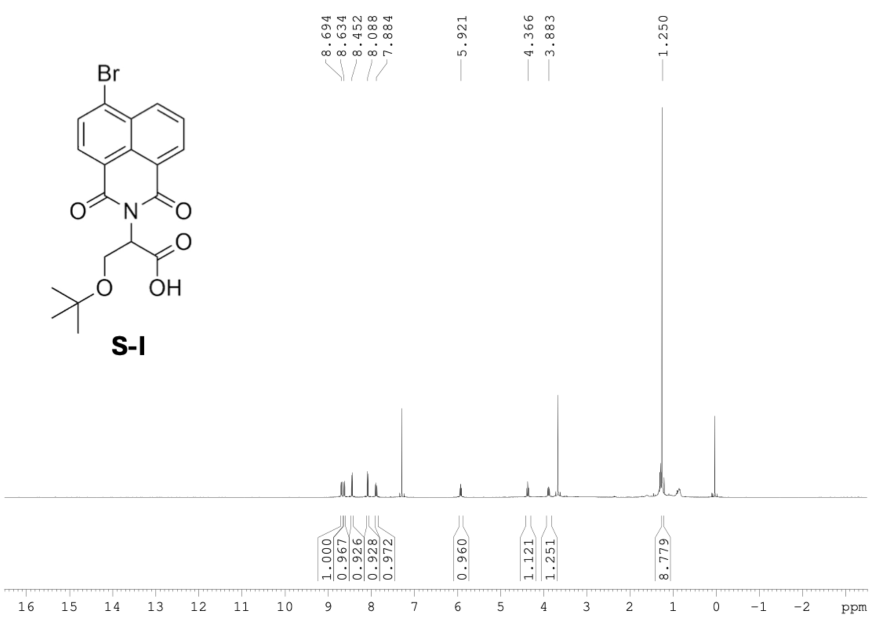
8.694
8.634
8.452
8.088
7.884
5.921
4.366
3.883
1.250
Br
O
N
O
O
O
OH
S-I
1.000
0.967
0.926
0.928
0.972
0.960
1.121
1.251
8.779
16 15 14 13 12 11 10 9 8 7 6 5 4 3 2 1 0 -1 -2 ppm

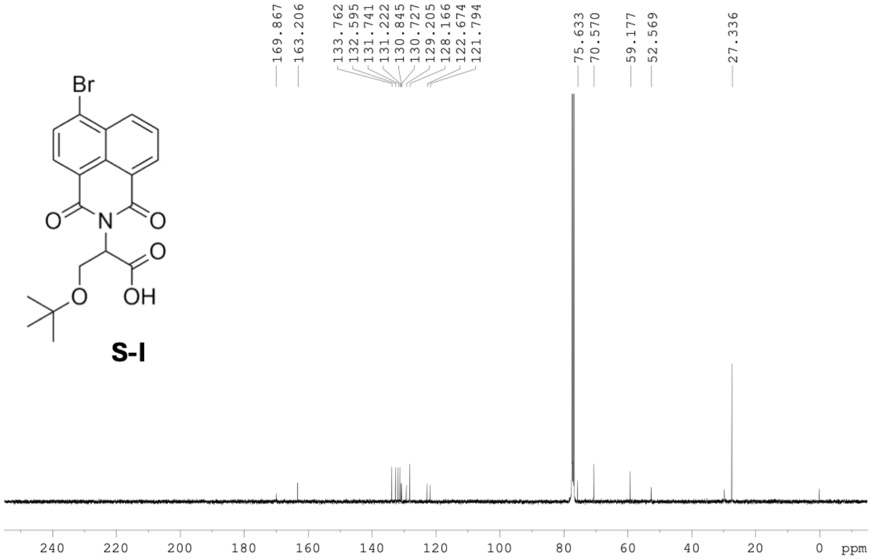
169.867
163.206
133.762
132.595
131.741
131.222
130.845
130.727
129.205
128.166
122.674
121.794
75.633
70.570
59.177
52.569
27.336
Br
O
N
O
O
O
OH
S-I
240 220 200 180 160 140 120 100 80 60 40 20 0 ppm

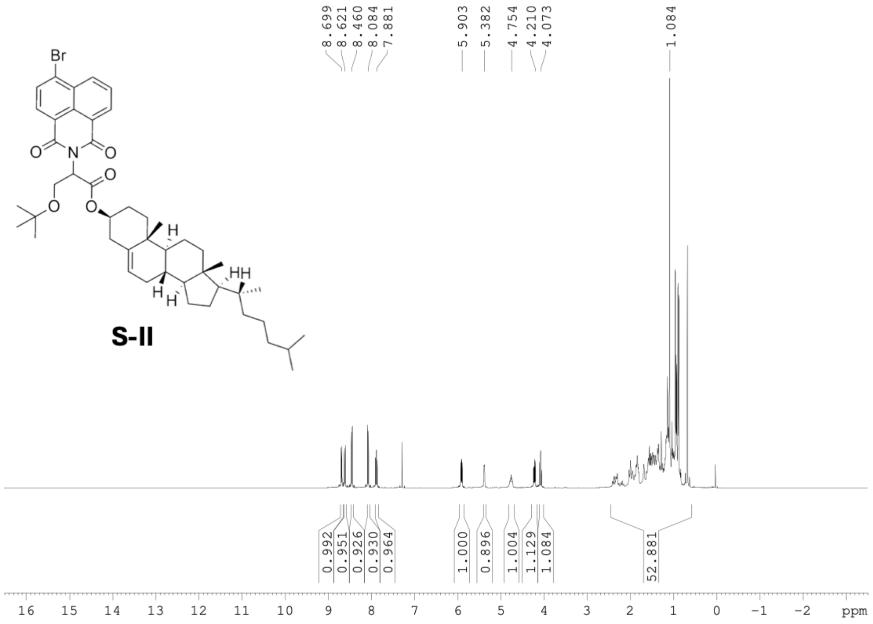
8.699
8.621
8.460
8.084
7.881
5.903
5.382
4.754
4.210
4.073
1.084
Br
O
N
O
O
O
O
H
H
H
HH
S-II
0.992
0.951
0.926
0.930
0.964
1.000
0.896
1.004
1.129
1.084
52.881
16 15 14 13 12 11 10 9 8 7 6 5 4 3 2 1 0 −1 −2 ppm

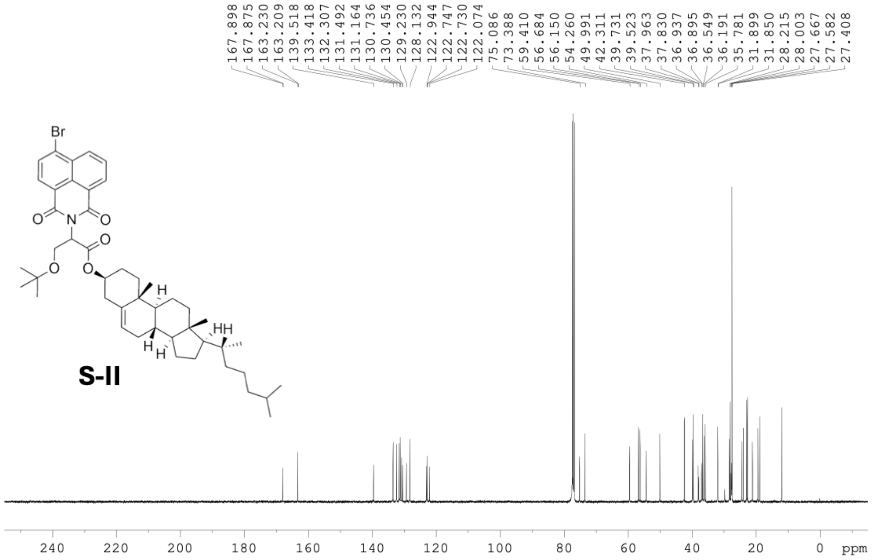
167.898
167.875
163.230
163.209
139.518
133.418
132.307
131.492
131.164
130.736
130.454
129.230
128.132
122.944
122.747
122.730
122.074
75.086
73.388
59.410
56.684
56.150
54.260
49.991
42.311
39.731
39.523
37.963
37.830
36.937
36.895
36.549
36.191
35.781
31.899
31.850
28.215
28.003
27.667
27.582
27.408
Br
O
N
O
O
O
O
H
H
HH
S-II
240 220 200 180 160 140 120 100 80 60 40 20 0 ppm

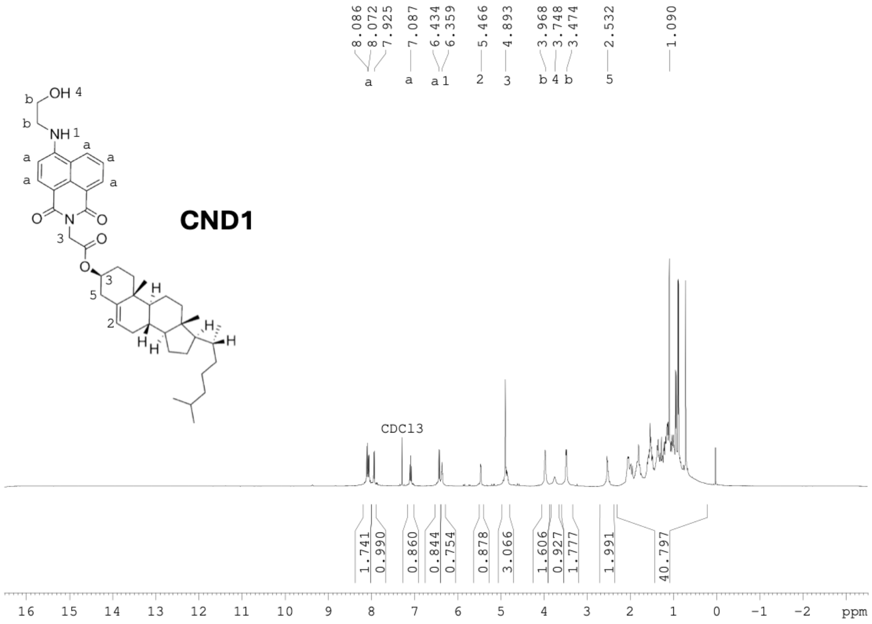
CND1
CDCl3
ppm

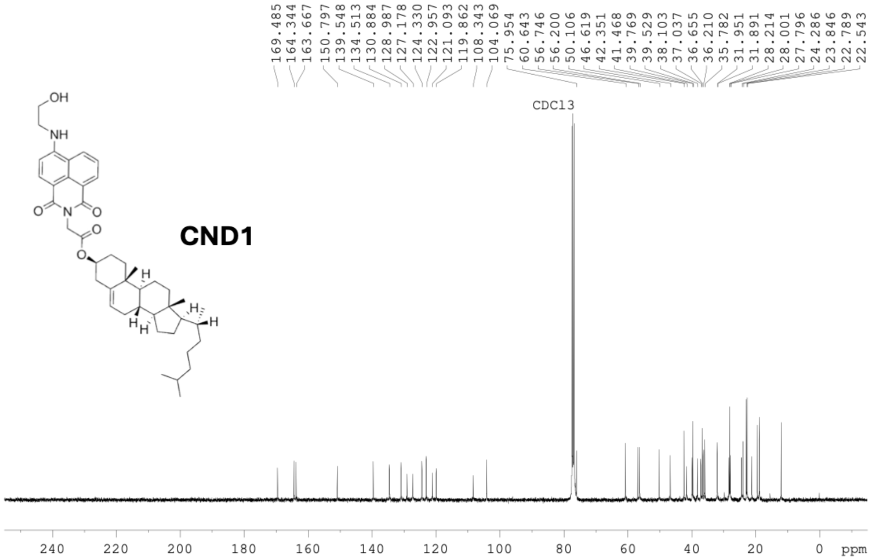
CND1
CDCl3
ppm

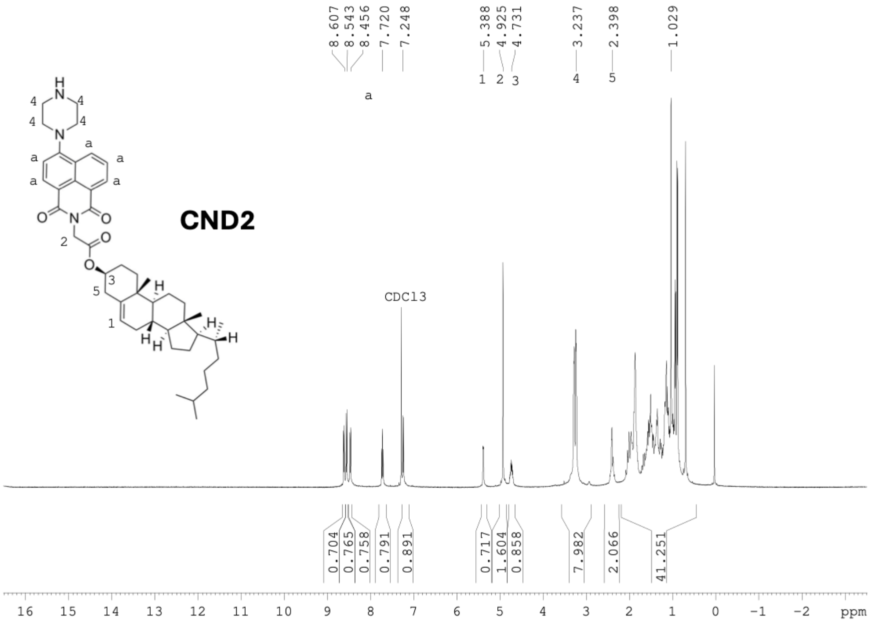
CND2
CDCl3
8.607
8.543
8.456
7.720
7.248
5.388
4.925
4.731
3.237
2.398
1.029
0.704
0.765
0.758
0.791
0.891
0.717
1.604
0.858
7.982
2.066
41.251
ppm

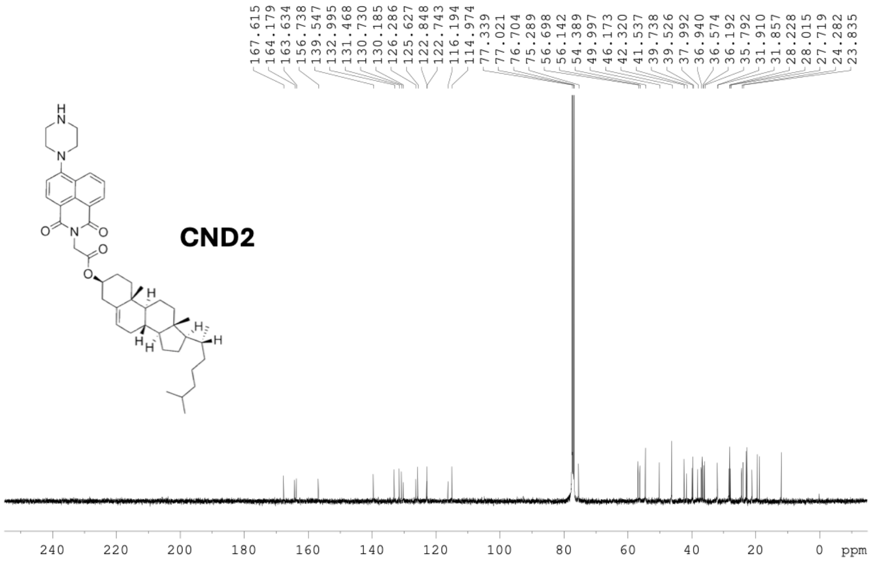
CND2
167.615
164.179
163.634
156.738
139.547
132.995
131.468
130.730
130.185
126.286
125.627
122.848
122.743
116.194
114.974
77.339
77.021
76.704
75.289
56.698
56.142
54.389
49.997
46.173
42.320
41.537
39.738
39.526
37.992
36.940
36.574
36.192
35.792
31.910
31.857
28.228
28.015
27.719
24.282
23.835
ppm

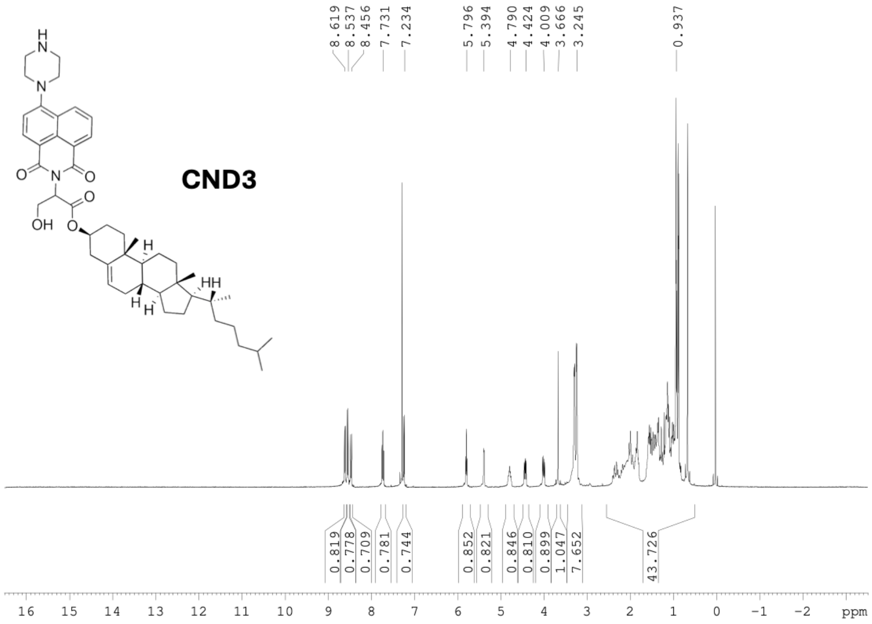
CND3
8.619
8.537
8.456
7.731
7.234
5.796
5.394
4.790
4.424
4.009
3.666
3.245
0.937
0.819
0.778
0.709
0.781
0.744
0.852
0.821
0.846
0.810
0.899
1.047
7.652
43.726
16 15 14 13 12 11 10 9 8 7 6 5 4 3 2 1 0 -1 -2 ppm

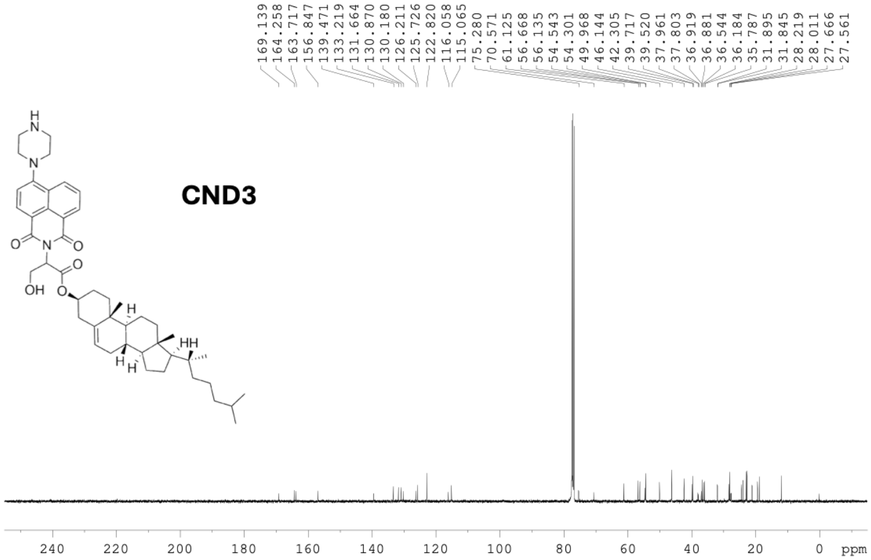
CND3
169.139
164.258
163.717
156.847
139.471
133.219
131.664
130.870
130.180
126.211
125.726
122.820
116.058
115.065
75.280
70.571
61.125
56.668
56.135
54.543
54.301
49.968
46.144
42.305
39.717
39.520
37.961
37.803
36.919
36.881
36.544
36.184
35.787
31.895
31.845
28.219
28.011
27.666
27.561
240 220 200 180 160 140 120 100 80 60 40 20 0 ppm

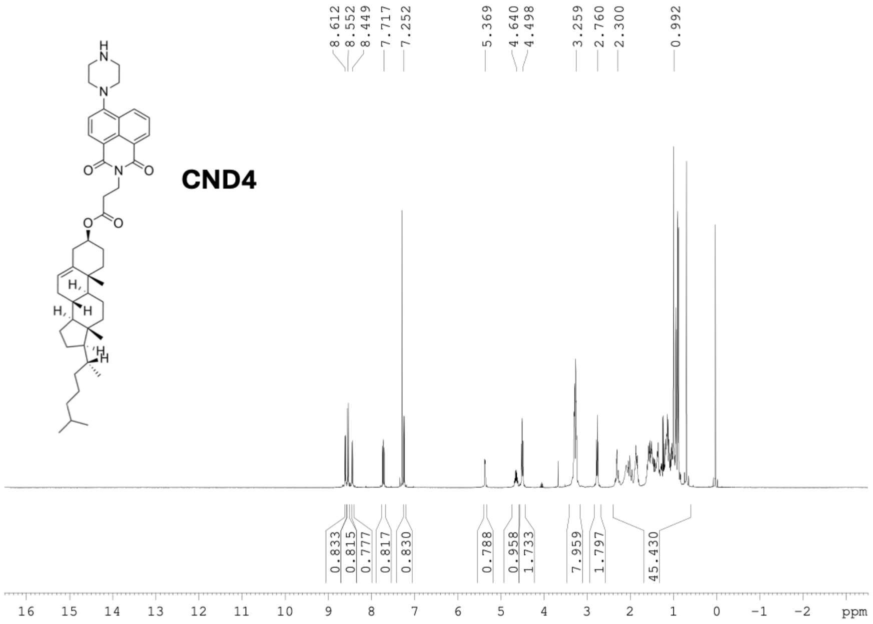
CND4
8.612
8.552
8.449
7.717
7.252
5.369
4.640
4.498
3.259
2.760
2.300
0.992
0.833
0.815
0.777
0.817
0.830
0.788
0.958
1.733
7.959
1.797
45.430
ppm

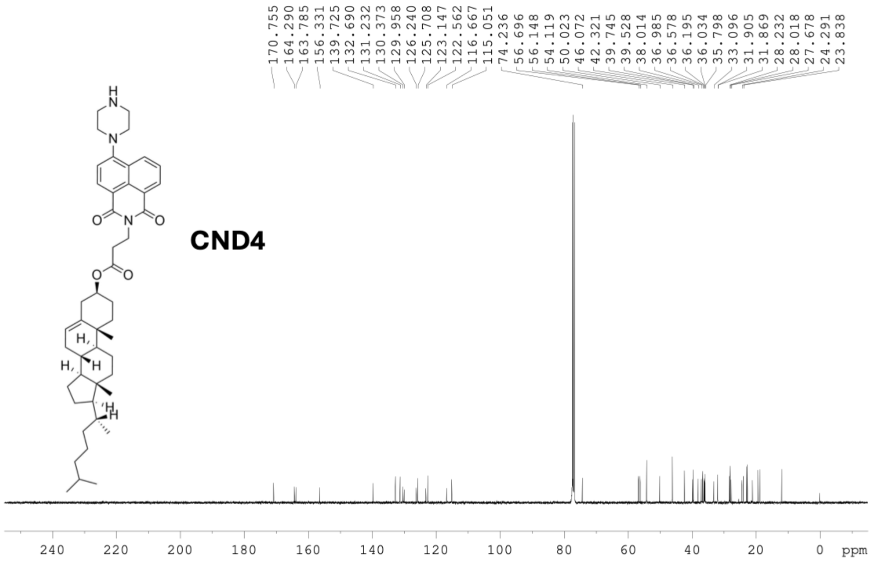
CND4
170.755
164.290
163.785
156.331
139.725
132.690
131.232
130.373
129.958
126.240
125.708
123.147
122.562
116.667
115.051
74.236
56.696
56.148
54.119
50.023
46.072
42.321
39.745
39.528
38.014
36.985
36.578
36.195
36.034
35.798
33.096
31.905
31.869
28.232
28.018
27.678
24.291
23.838
ppm

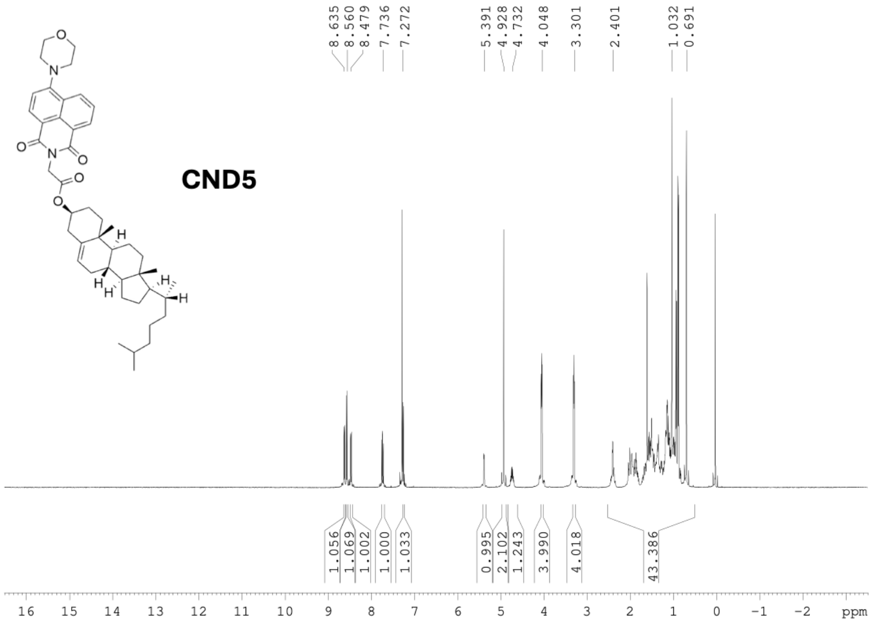

CND5
8.635
8.560
8.479
7.736
7.272
5.391
4.928
4.732
4.048
3.301
2.401
1.032
0.691
1.056
1.069
1.002
1.000
1.033
0.995
2.102
1.243
3.990
4.018
43.386
ppm


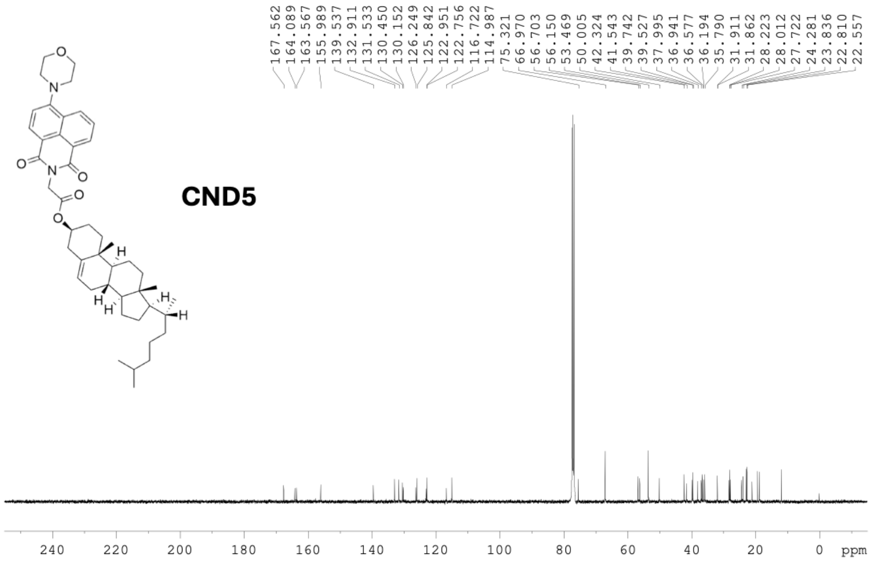

CND5
167.562
164.089
163.567
155.989
139.537
132.911
131.533
130.450
130.152
126.249
125.842
122.951
122.756
116.722
114.987
75.321
66.970
56.703
56.150
53.469
50.005
42.324
41.543
39.742
39.527
37.995
36.941
36.577
36.194
35.790
31.911
31.862
28.223
28.012
27.722
24.281
23.836
22.810
22.557
ppm

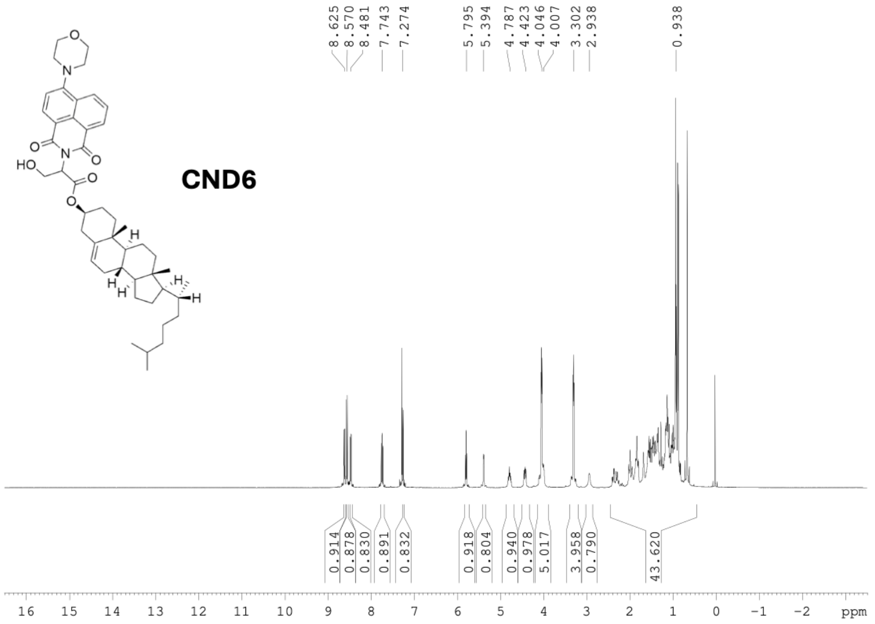

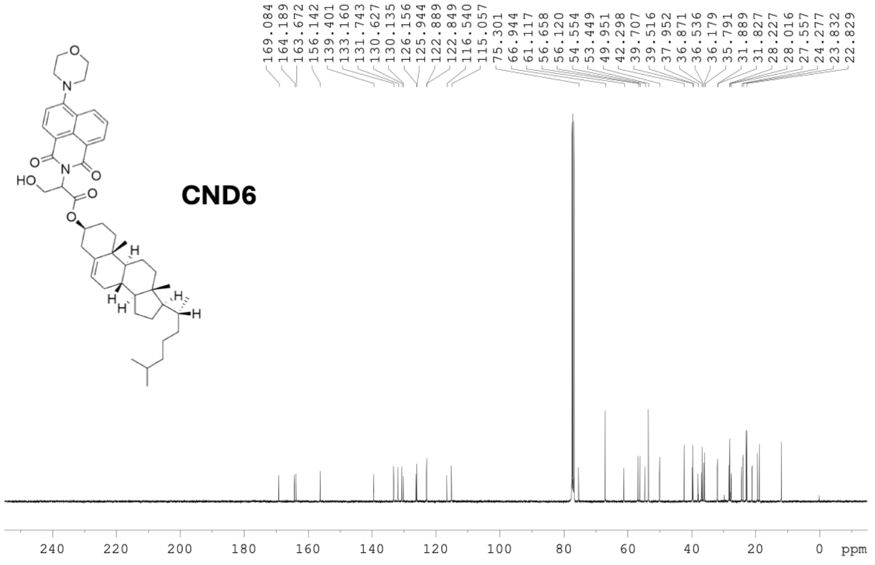

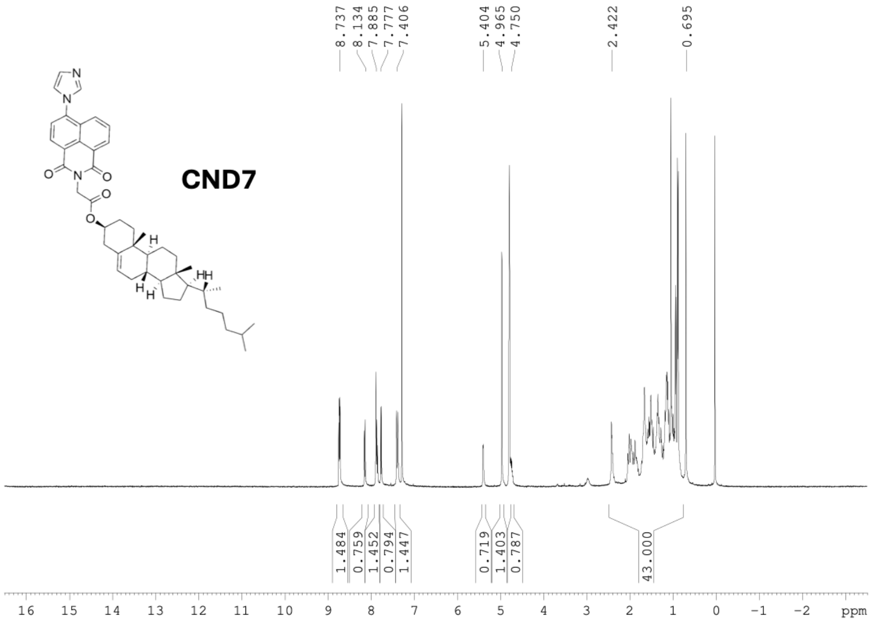
CND7
8.737
8.134
7.885
7.777
7.406
5.404
4.965
4.750
2.422
0.695
1.484
0.759
1.452
0.794
1.447
0.719
1.403
0.787
43.000
16 15 14 13 12 11 10 9 8 7 6 5 4 3 2 1 0 -1 -2 ppm

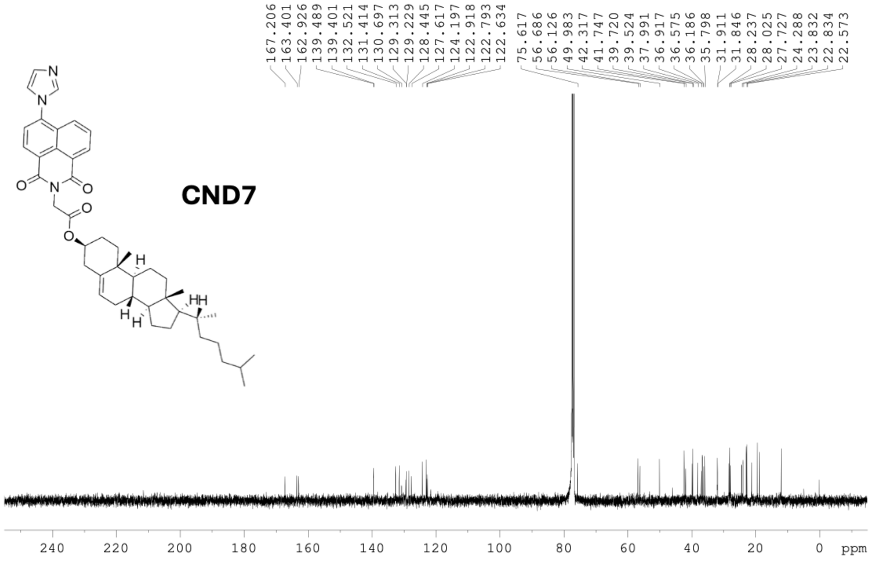
CND7
167.206
163.401
162.926
139.489
139.401
132.521
131.414
130.697
129.313
129.229
128.445
127.617
124.197
122.918
122.793
122.634
75.617
56.686
56.126
49.983
42.317
41.747
39.720
39.524
37.991
36.917
36.575
36.186
35.798
31.911
31.846
28.237
28.025
27.727
24.288
23.832
22.834
22.573
240 220 200 180 160 140 120 100 80 60 40 20 0 ppm

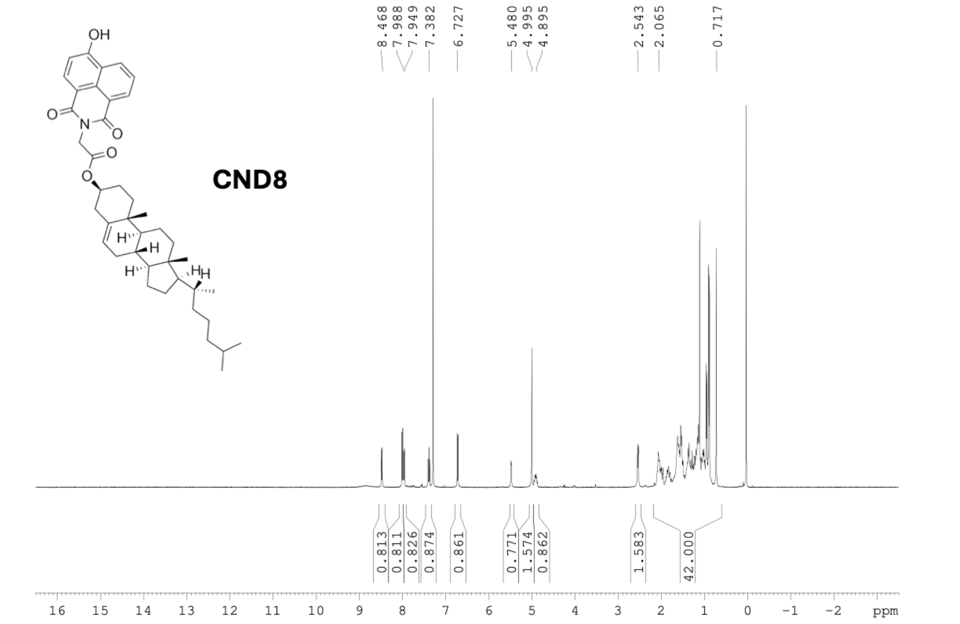
OH
O
N
O
O
O
H
H
H
H
CND8
8.468
7.988
7.949
7.382
6.727
5.480
4.995
4.895
2.543
2.065
0.717
0.813
0.811
0.826
0.874
0.861
0.771
1.574
0.862
1.583
42.000
16 15 14 13 12 11 10 9 8 7 6 5 4 3 2 1 0 -1 -2 ppm

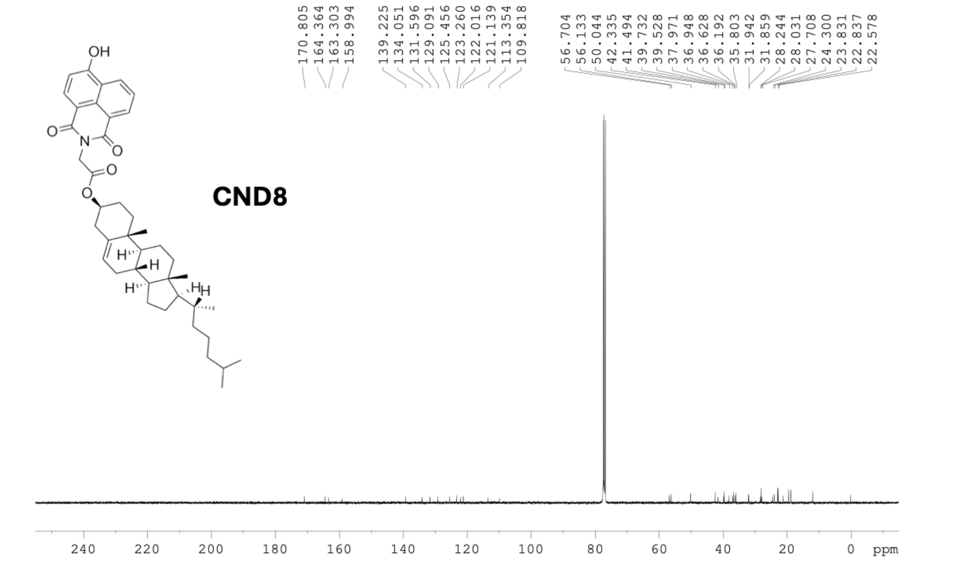
OH
O
N
O
O
O
H
H
H
H
CND8
170.805
164.364
163.303
158.994
139.225
134.051
131.596
129.091
125.456
123.260
122.016
121.139
113.354
109.818
56.704
56.133
50.044
42.335
41.494
39.732
39.528
37.971
36.948
36.628
36.192
35.803
31.942
31.859
28.244
28.031
27.708
24.300
23.831
22.837
22.578
240 220 200 180 160 140 120 100 80 60 40 20 0 ppm

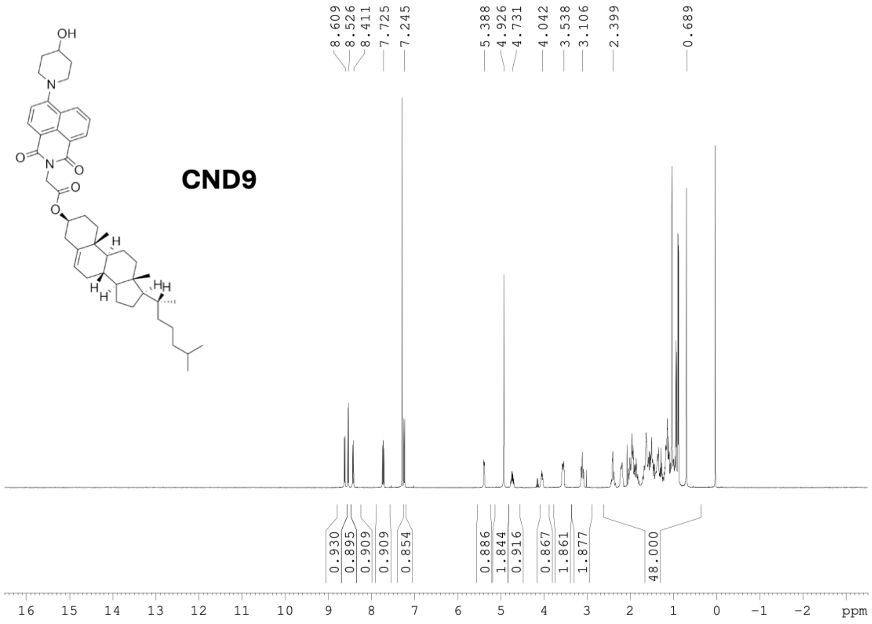

OH
N
O
N
O
O
O
H
H
H
H
CND9
8.609
8.526
8.411
7.725
7.245
5.388
4.926
4.731
4.042
3.538
3.106
2.399
0.689
0.930
0.895
0.909
0.909
0.854
0.886
1.844
0.916
0.867
1.861
1.877
48.000
16 15 14 13 12 11 10 9 8 7 6 5 4 3 2 1 0 −1 −2 ppm


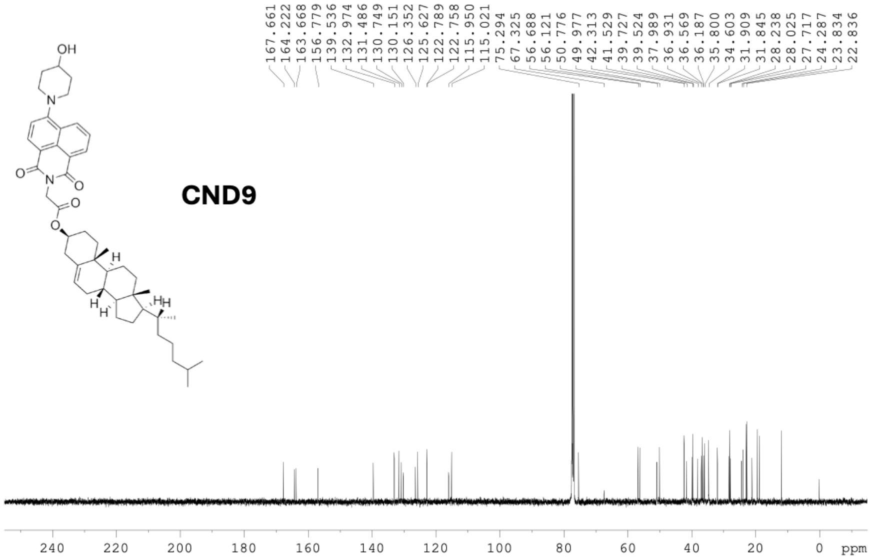

OH
N
O
N
O
O
O
H
H
H
H
CND9
167.661
164.222
163.668
156.779
139.536
132.974
131.486
130.749
130.151
126.352
125.627
122.789
122.758
115.950
115.021
75.294
67.325
56.688
56.121
50.776
49.977
42.313
41.529
39.727
39.524
37.989
36.931
36.569
36.187
35.800
34.603
31.909
31.845
28.238
28.025
27.717
24.287
23.834
22.836
240 220 200 180 160 140 120 100 80 60 40 20 0 ppm

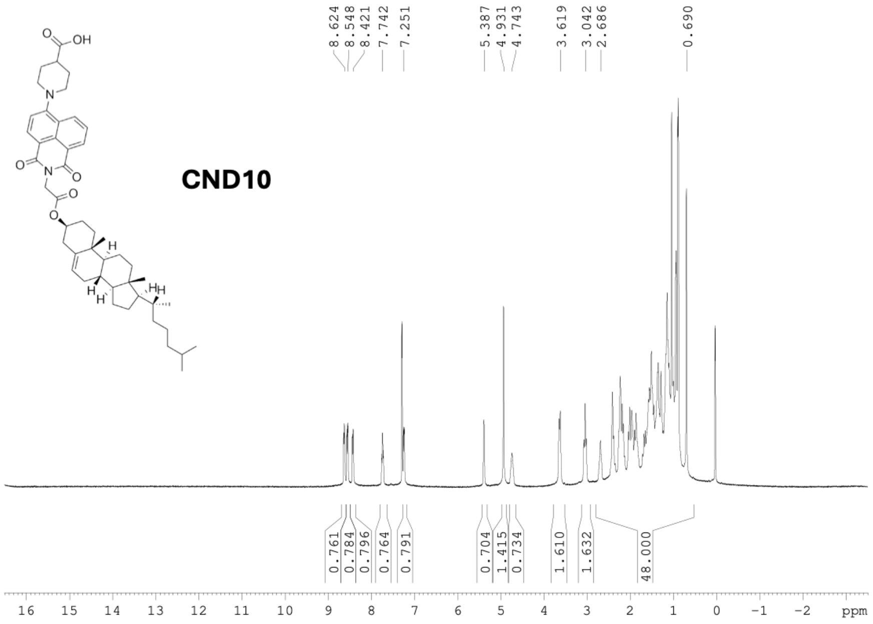
CND10
8.624
8.548
8.421
7.742
7.251
5.387
4.931
4.743
3.619
3.042
2.686
0.690
0.761
0.784
0.796
0.764
0.791
0.704
1.415
0.734
1.610
1.632
48.000
16 15 14 13 12 11 10 9 8 7 6 5 4 3 2 1 0 −1 −2 ppm

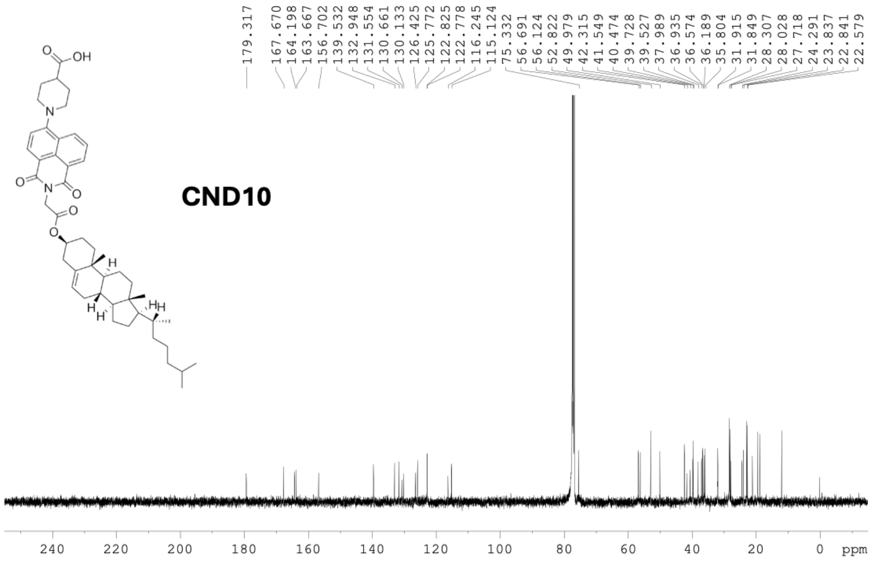
CND10
179.317
167.670
164.198
163.667
156.702
139.532
132.948
131.554
130.661
130.133
126.425
125.772
122.825
122.778
116.245
115.124
75.332
56.691
56.124
52.822
49.979
42.315
41.549
40.474
39.728
39.527
37.989
36.935
36.574
36.189
35.804
31.915
31.849
28.307
28.028
27.718
24.291
23.837
22.841
22.579
240 220 200 180 160 140 120 100 80 60 40 20 0 ppm

HR-MS spectra of all compounds

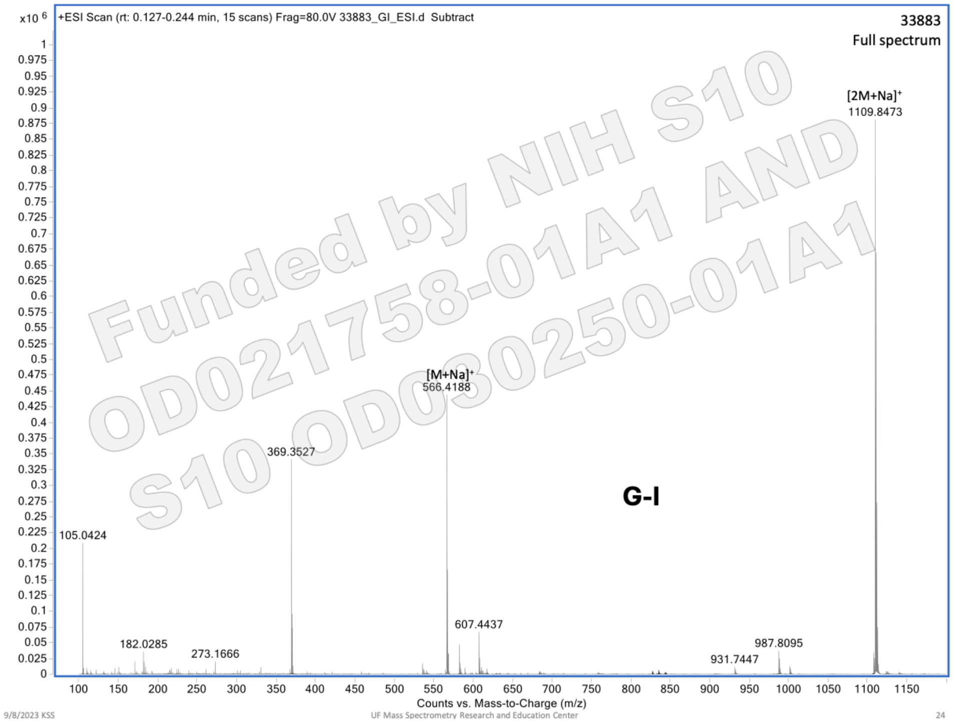


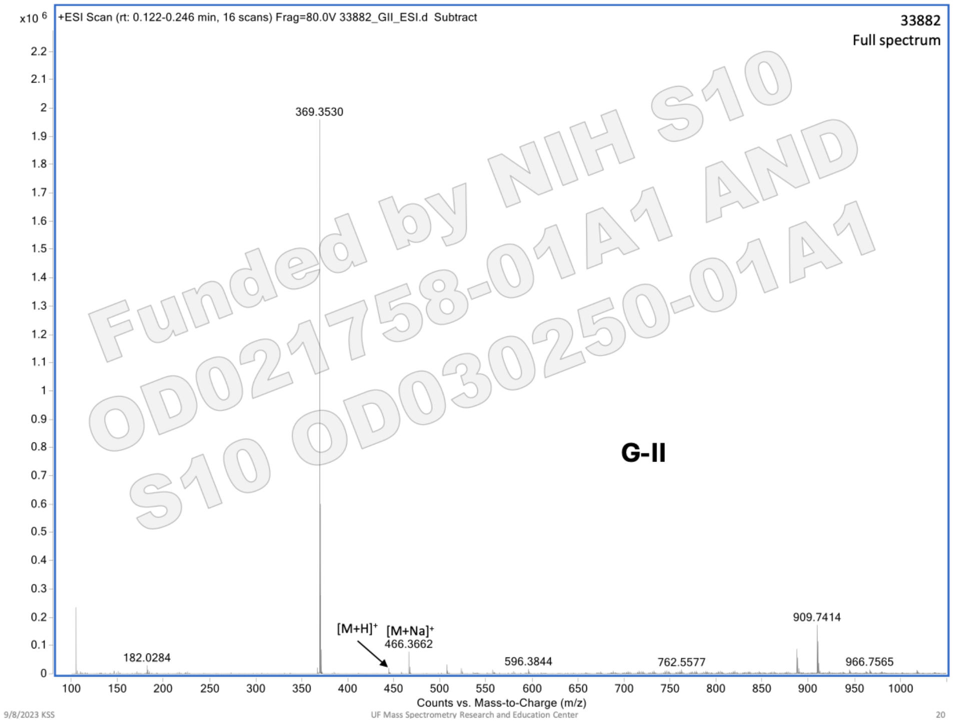

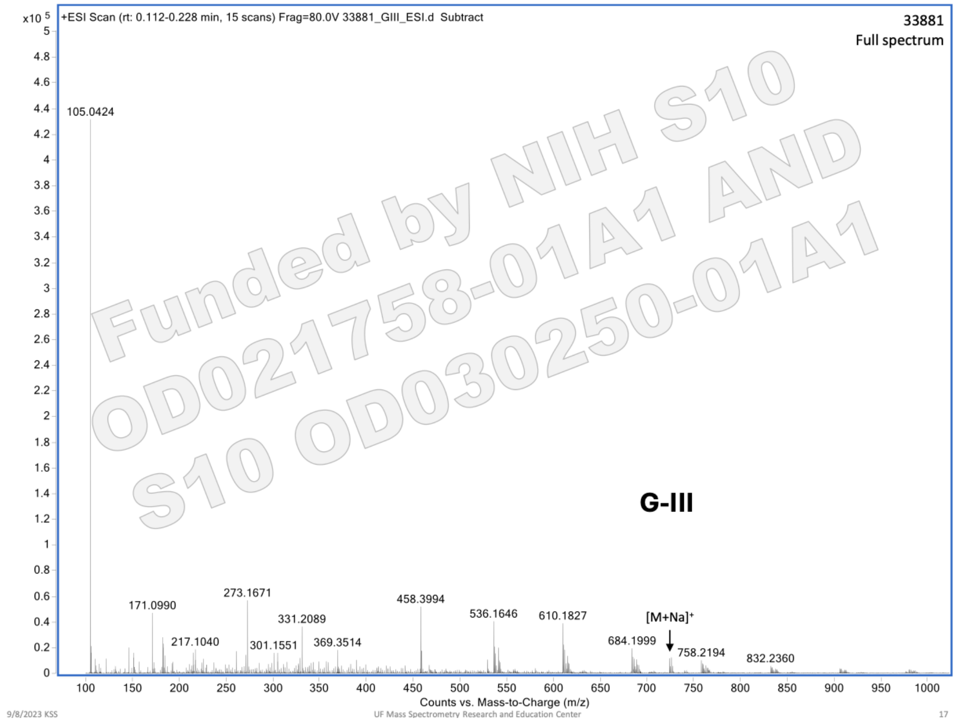
+ESI Scan (rt: 0.112-0.228 min, 15 scans) Frag=80.0V 33881_GIII_ESI.d Subtract
33881
Full spectrum
x10 5
105.0424
171.0990
217.1040
273.1671
301.1551
331.2089
369.3514
458.3994
536.1646
610.1827
684.1999
[M+Na]+
758.2194
832.2360
G-III
Counts vs. Mass-to-Charge (m/z)
Funded by NIH S10 OD021758-01A1 AND S10 OD030250-01A1
9/8/2023 KSS
UF Mass Spectrometry Research and Education Center
17

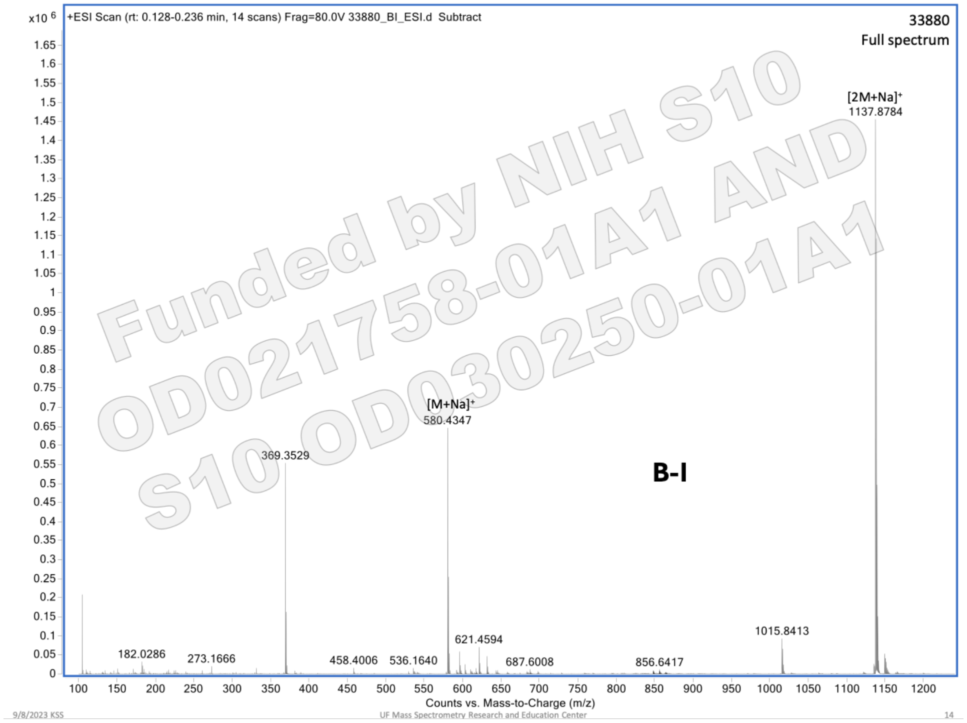
+ESI Scan (rt: 0.128-0.236 min, 14 scans) Frag=80.0V 33880_BI_ESI.d Subtract
33880
Full spectrum
x10 6
[2M+Na]+
1137.8784
[M+Na]+
580.4347
369.3529
182.0286
273.1666
458.4006
536.1640
621.4594
687.6008
856.6417
1015.8413
B-I
Counts vs. Mass-to-Charge (m/z)
Funded by NIH S10 OD021758-01A1 AND S10 OD030250-01A1
9/8/2023 KSS
UF Mass Spectrometry Research and Education Center
14

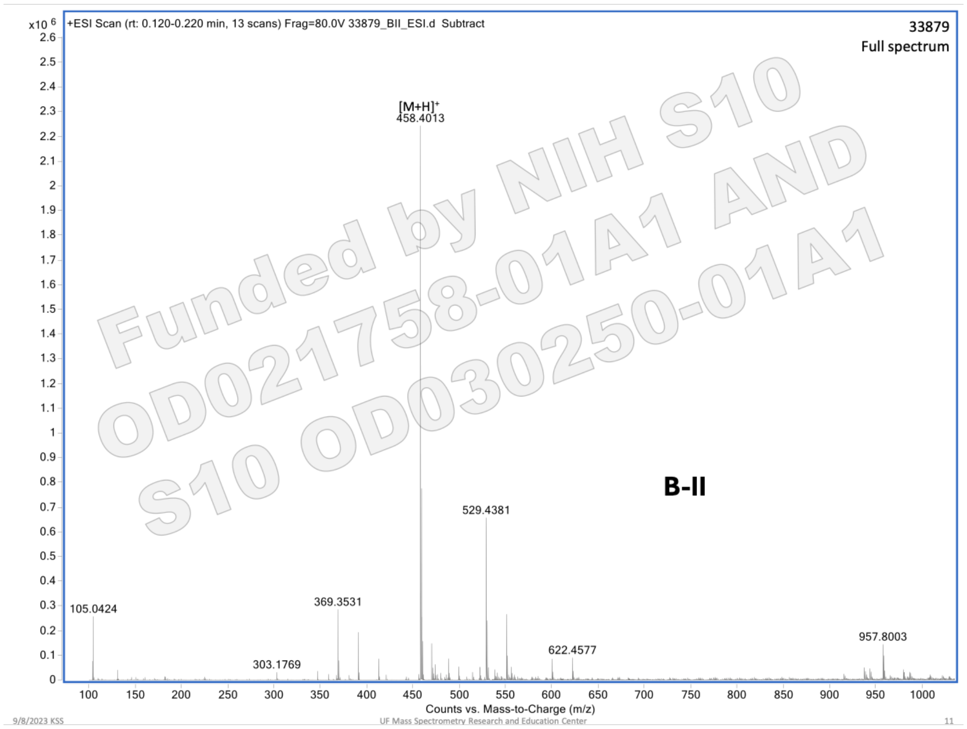
+ESI Scan (rt: 0.120-0.220 min, 13 scans) Frag=80.0V 33879_BII_ESI.d Subtract
33879
Full spectrum
[M+H]+
458.4013
529.4381
369.3531
105.0424
303.1769
622.4577
957.8003
B-II
Funded by NIH S10 OD021758-01A1 AND S10 OD030250-01A1
Counts vs. Mass-to-Charge (m/z)
9/8/2023 KSS
UF Mass Spectrometry Research and Education Center
11

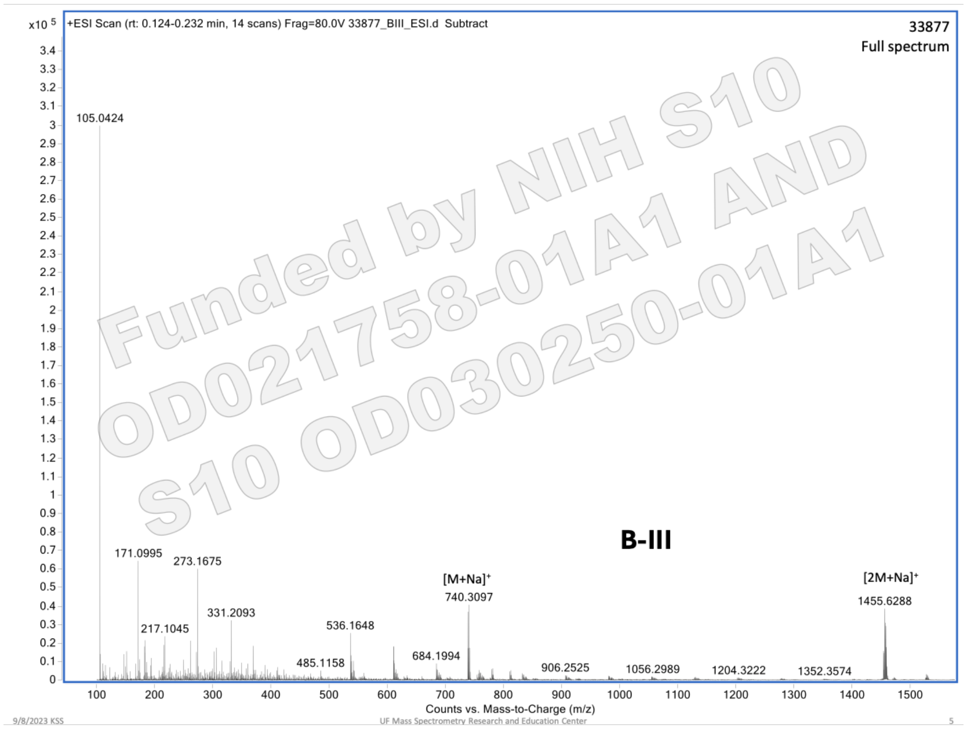
+ESI Scan (rt: 0.124-0.232 min, 14 scans) Frag=80.0V 33877_BIII_ESI.d Subtract
33877
Full spectrum
x10 5
105.0424
171.0995
273.1675
217.1045
331.2093
536.1648
485.1158
[M+Na]+
740.3097
684.1994
906.2525
1056.2989
1204.3222
1352.3574
[2M+Na]+
1455.6288
B-III
Counts vs. Mass-to-Charge (m/z)
9/8/2023 KS5
UF Mass Spectrometry Research and Education Center
5

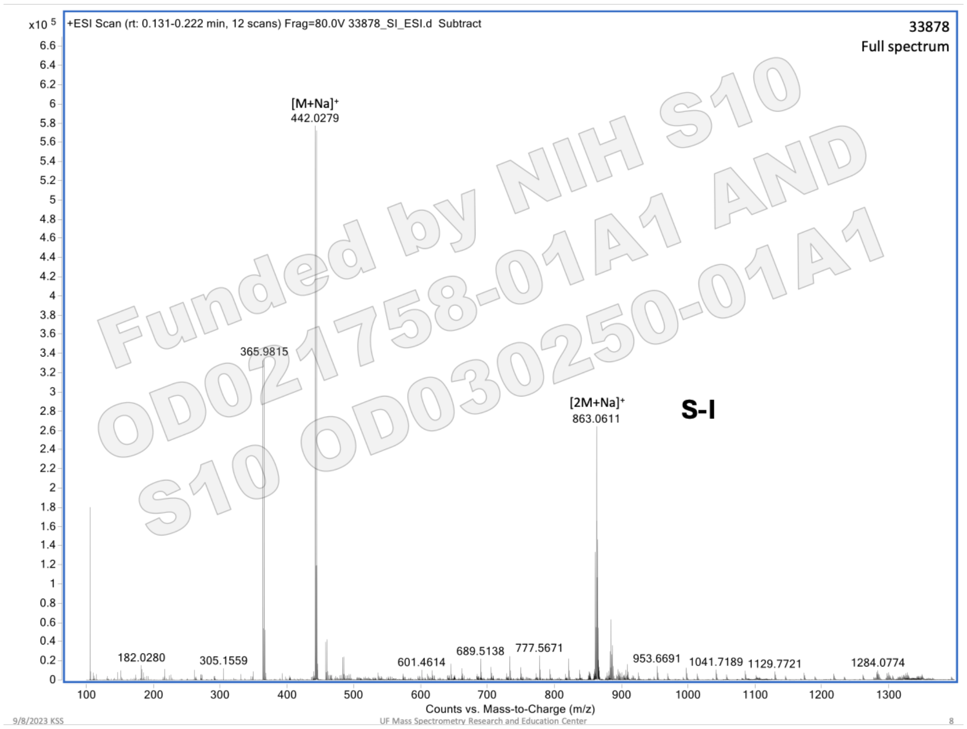
+ESI Scan (rt: 0.131-0.222 min, 12 scans) Frag=80.0V 33878_SI_ESI.d Subtract
33878
Full spectrum
x10 5
[M+Na]+
442.0279
365.9815
[2M+Na]+
863.0611
S-I
182.0280
305.1559
601.4614
689.5138
777.5671
953.6691
1041.7189
1129.7721
1284.0774
Counts vs. Mass-to-Charge (m/z)
9/8/2023 KS5
UF Mass Spectrometry Research and Education Center
8

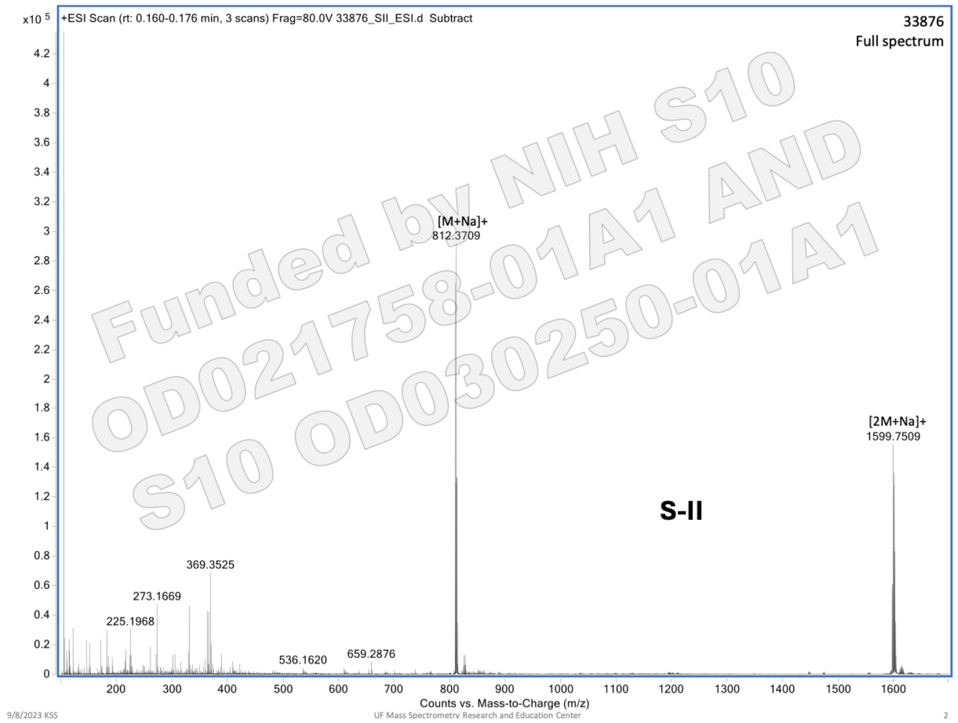

+ESI Scan (rt: 0.160-0.176 min, 3 scans) Frag=80.0V 33876_SII_ESI.d Subtract
33876
Full spectrum
[M+Na]+
812.3709
[2M+Na]+
1599.7509
S-II
369.3525
273.1669
225.1968
536.1620
659.2876
Counts vs. Mass-to-Charge (m/z)
9/8/2023 KSS
UF Mass Spectrometry Research and Education Center
2


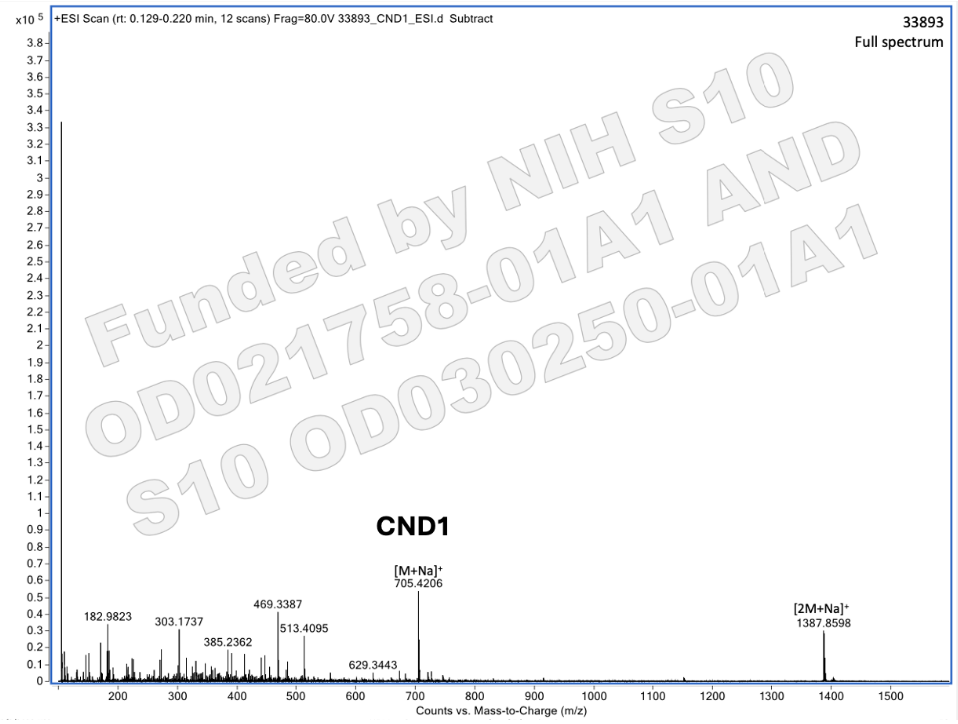

+ESI Scan (rt: 0.129-0.220 min, 12 scans) Frag=80.0V 33893_CND1_ESI.d Subtract
33893
Full spectrum
CND1
[M+Na]+
705.4206
[2M+Na]+
1387.8598
182.9823
303.1737
385.2362
469.3387
513.4095
629.3443
Counts vs. Mass-to-Charge (m/z)

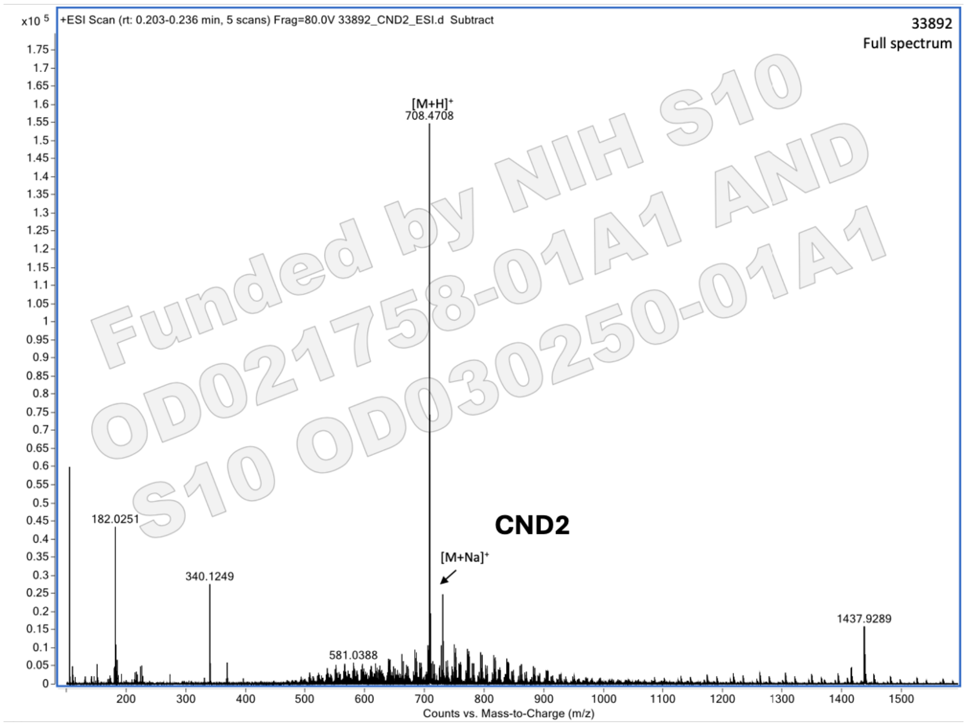
+ESI Scan (rt: 0.203-0.236 min, 5 scans) Frag=80.0V 33892_CND2_ESI.d Subtract
33892
Full spectrum
x10 5
[M+H]+
708.4708
182.0251
340.1249
581.0388
[M+Na]+
1437.9289
CND2
Counts vs. Mass-to-Charge (m/z)
Funded by NIH S10 OD021758-01A1 AND S10 OD030250-01A1

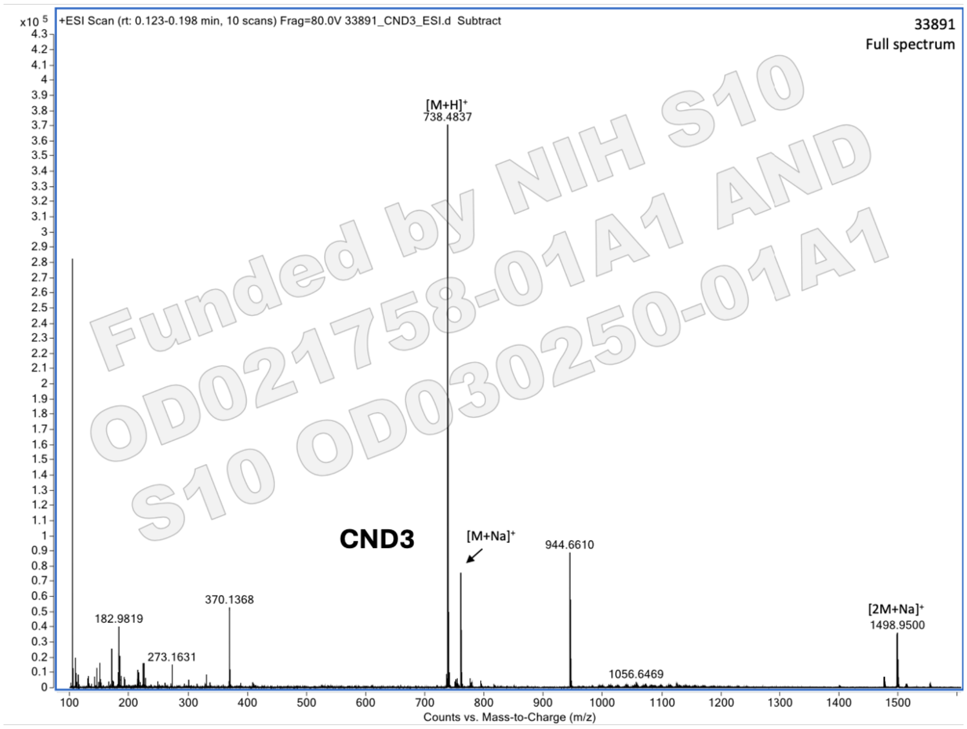
+ESI Scan (rt: 0.123-0.198 min, 10 scans) Frag=80.0V 33891_CND3_ESI.d Subtract
33891
Full spectrum
x10 5
[M+H]+
738.4837
CND3
[M+Na]+
944.6610
370.1368
182.9819
273.1631
1056.6469
[2M+Na]+
1498.9500
Counts vs. Mass-to-Charge (m/z)
Funded by NIH S10 OD021758-01A1 AND S10 OD030250-01A1

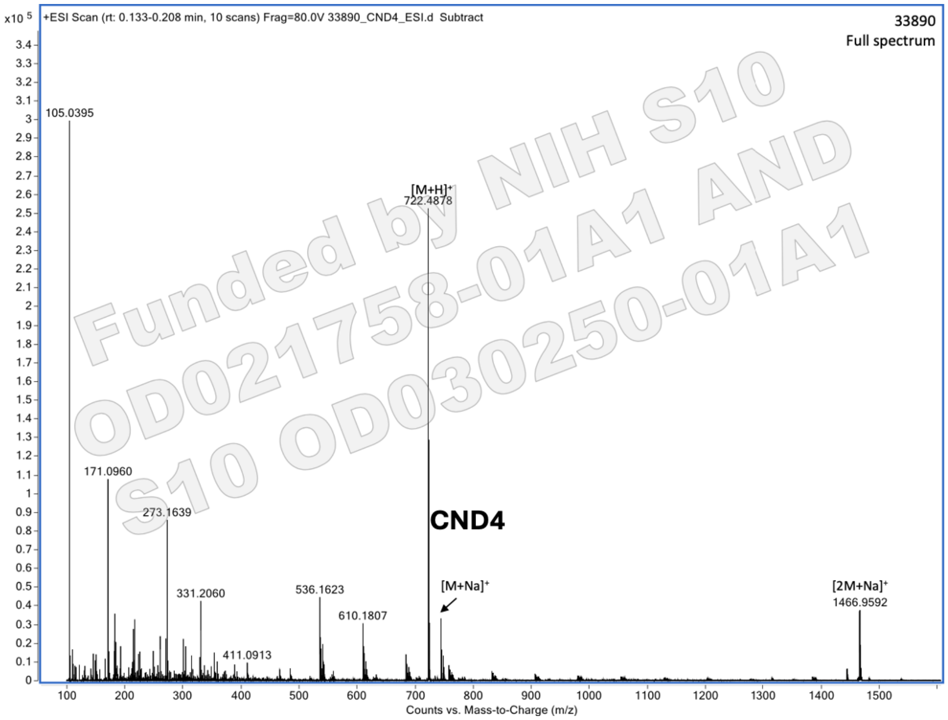
x10 5
+ESI Scan (rt: 0.133-0.208 min, 10 scans) Frag=80.0V 33890_CND4_ESI.d Subtract
33890
Full spectrum
105.0395
[M+H]+
722.4878
171.0960
273.1639
CND4
331.2060
536.1623
610.1807
[M+Na]+
[2M+Na]+
1466.9592
411.0913
Counts vs. Mass-to-Charge (m/z)
Funded by NIH S10 OD021758-01A1 AND S10 OD030250-01A1

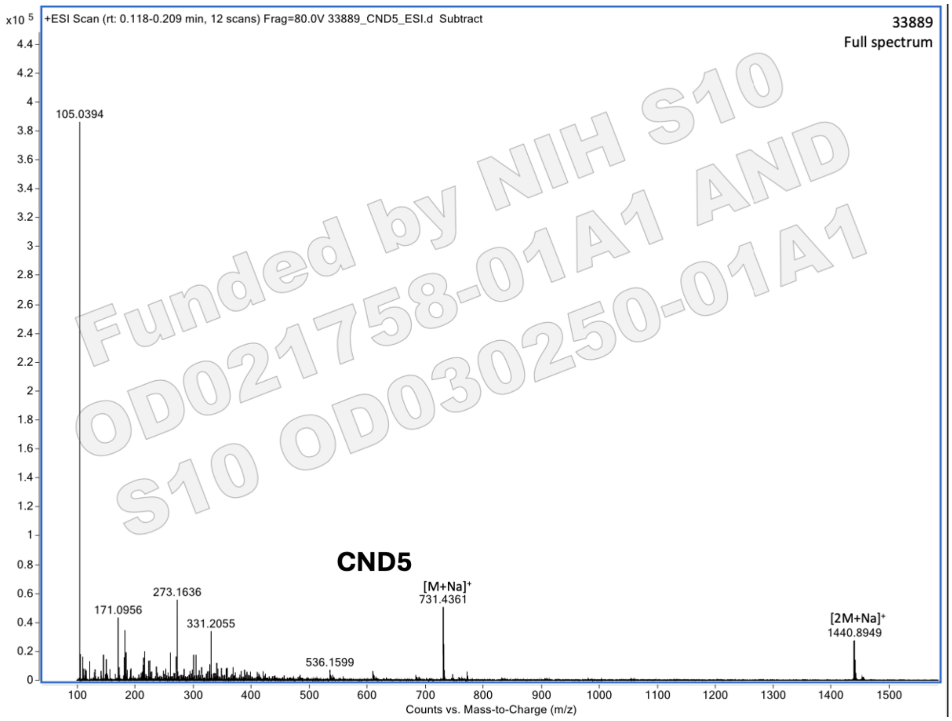
x10 5
+ESI Scan (rt: 0.118-0.209 min, 12 scans) Frag=80.0V 33889_CND5_ESI.d Subtract
33889
Full spectrum
105.0394
CND5
[M+Na]+
731.4361
273.1636
171.0956
331.2055
[2M+Na]+
1440.8949
536.1599
Counts vs. Mass-to-Charge (m/z)
Funded by NIH S10 OD021758-01A1 AND S10 OD030250-01A1

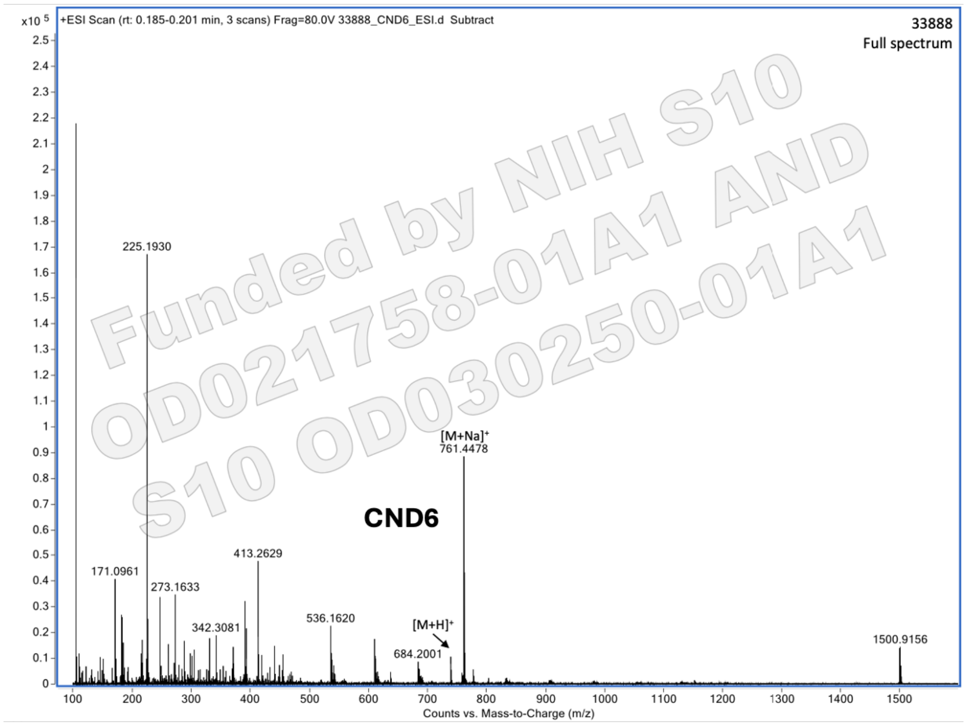
+ESI Scan (rt: 0.185-0.201 min, 3 scans) Frag=80.0V 33888_CND6_ESI.d Subtract
33888
Full spectrum
x10 5
225.1930
171.0961
273.1633
342.3081
413.2629
536.1620
[M+H]+
684.2001
[M+Na]+
761.4478
CND6
1500.9156
Counts vs. Mass-to-Charge (m/z)
Funded by NIH S10 OD021758-01A1 AND S10 OD030250-01A1

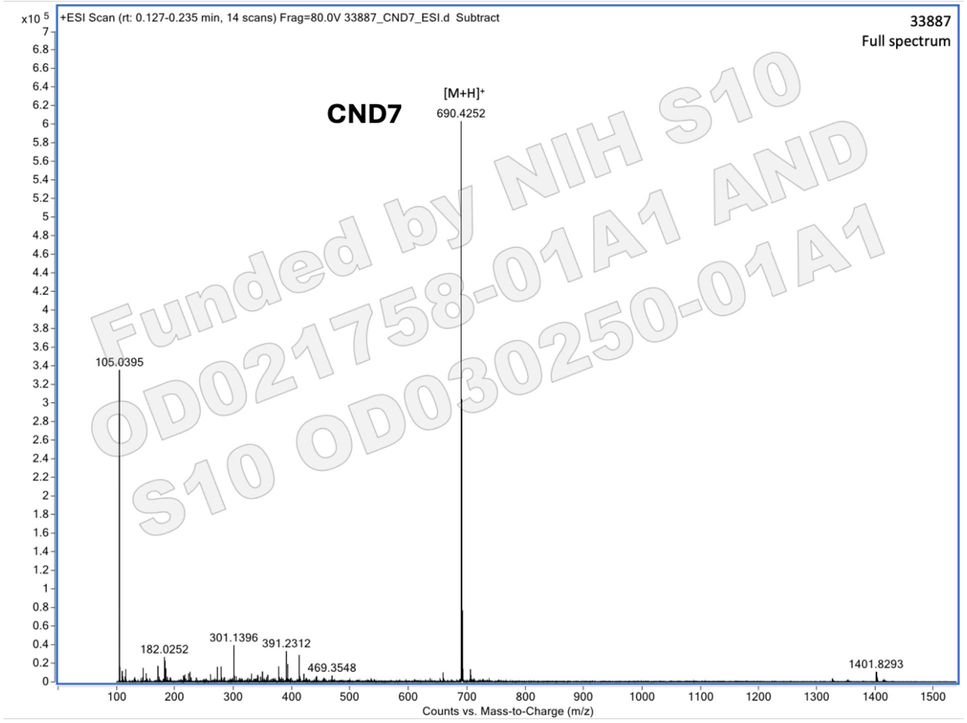
+ESI Scan (rt: 0.127-0.235 min, 14 scans) Frag=80.0V 33887_CND7_ESI.d Subtract
33887
Full spectrum
x10 5
CND7
[M+H]+
690.4252
105.0395
182.0252
301.1396
391.2312
469.3548
1401.8293
Counts vs. Mass-to-Charge (m/z)
Funded by NIH S10 OD021758-01A1 AND S10 OD030250-01A1

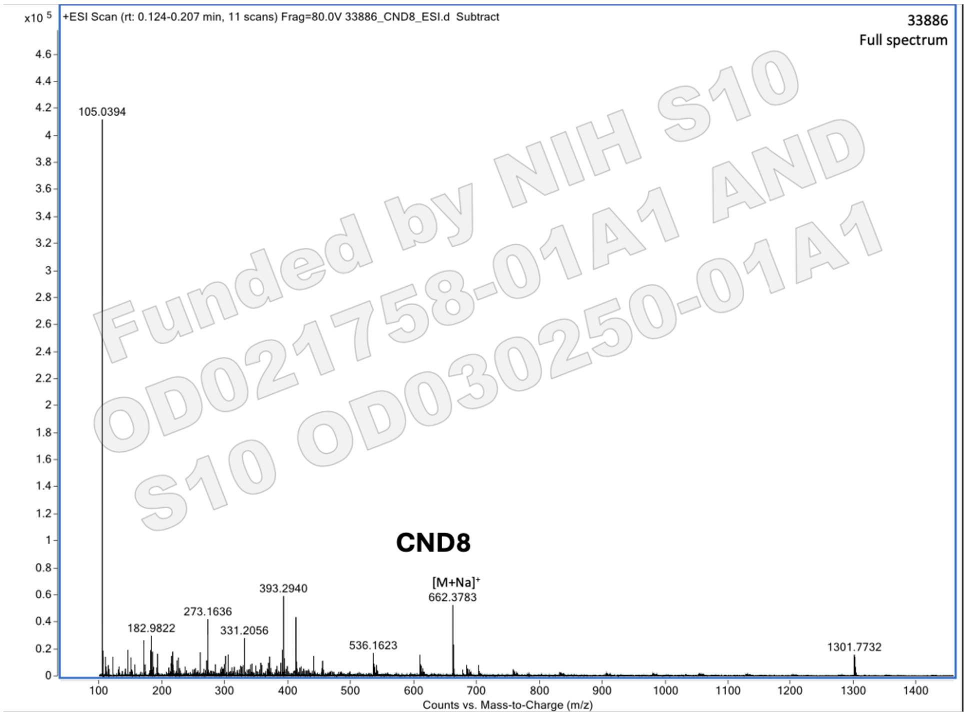


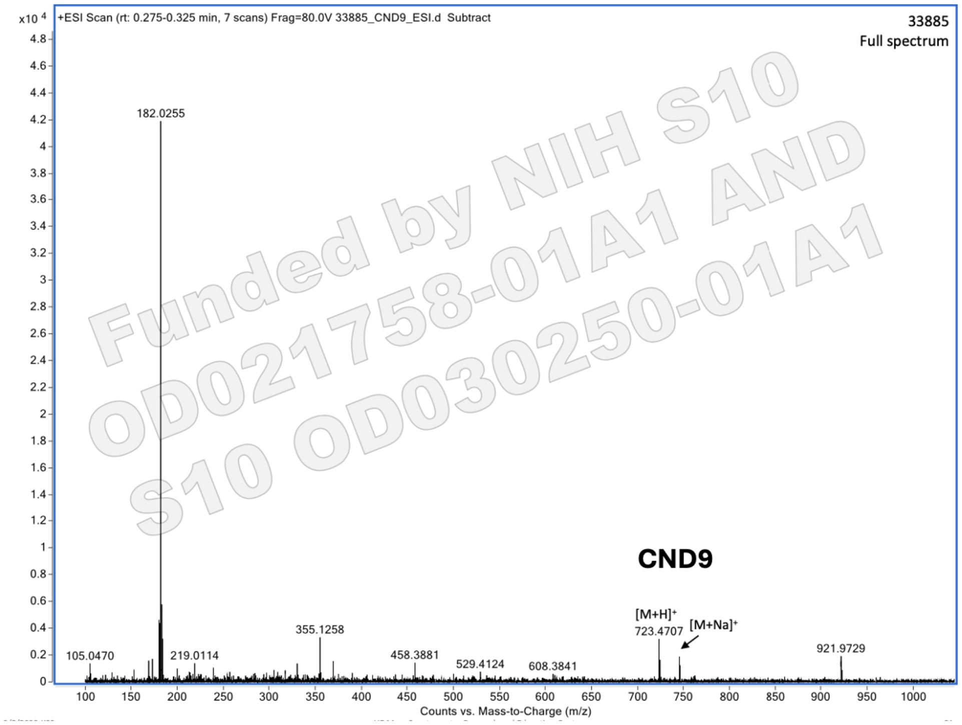


Funded by NIH S10 OD021758-01A1 AND S10 OD030250-01A1

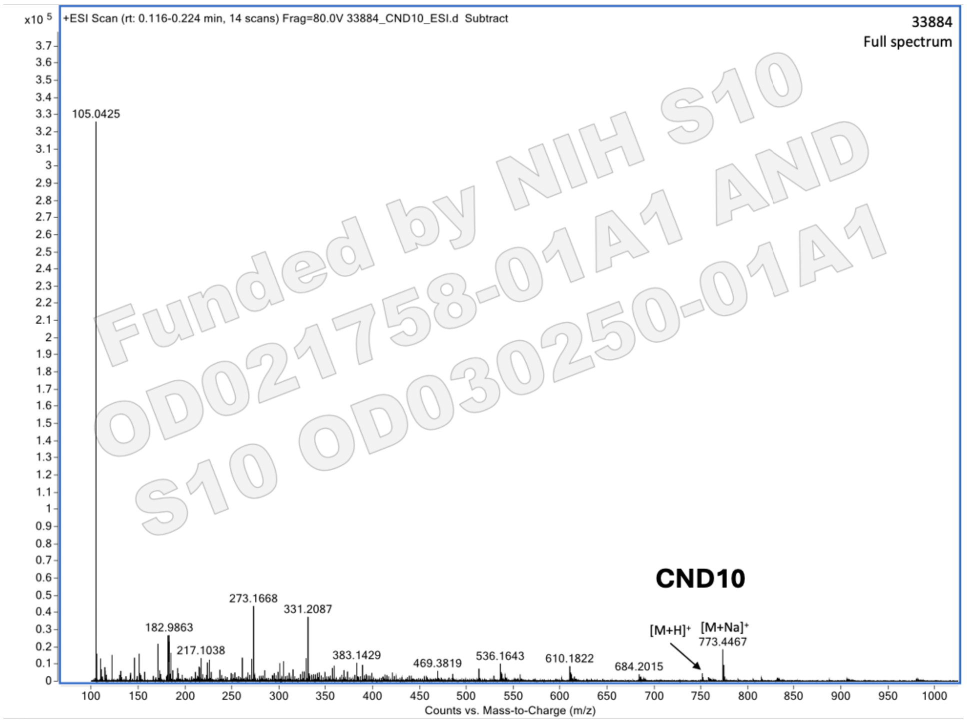

Funded by NIH S10 OD021758-01A1 AND S10 OD030250-01A1

(13) Sych, T.; Schubert, T.; Vauchelles, R.; Madl, J.; Omidvar, R.; Thuenauer, R.; Richert, L.; Mély, Y.; Römer, W. GUV-AP: Multifunctional Fiji-Based Tool for Quantitative Image

Analysis of Giant Unilamellar Vesicles. *Bioinformatics* **2019**, *35* (13), 2340–2342. https://doi.org/10.1093/bioinformatics/bty962.
(14) Kim, S.; Lee, J.; Jo, S.; Brooks, C. L.; Lee, H. S.; Im, W. CHARMM-GUI Ligand Reader and Modeler for CHARMM Force Field Generation of Small Molecules. *J Comput Chem* **2017**, *38* (21), 1879–1886. https://doi.org/10.1002/jcc.24829.
(15) Brooks, B. R.; Brooks, C. L.; Mackerell, A. D.; Nilsson, L.; Petrella, R. J.; Roux, B.; Won, Y.; Archontis, G.; Bartels, C.; Boresch, S.; Caflisch, A.; Caves, L.; Cui, Q.; Dinner, A. R.; Feig, M.; Fischer, S.; Gao, J.; Hodoscek, M.; Im, W.; Kuczera, K.; Lazaridis, T.; Ma, J.; Ovchinnikov, V.; Paci, E.; Pastor, R. W.; Post, C. B.; Pu, J. Z.; Schaefer, M.; Tidor, B.; Venable, R. M.; Woodcock, H. L.; Wu, X.; Yang, W.; York, D. M.; Karplus, M. CHARMM: The Biomolecular Simulation Program. *J Comput Chem* **2009**, *30* (10), 1545–1614. https://doi.org/10.1002/jcc.21287.
(16) Jo, S.; Kim, T.; Iyer, V. G.; Im, W. CHARMM-GUI: A Web-Based Graphical User Interface for CHARMM. *J Comput Chem* **2008**, *29* (11), 1859–1865. https://doi.org/10.1002/jcc.20945.
(17) Vanommeslaeghe, K.; Hatcher, E.; Acharya, C.; Kundu, S.; Zhong, S.; Shim, J.; Darian, E.; Guvench, O.; Lopes, P.; Vorobyov, I.; Mackerell, A. D. CHARMM General Force Field: A Force Field for Drug-like Molecules Compatible with the CHARMM All-atom Additive Biological Force Fields. *J Comput Chem* **2010**, *31* (4), 671–690. https://doi.org/10.1002/jcc.21367.
(18) Wu, E. L.; Cheng, X.; Jo, S.; Rui, H.; Song, K. C.; Dávila-Contreras, E. M.; Qi, Y.; Lee, J.; Monje-Galvan, V.; Venable, R. M.; Klauda, J. B.; Im, W. CHARMM-GUI Membrane Builder toward Realistic Biological Membrane Simulations. *Journal of Computational Chemistry*. John Wiley and Sons Inc. October 15, 2014, pp 1997–2004. https://doi.org/10.1002/jcc.23702.
(19) Abraham, M. J.; Murtola, T.; Schulz, R.; Páll, S.; Smith, J. C.; Hess, B.; Lindah, E. GROMACS: High Performance Molecular Simulations through Multi-Level Parallelism from Laptops to Supercomputers. *SoftwareX* **2015**, *1–2*, 19–25. https://doi.org/10.1016/J.SOFTX.2015.06.001.
(20) Hess, B.; Bekker, H.; Berendsen, H. J. C.; Fraaije, J. G. E. M. *LINCS: A Linear Constraint Solver for Molecular Simulations*; John Wiley & Sons, Inc, 1997; Vol. 18.
(21) Berendsen, H. J. C.; Postma, J. P. M.; van Gunsteren, W. F.; Dinola, A.; Haak, J. R. Molecular Dynamics with Coupling to an External Bath. *J Chem Phys* **1984**, *81* (8), 3684–3690. https://doi.org/10.1063/1.448118.
(22) Humphrey, W.; Dalke, A.; Schulten, K. *VMD: Visual Molecular Dynamics*; 1996.
(23) Guixà-González, R.; Rodriguez-Espigares, I.; Ramírez-Anguita, J. M.; Carrió-Gaspar, P.; Martinez-Seara, H.; Giorgino, T.; Selent, J. MEMBPLUGIN: Studying Membrane Complexity in VMD. *Bioinformatics* **2014**, *30* (10), 1478–1480. https://doi.org/10.1093/bioinformatics/btu037.
(24) Schindelin, J.; Arganda-Carreras, I.; Frise, E.; Kaynig, V.; Longair, M.; Pietzsch, T.; Preibisch, S.; Rueden, C.; Saalfeld, S.; Schmid, B.; Tinevez, J. Y.; White, D. J.; Hartenstein, V.; Eliceiri, K.; Tomancak, P.; Cardona, A. Fiji: An Open-Source Platform for Biological-Image Analysis. *Nat Methods* **2012**, *9* (7), 676–682. https://doi.org/10.1038/nmeth.2019.
(25) Arganda-Carreras, I.; Kaynig, V.; Rueden, C.; Eliceiri, K. W.; Schindelin, J.; Cardona, A.; Seung, H. S. Trainable Weka Segmentation: A Machine Learning Tool for Microscopy Pixel

Classification. *Bioinformatics* **2017**, *33* (15), 2424–2426. https://doi.org/10.1093/bioinformatics/btx180.

(26) Dunn, K. W.; Kamocka, M. M.; McDonald, J. H. A Practical Guide to Evaluating Colocalization in Biological Microscopy. *Am J Physiol Cell Physiol* **2011**, *300* (4), C723–C742. https://doi.org/10.1152/ajpcell.00462.2010.

(27) Costes, S. V.; Daelemans, D.; Cho, E. H.; Dobbin, Z.; Pavlakis, G.; Lockett, S. Automatic and Quantitative Measurement of Protein-Protein Colocalization in Live Cells. *Biophys J* **2004**, *86* (6), 3993–4003. https://doi.org/10.1529/biophysj.103.038422.

(28) Stauffer, W.; Sheng, H.; Lim, H. N. EzColocalization: An ImageJ Plugin for Visualizing and Measuring Colocalization in Cells and Organisms. *Sci Rep* **2018**, *8* (1), 15764. https://doi.org/10.1038/s41598-018-33592-8.
